# RNA-mediated Genomic Arrangements in Mammalian Cells

**DOI:** 10.1101/151241

**Authors:** Sachin Kumar Gupta, Liming Luo, Laising Yen

## Abstract

One of the hallmarks of cancer is the formation of oncogenic fusion genes as a result of chromosomal translocations. Fusion genes are presumed to occur prior to fusion RNA expression. However, studies have reported the presence of fusion RNAs in individuals who were negative for chromosomal translocations. These observations give rise to “the cart before the horse” hypothesis, in which fusion RNA precedes the fusion gene and guides the genomic rearrangements that ultimately result in gene fusions. Yet RNA-mediated genomic rearrangement in mammalian cells has never been demonstrated. Here we provide evidence that expression of a chimeric RNA drives formation of a specified gene fusion via genomic rearrangement in mammalian cells. The process is (1) specified by the sequence of chimeric RNA involved, (2) facilitated by physiological hormone levels, (3) permissible regardless of intra-chromosomal (*TMPRSS2-ERG*) or inter-chromosomal (*TMPRSS2-ETV1*) fusion, and (4) can occur in normal cells prior to malignant transformation. We demonstrate that, contrary to “the cart before the horse” model, it is the antisense rather than sense chimeric RNAs that effectively drive gene fusion, and that this disparity can be explained by transcriptional conflict. Furthermore, we identified an endogenous RNA *AZI1* that acts as the ‘initiator’ RNA to induce *TMPRSS2-ERG* fusion. RNA-driven gene fusion demonstrated in this report provides important insight in early disease mechanism, and could have fundamental implications in the biology of mammalian genome stability, as well as gene editing technology via mechanisms native to mammalian cells.

## Introduction

Fusion genes are among the most cancer-specific molecular signatures known. They are important for understanding cancer mechanisms and developing useful clinical biomarkers and anti-cancer therapies (Mitelman et al., 2007). Fusion gene formation as a result of chromosomal translocations is presumed to occur prior to fusion RNA expression. However, several studies have reported the presence of fusion transcripts in individuals without detectable fusion genes at the genomic DNA level. For instance, the *AML1-ETO* fusion transcript, associated with a subtype of acute myeloid leukemia, was present in some patients who were negative for chromosomal translocations (Langabeer et al., 1997). Other fusion RNAs, such as *BCR-ABL*, *MLL-AF4*, *TEL-AML1*, *PML-RARα*, and *NPM-ALK*, were reported in healthy individuals (Janz et al., 2003). Although the discrepancy between the presence of fusion transcripts and the absence of fusion genes could result from detection limitations of the methodologies employed, fusion transcripts in normal cells could also arise from RNA trans-splicing in the absence of chromosomal translocations (Zaphiropoulos, 2011). Indeed, *JAZF1-JJAZ1* fusion transcripts are expressed in normal human endometrial tissue and an endometrial cell line in the absence of chromosomal translocation (Li et al., 2008). Furthermore, trans-splicing between *JAZF1* and *JJAZ1* was demonstrated to occur *in vitro* using cellular extracts, resulting in a fusion RNA similar to that transcribed from the *JAZF1-JJAZ1* fusion gene in endometrial stromal sarcomas (Li et al., 2008). These observations raise the possibility that cellular fusion RNAs created by trans-splicing act as guide RNAs to mediate genomic rearrangements. A precedent for RNA-mediated genomic arrangements is found in lower organisms such as ciliates (Fang and Landweber, 2012; Nowacki et al., 2008). Rowley and Blumenthal (Rowley and Blumenthal, 2008) coined this as “the cart before the horse” hypothesis, in that “RNA before DNA” defies the normal order of the central dogma of biology: DNA → RNA → protein (Crick, 1970). Despite important implications in biology and human cancer, RNA-mediated genomic rearrangement in mammalian cells has not been directly demonstrated. In this report, we provide the first evidence that expression of a specific chimeric RNA can lead to specified gene fusion in mammalian cells.

## Results

To test whether the expression of a fusion RNA in mammalian cells can lead to a specific gene fusion, the *TMPRSS2-ERG* fusion (Perner et al., 2006; Tomlins et al., 2005), found in ~50% of prostate cancers, was selected as a model. Both the *TMPRSS2* and *ERG* genes are located on chromosome 21, an intra-chromosomal configuration prone to rearrangements. To recapitulate *TMPRSS2-ERG* fusion gene formation, we used the LNCaP prostate cancer cell line that lacks the *TMPRSS2-ERG* fusion (Horoszewicz et al., 1980; Tomlins et al., 2005). Furthermore, treating LNCaP cells with androgen increases the chromosomal proximity between the *TMPRSS2* and *ERG* genes (Bastus et al., 2010; Lin et al., 2009; Mani et al., 2009), which was thought to increase the possibility of gene fusion. To test “the cart before the horse” hypothesis (Rowley and Blumenthal, 2008; Zaphiropoulos, 2011), we transiently expressed a short fusion RNA consisting of two exons, *TMPRSS2* exon-1 joined to *ERG* exon-4, which is a short fragment of a full-length *TMPRSS2-ERG* fusion RNA that is most common in prostate cancer (Fig. 1A, upper panel). This short fusion RNA mirrors the presumptive trans-spliced fusion RNA product that is generated only in the sense orientation because the correct splice sites are absent in the antisense orientation. However, because the ‘antisense’ sequence should, in theory, contain the same template information for guiding genomic rearrangements, we tested both the sense and antisense short fusion RNA. Each was individually expressed using either a CMV or a U6 promoter (Fig. 1A, upper panel) and designated as ‘input RNA’ to distinguish them from the ‘endogenous’ full-length fusion RNA transcribed from the genome.

**Fig 1.**
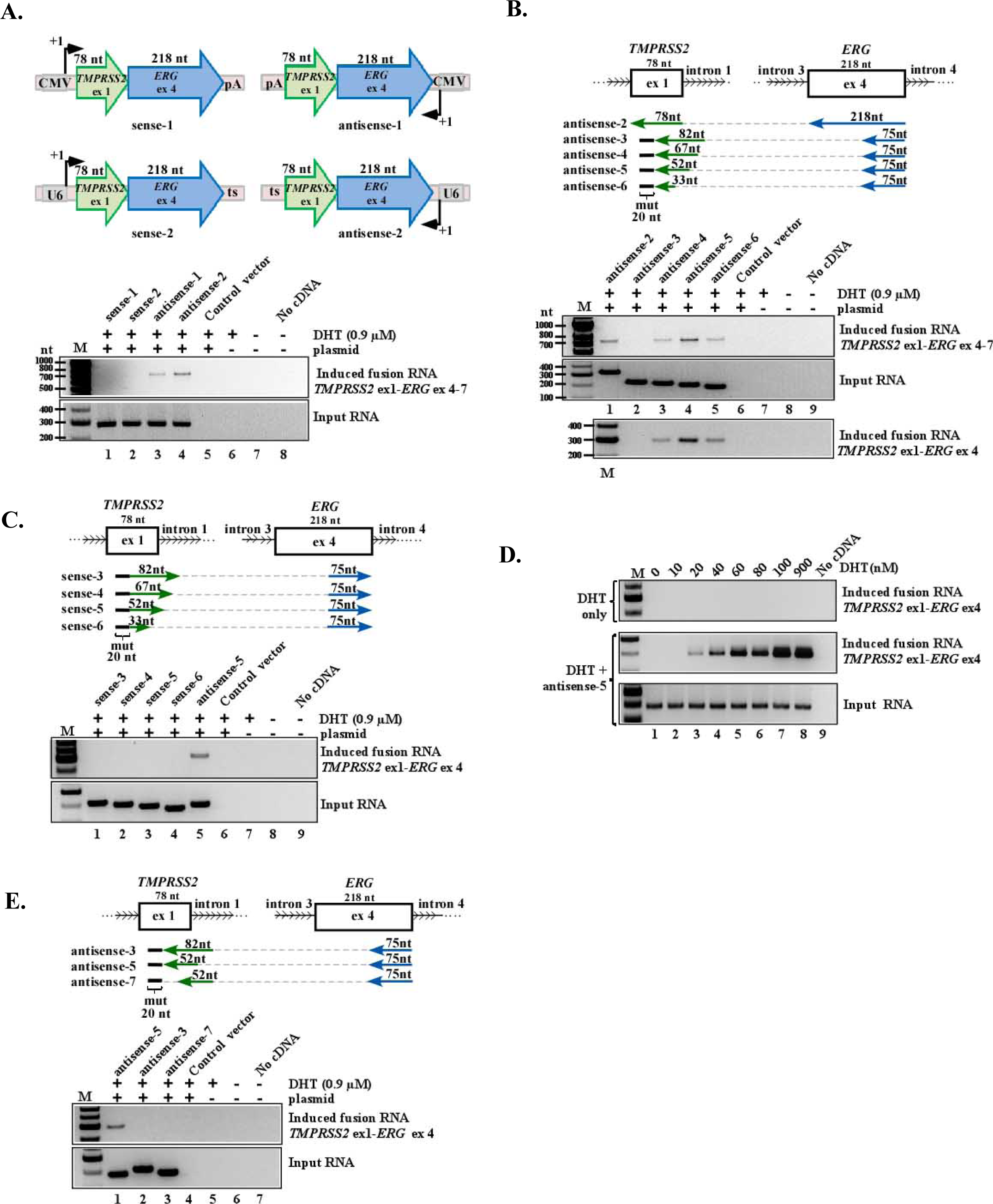
Exogenously expressed input chimeric RNAs induce the expression of endogenous fusion transcripts. **A. Upper**: Schematics of the designed input RNAs containing complete *TMPRSS2* exon-1 (78 nt, uc002yzj.3) and *ERG* exon-4 (218 nt, uc021wjd.1), expressed in the sense or antisense orientation from the CMV or U6 promoters. pA: poly-A signal. ts: transcriptional stop “TTTTTT” for U6 promoter. **Lower**: RT-PCR detection of induced fusion transcript (top gel) and input RNA (bottom gel). Parental plasmid vector containing mCherry sequence, DHT treatment without plasmid transfection, and PCR reaction without cDNA served as the controls (land 5, 6, 8). **B.** Length and positional effect of antisense input RNAs. **Upper**: Antisense input RNAs with 75 nt (blue) targeting *ERG* exon-4 and varying lengths (82, 67, 52, and 33 nt, green) targeting *TMPRSS2.* Dashed line links *ERG* and *TMPRSS2* sequence within the input RNA and contains no sequence. A 20-nt mutation (black line) was introduced to the input RNAs to discern expressed input RNAs from the induced fusion transcript (see Fig. S1B). **Lower**: RT-PCR detection of induced fusion transcript (top gel), input RNA (middle gel), or detection of induced fusion transcript using a different primer pair (bottom gel). **C.** Corresponding sense input RNAs all failed to induce the fusion transcript. **D.** Induction by antisense-5 occurred at physiologically relevant DHT concentrations as low as 20 nM. Three-rounds of nested PCR were performed to reveal the lowest amount of DHT required. **E**. Antisense-5 led to clear induction while antisense-3 and 7 did not, indicating that it is not the length of input RNAs but targeted regions that is critical.

We transiently transfected LNCaP cells with either plasmid and treated the cells with dihydrotestosterone (DHT, a metabolite of testosterone) for 3 days. If the expression of an input RNA leads to a *TMPRSS2-ERG* gene fusion, it is expected that the endogenous full-length fusion RNAs would be transcribed from the newly induced fusion gene. Specific RT-PCR assays were designed to distingwish between endogenous full-length fusion RNAs and the input RNAs exogenously expressed from the plasmids (see Fig. S1A for primer designs). As shown in Fig. 1A, expression of the sense short fusion RNA resembling the trans-spliced product, either by the CMV or U6 promoter (Fig. 1A lower panel, lane 1 and 2 respectively), led to no detection of an induced endogenous fusion transcript. Expression of a longer version of sense fusion RNA consisting of four exons (*TMPRSS2* exon-1 joined to *ERG* exon-4/5/6) also failed to induce the endogenous fusion transcript (Fig. S2). In contrast, expression of antisense short fusion RNAs induced a clear band of 721 bp (Fig. 1A lower panel, lane 3 and 4). Sanger sequencing revealed that the induced band contains *TMPRSS2* exon-1 fused to *ERG* exons-4/5/6/7 (Fig. S3), and that the exons are joined by annotated splice sites, which would be expected of mature endogenous fusion mRNA derived from the *TMPRSS2-ERG* fusion gene. This induced fusion transcripts cannot possibly arise from the sequence of input RNAs as the expression plasmids used contain only *TMPRSS2* exon-1 and *ERG* exon-4 without the *ERG* exon-5/6/7 sequence. Notably, the induction was more pronounced when the antisense input RNA was driven by the U6 promoter (Fig. 1A lower panel, antisense-2, lane 4) compared to the CMV promoter (antisense-1, lane 3). These differences (antisense vs. sense, U6 vs. CMV) are not caused by differing amounts of input RNA because all input RNAs were expressed at relatively equal levels (Fig. 1A, lower panel). Transfection with a parental plasmid containing mCherry sequence (Fig. 1A, lane 5), DHT treatment without plasmid transfection (Fig. 1A, lane 6), and PCR reaction without cDNA served as RT-PCR controls (Fig. 1A, lane 8), all resulted in no endogenous fusion transcript. In addition, all experiments were performed independently at least four times and results were identical. Together, the data suggest that expression of an input RNA with chimeric sequence can lead to the induction of a specified endogenous fusion transcript in human cells. Surprisingly, the antisense, rather than the sense version of input RNA, exhibits the capacity of induction.

Antisense input RNAs described above contain 218 nt against the entire *ERG* exon-4 and 78 nt against the entire *TMPRSS2* exon-1 (Fig. 1A), suggesting that 78 nt is sufficient to specify a parental gene for gene fusion. Furthermore, because the effective input RNAs are of the ‘antisense’ orientation, the data imply that the input RNAs may not require an RNA junction resembling that of the *TMPRSS2-ERG* fusion transcript generated by splicing in the sense orientation. To further analyze the sequence requirement, we used the U6 promoter to express a series of antisense input RNAs with 75 nt complementary to *ERG* exon-4 joined to various segments (33, 52, 67, 82 nt) that are complementary to *TMPRSS2* near the exon-1/intron-1 boundary (Fig. 1B). A parallel set of sense input RNAs were also tested as controls (Fig. 1C). As shown in Fig. 1B (lower panel), all antisense RNAs, with the exception of antisense-3, induced fusion transcripts even though their target regions span the exon/intron boundary. The level of induction peaked for antisense-5, which contains 52 nt designed to anneal with *TMPRSS2*, suggesting that this number of nucleotides might be optimal to specify a parental gene for induction. The results were confirmed using a different, but more efficient, primer pair (Fig. 1B, lower panel; primer design in Fig. S1B) followed by Sanger sequencing of the induced band (Fig. S4). In contrast to the antisense input RNA, all corresponding sense input RNAs failed to induce endogenous fusion transcripts (Fig. 1C, lower panel). This was true even when the sense input RNA was intentionally expressed at a much higher level than the antisense RNA (Fig. S5). Additional experiments using plasmids with a severed U6 promoter (Fig. S6), to eliminate input RNA expression, also confirmed that it is the antisense input RNAs expressed from plasmids, not the DNA sequence of plasmids, that induce the observed *TMPRSS2-ERG* fusion transcripts.

As shown in Fig. 1D, the amount of endogenous fusion transcript induced by antisense-5 (the most effective antisense input RNA) appears to correlate with the concentration of DHT used, presumably because the hormone increases the chromosomal proximity between the *TMPRSS2* and *ERG* genes (Bastus et al., 2010; Lin et al., 2009; Mani et al., 2009). Antisense-5 was effective at DHT concentrations as low as 20 nM as revealed by sensitive nested PCR (Fig. 1D lane 3), indicating that fusion events induced by input RNA can occur under physiologically relevant androgen conditions (Boyce et al., 2004). As a control, DHT treatment alone up to 2 μM failed to induce fusion (Fig. S7). Titration of DHT showed that the induction by antisense-5 reaches the 50% of maximal level (EC_50_) at 0.9 μM DHT (Fig. S7). Under this standard EC_50_ condition, we estimated that the percentage of LNCaP cells induced by antisense-5 to express the *TMPRSS2-ERG* fusion transcript is approximately 1 in 10^3^ or 10^4^ cells (see assay in Fig. S8). Together, these results demonstrated that the induction by input RNA can occur at physiologically relevant hormone levels, but does not represent a high frequency event.

Although induced fusion is infrequent, all antisense RNAs described in Fig. 1B successfully induced endogenous fusion RNA except antisense-3, which is only 30 nt longer than antisense-5 in the arm targeting *TMPRSS2* intron-1 (Fig. 1B, upper panel). To test whether its inability to induce was due to input RNA length or the specific target sequence in *TMPRSS2* intron-1, we constructed a hybrid antisense (antisense-7) that shifted the 52 nt recognition window of antisense-5 to target the *TMPRSS2* intron-1 region covered by antisense-3 (Fig. 1E). This alteration resulted in the loss of induction (Fig. 1E, lane 1 vs. 3), implying that the inability of antisense-3 to induce is not reflective of input RNA length. Rather its targeting arm may interfere with a motif important for the fusion process. BLAST alignment of the genomic DNA sequence revealed an imperfect stem (named stem A) potentially formed by the sense genomic *TMPRSS2* sequence complementary to the sense genomic *ERG* sequence (Fig. 2A, left). We reasoned that this genomic DNA stem (Tm= ~44°C) could potentially stabilize a three-way junction that involves an RNA/DNA duplex formed by the antisense-5 RNA and its targeted genomic DNA in a sequence-specific manner (Fig. 2A, left). If correct, then the formation of this putative three-way junction would be disrupted by antisense-3 because its recognition sequence invades the genomic DNA stem (Fig. 2A, left). Consistent with the idea that induction requires bringing *TMPRSS2* and *ERG* gene in close proximity, expression of antisense-5 as two separate halves (Fig. 2A right panel, antisense-5A and -5B) severed the link between *TMPRSS2* (52 nt) and *ERG* (75 nt) sequence within the input RNA, resulting in the loss of induction (Fig. 2B, lane 1 to 3).

**Fig 2.**
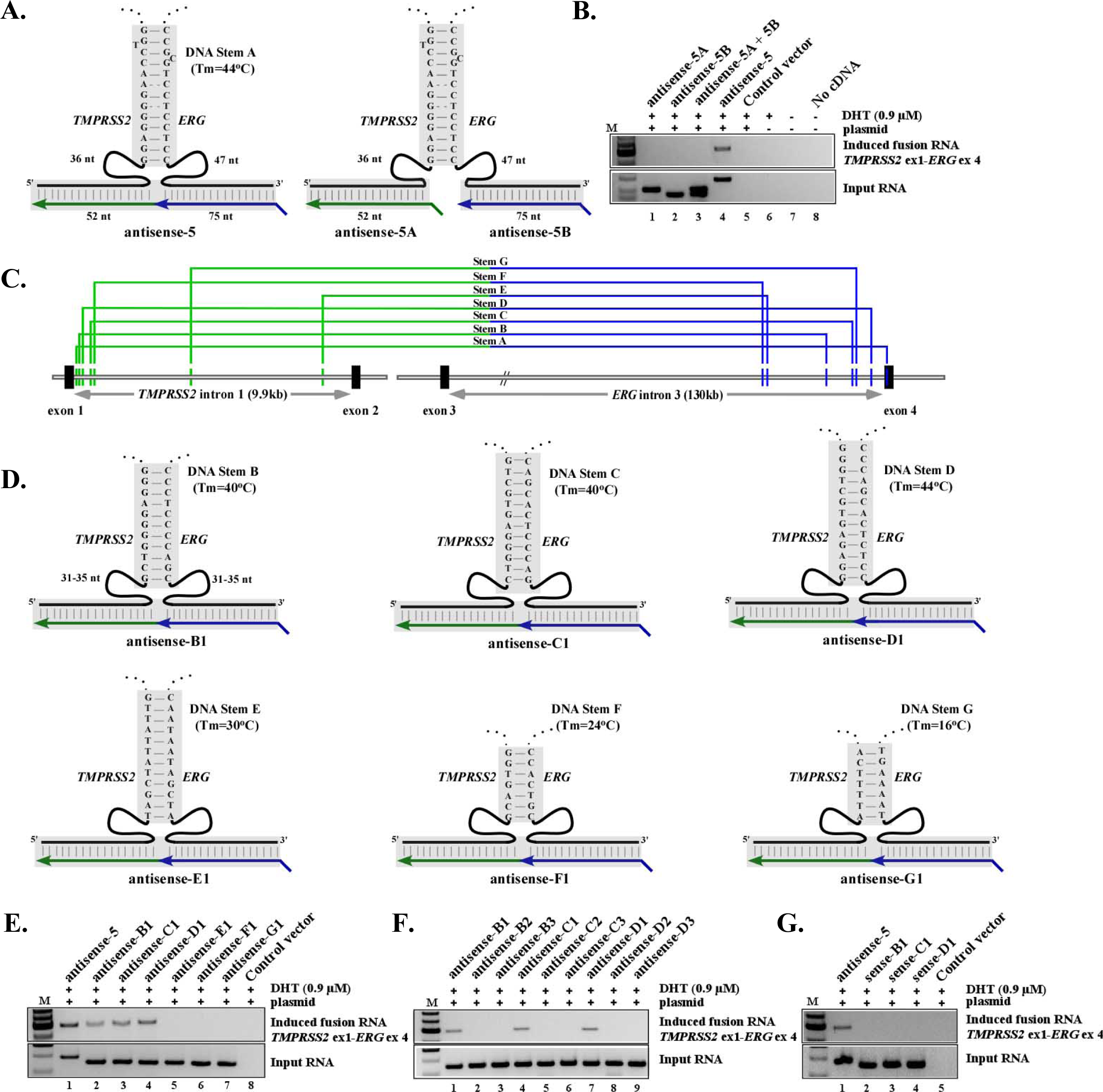
Formation of a three-way junction may facilitate fusion induction. **A. Left:** Schematics of three-way junction that could be formed between genomic DNA (black) and antisense-5 input RNA (green/blue). The sense genomic strands of both *TMPRSS2* and *ERG* genes are on the minus strand of chromosome 21, separated by 3 Mb. Short lines in shaded regions represent base-pairings. Imperfect stem A includes a high energy G·T and A·C wobble pair known to have Watson-Crick-like geometry in a DNA double helix (Kimsey and Al-Hashimi, 2014; Watson and Crick, 1953). A spacer region of 36 nt and 47 nt separate stem A from the regions targeted by antisense-5 input RNA. **Right:** Expressing antisense-5 as two separate halves that severed the link between *TMPRSS2* (52 nt) and *ERG* (75 nt) sequence within the input RNA. **B.** RT-PCR assays of fusion transcripts showed that the severed input RNAs resulted in the loss of induction. **C.** Locations of putative stems A to F identified by BLAST analyses. Genomic coordinates are listed in Fig. S9. **D.** The putative three-way junction formed between the indicated genomic DNA stem B to G (black) and designed antisense input RNA (green/blue). **E.** Targeting genomic DNA stem B, C, and D that exhibit higher DNA stem stability (Tm= 40°C, 40°C, and 44°C, respectively) by the corresponding antisense input RNAs induced fusion transcripts (lanes 2 to 4). In contrast, targeting less stable stem E, F, and G (Tm= 30°C, 24°C, and 16°C, respectively) failed to induce fusion transcripts (lanes 5 to 7). **F.** Antisense input RNAs designed to invade each of the respective genomic DNA stem B, C, and D resulted in the loss of induction. **G.** Corresponding sense input RNAs targeting stem B, C, and D failed to induce fusion transcripts (lanes 2 to 4).

To test whether the proposed three-way junction formation could facilitate induction, we used BLAST alignment to identify several intron locations where genomic DNA stems could be formed by the sense genomic *TMPRSS2* sequence paired with the sense genomic *ERG* sequence (stems B to G in Fig. 2C and 2D; genomic coordinates in Fig. S9; sequences flanking the stems in Fig. S10). Matching antisense input RNAs (termed antisense-B1 to G1) were then designed to facilitate the formation of a three-way junction with the possible intron stems (Fig. 2D) that would mirror the three-way junction formed by antisense-5 on stem A as postulated in Fig 2A. Because these input RNAs target the introns (Fig. 2C) and contain no exon sequence, any observed induction of endogenous fusion transcripts composed of exons cannot arise from the sequence of input RNAs or plasmids used to express them. As shown in Fig. 2E, targeting genomic DNA stem B, C, and D that exhibit higher DNA stem stability (Tm= 40°C, 40°C, and 44°C, respectively) by the corresponding antisense input RNAs clearly induced fusion transcripts (Fig. 2E, lanes 2 to 4). In contrast, targeting less a stable stem E, F, and G (Tm= 30°C, 24°C, and 16°C, respectively) failed to induce fusion transcripts (Fig. 2E, lanes 5 to 7). To disrupt the three-way junction invovling stem B, C, and D, six additional antisense RNAs (antisense-B2, B3, C2, C3, D2, and D3) were designed with one side of their recognition sequence altered to invade each of the respective genomic DNA stems on the *TMPRSS2* side or the *ERG* side (Fig. S11). These modifications were chosen to mirror the interference on stem A by antisense-3. Similar antisense RNAs were also designed to invade stem A (Fig. S12). In all cases, invasion of the genomic DNA stems by the modified input RNAs resulted in the complete loss of induction (Fig. 2F). While these results by no means necessitate that a three-way junction is required for fusion transcript induction, they nevertheless suggest that such transiently stabilized structures may ‘facilitate’ the process and could have important implications in developing gene editing technologies via mechanisms native to mammalian cells. Consistent with earlier observations, the corresponding sense version of the effective antisense input RNAs (sense-B1, C1, D1) all failed to induce fusion transcripts (Fig. 2G, lanes 2 to 4).

The fact that antisense input RNAs, but not their sense counterparts, induce fusion transcripts, raises the possibility that the former act as a docking station to mediate trans-splicing between endogenous sense *TMPRSS2* and *ERG* pre-mRNAs. Because that the antisense, but not the sense input RNAs, are complimentary to both sense *TMPRSS2* and *ERG* pre-mRNAs, they can base-pare with both parental pre-mRNAs, thus resulting in spliced fusion transcripts without the requirement of genomic rearrangement. This mechanism, however, is unlikely as the major contributor to the observed induction for the following reasons. First, although *TMPRSS2* is expressed in LNCaP cells (Fig. 3A, upper panel), endogenous *ERG* mRNA is not detected in LNCaP cells (Tomlins et al., 2005) in the presence or absence of DHT or before and after transfection of antisense-5 (Fig. 3A, middle and lower panel with different primer pairs). In fact, parental *ERG* mRNA was not detected even using three rounds of nested RT-PCR using various primer sets (Fig. S13). Therefore prior to and during induction, no or an insufficient number of parental *ERG* mRNAs are available in LNCaP cells as raw material for trans-splicing to account for the level of induced fusion transcript. Second, after initial transient transfection and DHT treatment for 3 days, we continued to propagate and enrich the induced LNCaP population for 52 days in the absence of DHT (experimental procedures described in Fig. S14). As shown in the lower panel of Fig. 3B, antisense-5 RNA transiently expressed by plasmids was degraded and completely absent beyond day 17. In contrast, the induced fusion transcript was continuously expressed and enriched up to day 52 in the absence of antisense input RNA and DHT (Fig. 3B upper panel), indicating the persistent nature of the induced fusion product. Taken together, these results strongly suggest that the induced expression of the *TMPRSS2-ERG* fusion transcript is the consequence of gene fusion at the DNA level, which has a permanent nature. This is in contrast to the result of induced trans-splicing at the RNA level mediated by antisense input RNA, which is transient and requires the continuous presence of input RNAs.

To provide definite evidence of gene fusion via genomic rearrangement, we used genomic PCR to identify the genomic breakpoint induced by antisense-5 in the enriched LNCaP population (primer designs in Fig. S15A and Fig. 3C). As shown in Fig. 3D, the un-rearranged wildtype *TMPRSS2* and *ERG* alleles were amplified by gene-specific primer pair A/B and C/D both in untransfected cells (Fig. 3D, lane 1 and 2) and enriched LNCaP cells (Fig. 3D, lane 4 and 5). In contrast, a genomic fusion band of ~862 bp amplified by fusion-specific primer pair A/D was present only in the enriched LNCaP population (Fig. 3D, lane 6) and absent in untransfected LNCaP cells (lane 3). Sanger sequencing of the excised fusion band (Fig. 3D, lane 6) revealed the exact genomic breakpoint located within *TMPRSS2* intron-1 (chr21:41502038, GRCh38/hg38) and *ERG* intron-3 (chr21:38501207, GRCh38/hg38) (Fig. 3E; full-length Sanger sequence shown in S16). Intriguingly, within *TMPRSS2* intron-1 the induced breakpoint lies within an Alu, a transposable element known to contribute to genomic arrangements (Rudiger et al., 1995). In *ERG* intron-3, the breakpoint resides in a hot spot clustered with genomic breakpoints previously identified in prostate cancer patients (Fig. S15B) (Weier et al., 2013). There is no obvious sequence homology between *TMPRSS2* and *ERG* at the genomic breakpoint except for a three nucleotide ‘CTG’ microhomology (Fig 3E and S16), suggesting that this gene fusion may be mediated by non-homologous break repair mechanisms (Lieber, 2010; Zhang et al., 2009).

To test whether antisense input RNA can cause *TMPRSS2-ERG* fusion in non-malignant cells prior to cancerous transformation, we performed experiments using immortalized normal prostate epithelium cells (PNT1A), that express very low levels of androgen receptors (Coll-Bastus et al., 2015). As shown in the lower panel of Fig. 3F, prolonged expression of antisense-5 for 12 days induced fusion transcripts (Sanger sequencing confirmation in Fig. S17). This induction was not due to prolonged exposure to DHT because continuous treatment of 0.9 μM DHT alone for up to 2 months resulted in no detectable fusion transcripts in PNT1A cells (Fig. 3F, lane 8). Thus, our results indicate that the induction of *TMPRSS2-ERG* fusion by antisense input RNA can occur in normal prostate epithelial cells prior to malignant transformation and is not restricted to the pathological cellular context of malignant cells.

**Fig 3.**
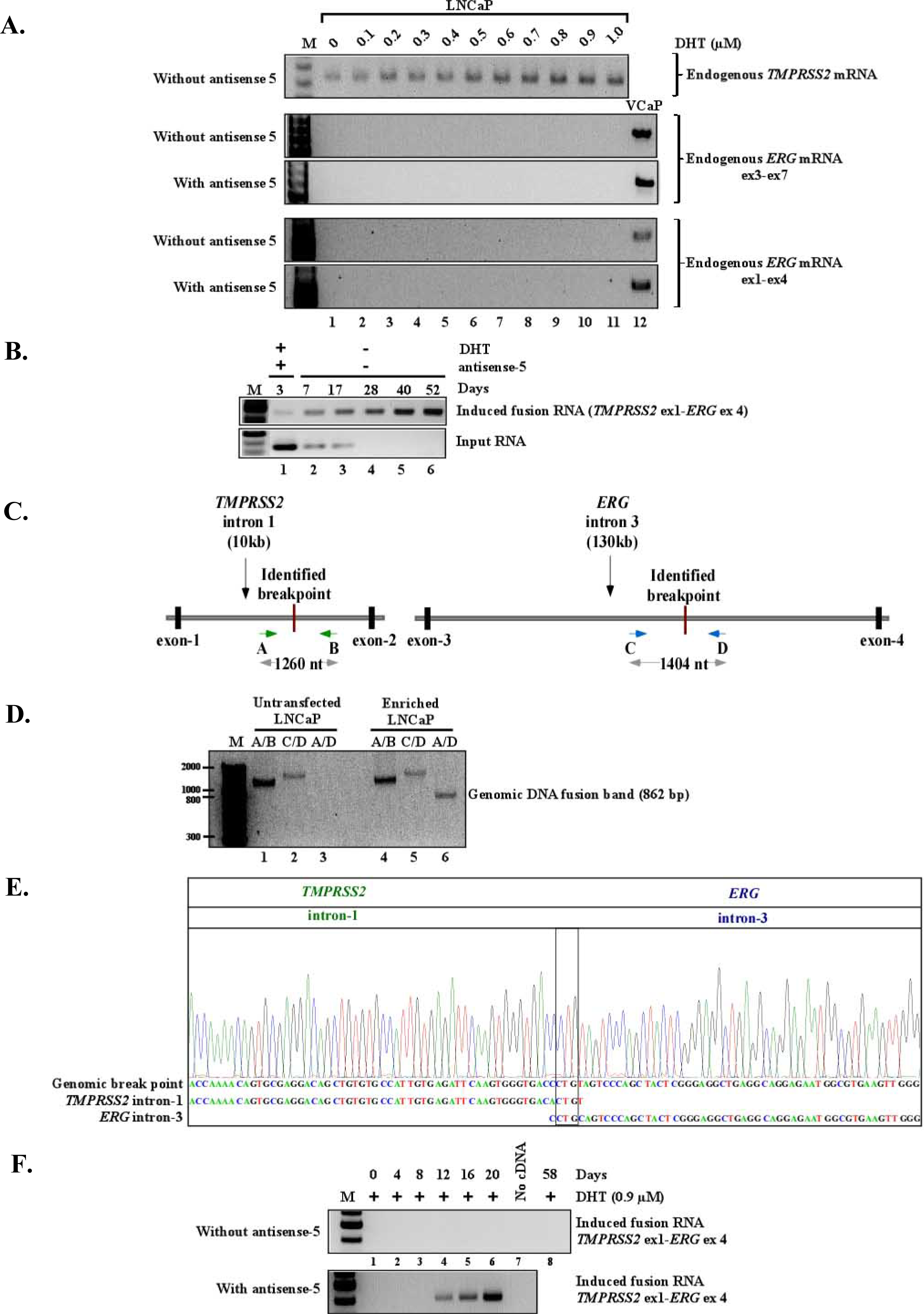
Induced *TMPRSS2-ERG* fusion is the result of genomic arrangements. **A. Upper:** RT-PCR shows that LNCaP cells express *TMPRSS2* mRNA, and that the expression is upregulated by DHT. Primers used are specific to *TMPRSS2* exon-2 and exon-4. **Middle and lower:** *ERG* mRNA however was not detected in LNCaP cells under a wide range of DHT in the presence or absence of antisense-5 (lanes 1 to 11). Assays were performed using two independent primer pairs, one specific to exon-3 and -7 (middle panel), the other to exon-1 and - 4 (lower panel), both capable of detecting *ERG* mRNA in VCaP cells (lane 12). These primer pairs were chosen because they selectively amplified endogenous *ERG* mRNA but not the induced *TMPRSS2-ERG* fusion transcript which has *ERG* exon-3 to exon-12. **B.** RT-PCR shows the transient nature of input RNA (lower panel) and the persistent nature of the induced fusion transcript (upper panel) in the enriched LNCaP population up to 52 days post initial treatment (see Fig. S14 for enrichment procedure). **C.** Schematics of identified genomic breakpoints and the primers used to amplify the breakpoints. The approximate location of genomic breakpoints were first determined using a primer set targeting *TMPRSS2* intron-1 multiplexed with a primer set targeting *ERG* intron-3 (see Fig. S15A). The region near the breakpoint was then further examined using primers A, B, C, and D. **D.** The un-rearranged wildtype *TMPRSS2* and *ERG* alleles were revealed by primer pair A/B (~1404 bp) and C/D (~1260 bp) respectively (lane 1, 2, 4, 5). The genomic fusion band of 862 bp amplified by fusion-specific primer pair A/D was present only in the enriched LNCaP population (lane 6) and absent in untransfected LNCaP cells (lane 3). **E.** Sanger sequencing of the fusion band showed a 500bp segment of *TMPRSS2* intron-1 fused to 362 bp of *ERG* intron-3. The genomic breakpoint contains a ‘CTG’ microhomology (boxed). The full-length Sanger sequence is shown in S16. **F.** Prolonged expression of antisense-5 for 12 days induced the *TMPRSS2-ERG* fusion transcript in PNT1A cells as detected by three-round nested PCR. The results indicate that the induction of *TMPRSS2-ERG* fusion by antisense input RNA can occur in normal prostate epithelial cells prior to malignant transformation.

To test whether an input RNA can specify a pair of genes to undergo fusion other than *TMPRSS2-ERG* in a sequence-specific manner, we designed a series of input RNAs to induce *TMPRSS2-ETV1*, an inter-chromosomal fusion gene found in approximately 1% of prostate cancers (Rubin et al., 2011; Tomlins et al., 2005). Eight antisense RNAs (Fig. S18) were designed to target different chosen regions in the introns where three-way junctions potentially can be forged between the genomic DNA and input RNAs (Fig. S18 and S19). Again, because these input RNAs target introns and contain no exon sequence, it rules out the possibility that induced endogenous fusion transcripts composed of exons arise from the sequence of input RNAs or the plasmids. As shown in Fig. 4A, targeting stem TMPRSS2-ETV1-A, which has the highest genomic DNA stem stability (Tm= 72°C) among this group, led to clear induction of the *TMPRSS2-ETV1* fusion transcript (Fig. 4A, lane 1). Sanger sequencing validated that the induced transcript contains *TMPRSS2* exon-1 joined with *ETV1* exon-3 (uc003ssw.4) by annotated splice sites (Fig. S20). Similar to earlier observations, targeting with sense versions of input RNAs (Fig. 4B, lane 1 vs. 2), or using antisense input RNAs designed to form three-way junctions with lower genomic DNA stem stabilities (Fig. 4A, lanes 2 to 8; Fig. S18), resulted in no detectable induction. Furthermore, the input RNA designed to target *TMPRSS2* and *ETV1* induced *TMPRSS2-ETV1* fusion but not *TMPRSS2-ERG* fusion (Fig. 4C, lane 2). Conversely, antisense-5 targeting *TMPRSS2* and *ERG* induced *TMPRSS2-ERG* fusion but not *TMPRSS2-ETV1* fusion (Fig. 4C, lane 1), indicating that fusion formation is specified by the sequence of input RNA and not secondary effects such as global genomic stability.

**Fig 4.**
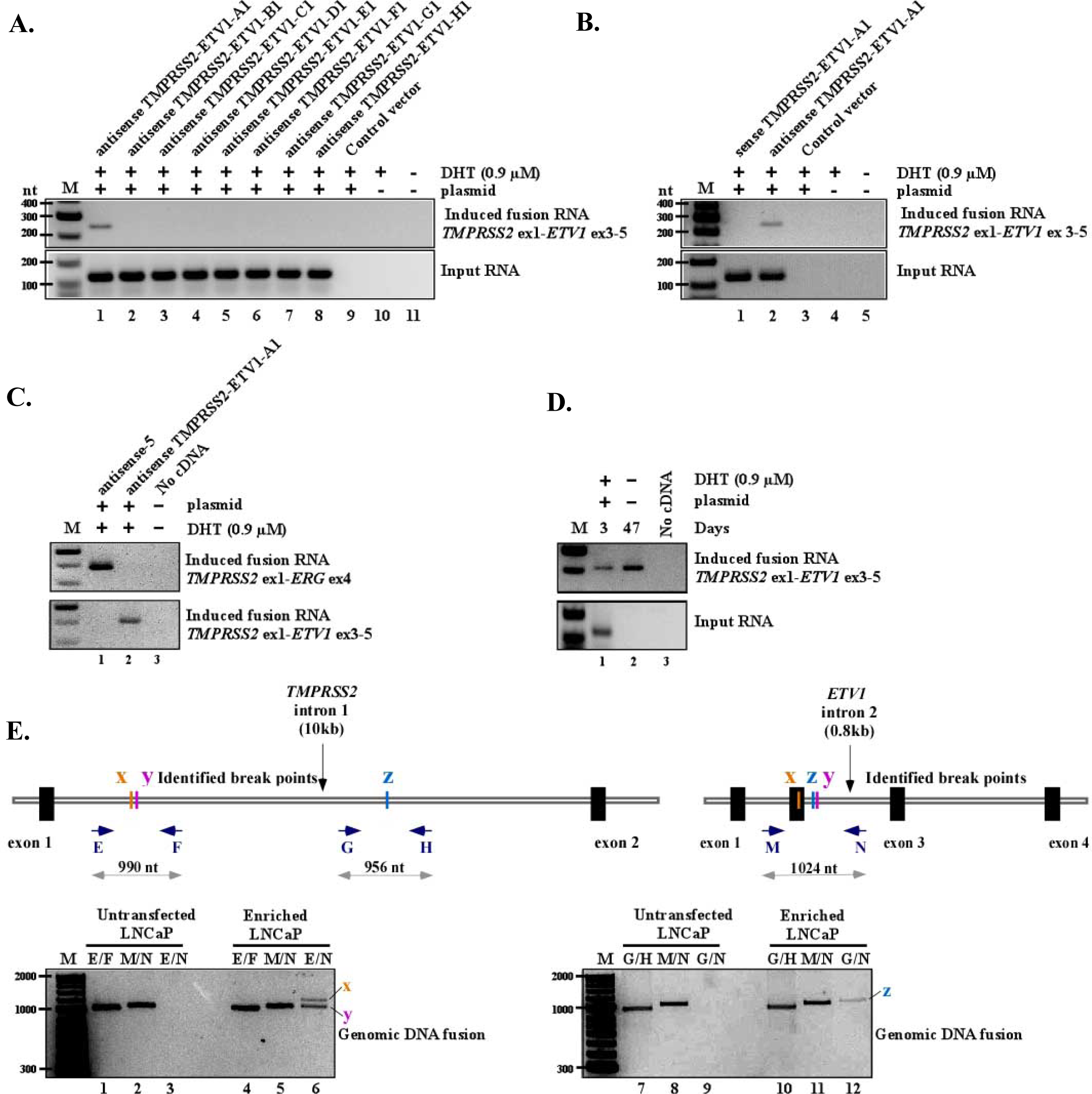
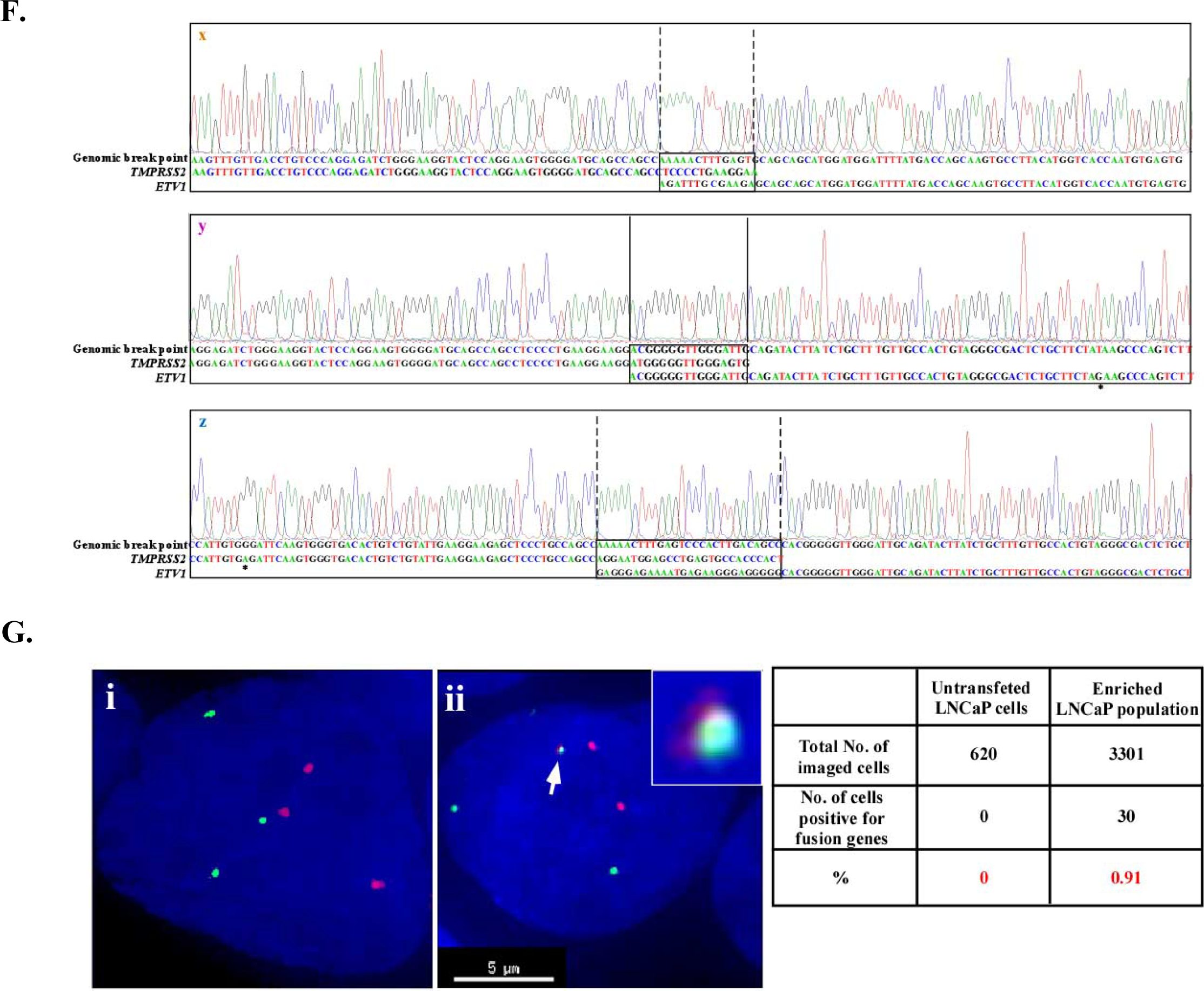
RNA-mediated inter-chromosomal gene fusion between *TMPRSS2* and *ETV1.* **A.** RT-PCR shows that only the antisense RNA designed to target stem TMPRSS2-ETV1-A, which has the highest stem stability (Tm=72°C), led to induced fusion transcript (lane 1). Antisense RNAs targeting locations with lower genomic DNA stem stabilities (lanes 2 to 8), resulted in no detectable induction. **B.** The corresponding sense input RNA targeting the same TMPRSS2-ETV1-A stem failed to induce fusion transcript (lane 1 vs. 2). **C**. Gene fusion is specified by the sequence of input RNA used. The input RNA targeting *TMPRSS2* and *ETV1* induced *TMPRSS2-ETV1* fusion but not *TMPRSS2-ERG* fusion (lane 2). Conversely, antisense-5 targeting *TMPRSS2* and *ERG* induced *TMPRSS2-ERG* fusion but not *TMPRSS2-ETV1* fusion (lane 1). **D.** RT-PCR shows the transient nature of input RNA which was present at day 3 but not day 47 post initial treatment (lower panel, lane 1 vs. 2), and the persistent nature of the induced fusion transcript (upper panel) up to 47 days post initial treatment in the enriched LNCaP population. **E.** Schematics of three identified genomic breakpoints marked as x, y, and z. The approximate location of genomic breakpoints were first determined using a primer set targeting *TMPRSS2* intron-1 multiplexed with a primer set targeting *ETV1* intron-2 (see Fig. S21). The un-rearranged wildtype *TMPRSS2* allele was then revealed by primer pair E/F (990 bp; lane 1 and 4) and G/H (956 bp; lane 7 and 10), and the un-rearranged wildtype *ETV1* allele by primer pair M/N (1024 bp; lane 2, 5, 8, 11). The genomic fusion band x (1150 bp) and y (1044 bp) amplified by fusion-specific primer pair E/N, and z (1043 bp) amplified by primer pair G/N, were present only in the induced and enriched LNCaP population but absent in untransfected LNCaP cells (lane 6 vs. 3, and lane 6 vs. 12). **F.** Sanger sequencing of the x, y, and z fusion band identified the exact genomic breakpoints. Region of microhomology at the breakpoints are boxed by solid lines, and indels by dash lines. The full-length Sanger sequences are shown in Fig. S22 to S24. **G.** FISH combined with 3D image reconstruction confirmed the gene fusion between *TMPRSS2* with *ETV1* in the enriched population. FISH probes against *TMPRSS2* gene (red) located on chromosome 21 and against *ETV1* gene (green) located on chromosome 7 were used to reveal gene fusion based on the co-localized FISH signals in reconstructed 3D images. Examples of FISH signal in an untransfected cell (i) and a cell carrying induced *TMPRSS2-ETV1* gene fusion (ii) are shown. Arrow points to the co-localized FISH signals indicative of *TMPRSS2-ETV1* fusion, which is shown at a higher magnification in the inlet. The table on the right shows the population analysis based on the 3D construction of FISH images. Approximately 0.9% of the enriched population (30 out of 3301 cells) was positive for *TMPRSS2-ETV1* fusion gene based on the co-localized FISH signals. In contrast, none of the cells from the untransfected population (0 out 620 cells) showed co-localized FISH signals.

To verify that *TMPRSS2-ETV1* as a second example of induced fusion that is indeed the consequence of genomic translocation, we propagated and enriched the induced LNCaP population for 47 days after the initial transfection of input RNA and DHT treatment (experimental procedures same as described for *TMPRSS2-ERG* enrichment in Fig. S14). The transiently expressed antisense input RNA had been degraded and was absent by day 47 (Fig. 4D, lane 1 vs. 2). The induced *TMPRSS2-ETV1* fusion transcript, however, was continuously expressed beyond day 47 (Fig. 4D, lane 2). Once again this observation indicated that the sustained expression of an induced fusion does not require the continuous presence of input RNA. Moreover, genomic PCR assays identified three distinct genomic breakpoints between *TMPRSS2* and *ETV1* gene (labeled as x, y, z in Fig. 4E) that were present only in the enriched LNCaP population but absent in untransfected LNCaP cells (Fig. 4E, lane 3 vs. 6, lane 9 vs. 12). Similar to earlier observations, no obvious sequence homology between *TMPRSS2* and *ETV1* was observed at the genomic breakpoints except for a few nt of microhomology (Fig. 4F, and Fig. S22 to S24), indicating that the gene fusion is mediated by non-homologous break repair mechanisms (Lieber, 2010; Zhang et al., 2009).

Unlike *TMPRSS2* and *ERG* that are located near each other on the same chromosome, *TMPRSS2* and *ETV1* are located on different chromosomes. Thus, gene fusion as a result of chromosomal translocation could be confirmed unequivocally by evidence of chromosomal co-localization of the latter pair. Using probes specific to *TMPRSS2* and *ETV1*, we performed fluorescence in situ hybridization (FISH) followed by deconvolution microscopic imaging of 3301 cells from the enriched LNCaP cell population and 620 cells from the control untransfected LNCaP population. Analyses of constructed 3D images showed that approximately 0.9% of the enriched population (30 out of 3301 cells) were positive for co-localization of *TMPRSS2* and *ETV1* gene in the cellular nucleus (Fig. 4G, examples of constructed 3D images are shown as movies in S25). In contrast, none of the cells from the untransfected population showed co-localized FISH signals as determined by the same 3D image criteria. These results, together with the genomic breakpoints identified by genomic PCR at single base resolution (Fig. 4E & 4F), provide strong evidence that the induced expression of the *TMPRSS2-ETV1* fusion transcript represents the consequence of gene fusion caused by chromosomal translocation.

One important observation emerging from our study is that all sense input RNAs failed to induce gene fusion. In particular, of the ten antisense input RNAs that were demonstrated to induce fusion (Fig. 1A, 1B, 2E, 4A), all of their corresponding sense counterparts failed to induce fusion (Fig. 1A, 1C, 2G, 4B). This specificity was oberserved despite the fact that sense input RNAs in theory can anneal to the same genomic sites targeted by their antisense counterparts and form similarly stable DNA/RNA hybrids. To test whether the disparity between antisense and sense is due to transcriptional activity of parental genes, we expressed the input RNAs by U6 (a pol-III promoter) for one day, followed by α-amanitin-mediated inhibition of pol-II transcription for various time periods to shut down parental gene transcription. α-amanitin was then removed to resume cellular transcription and the induction by sense vs. antisense input RNA were compared. As shown in Fig. 5 upper panel, the corresponding sense input RNAs that previously failed to induce fusion began to induce *TMPRSS2-ERG* after 12 hours of α-amanitin treatment (lane 9 and 10), and *TMPRSS2-ETV1* fusion (lane 20) after 24 hours of α-amanitin treatment. This latent induction is not a property of general cellular toxicity of α-amanitin because the toxicity caused by the same treatment actually reduced the induction by antisense input RNAs (lane 1 vs. 5 for *TMPRSS2*-*ERG*, lane 11 vs. 15 for *TMPRSS2-ETV1*). Furthermore, the input RNAs designed to target *TMPRSS2* and *ERG,* regardless of their sense or antisense nature, induced *TMPRSS2-ERG* fusion but not that of *TMPRSS2-ETV1* (lane 1 to 10). Conversely, input RNAs targeting *TMPRSS2* and *ETV1*, regardless of their directional orientation, induced *TMPRSS2-ETV1* fusion but not *TMPRSS2-ERG* fusion (lane 11 to 20). Additional control experiments using a parental plasmid vector lacking the input RNA sequences, DHT treatment without plasmid transfection, and PCR reactions without cDNA, all induced no endogenous fusion transcript under the same α-amanitin treatment (Fig. 5, lower panel). The specificity exhibited by these experiments argues against general toxicity effects. The results suggest that the antisense versus sense disparity is largely due to transcriptional conflict.

**Fig 5.**
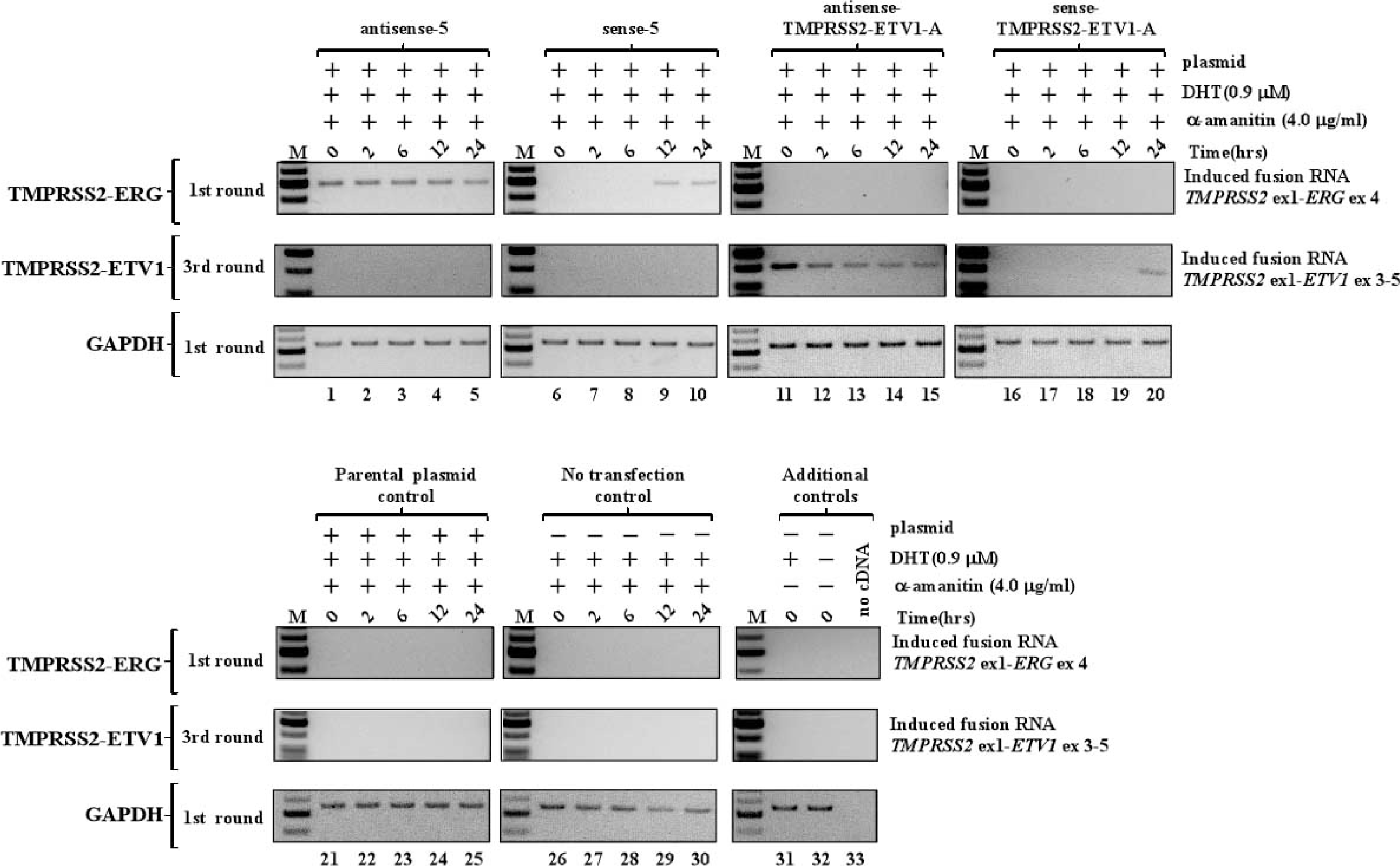
The disparity between antisense versus sense input RNA is due to transcriptional conflict. The input RNAs were expressed by U6 (a pol-III promoter) for one day, followed by α-amanitin-mediated inhibition of pol-II transcription for various time periods (0, 2, 6, 12 and 24 hrs) to shut down parental gene transcription. α-amanitin was then removed to resume cellular transcription and the induction by sense vs. antisense input RNA were compared. The corresponding sense input RNAs that previously failed to induce fusion, began to induce *TMPRSS2-ERG* (lane 9, 10) after 12 hours of α-amanitin treatment, and *TMPRSS2-ETV1* fusion (lane 20) after 24 hours of α-amanitin treatment, respectively. The results indicate that blocking the parental gene transcription by α-amanitin for a longer period (12-24 hrs for *TMPRSS2-ERG* and 24 hrs for *TMPRSS2-ETV1*) allows sense input RNA to induce gene fusion. Input RNAs used for the experiment: Antisense-5 vs. sense-5 for inducing *TMPRSS2-ERG* (upper panel, left two columns); antisense vs. sense TMPRSS2-ETV1-A for inducing *TMPRSS2-ETV1* (upper panel, right two columns). As controls, transfecting cells with a parental plasmid containing no input RNA sequences (lower panel, left column), cells without transfection (lower panel, center column), and DHT and α-amanitin controls (lower panel, right column), also failed to induce fusion. In addition, GAPDH is used as internal control for the amount of RNA in each lane.

With the plausibility of RNA-mediated gene fusion established, we then sought evidence that specific endogenous cellular RNAs can act as the ‘initiator’ to induce *TMPRSS2-ERG* fusion, which is found in ~50% of prostate cancers. To identify candidate cellular initiator RNAs, we analyzed an available mRNA-seq database consisting of prostate tumors and matched benign tissues (Kannan et al., 2011). However, we found no evidence of perfect endogenous antisense chimeric RNAs in which the *TMPRSS2* sequence was joined to any *ERG* sequence by discernable 5’ and 3’ splice sites in the antisense orientation. This suggests that if endogenous initiator RNAs do exist, they might arise from unrelated genomic sources that coincidentally resemble an imperfect chimeric RNA antisense to both *TMPRSS2* and *ERG*. In a detailed study to be presented elsewhere (manuscript in preparation), we have performed thermodynamic calculations of RNA/DNA hybrids to identify cellular RNAs with partial sequence complementarity to the *TMPRSS2* and *ERG* genes. We identified that *AZI1* mRNA (also known as *CEP131*) (Aoto et al., 1997; Aoto et al., 1995) could form high affinity RNA/DNA hybrids with *TMPRSS2* and *ERG* genomic sequences. As shown in Fig. 6A, overexpressing full-length *AZI1* mRNA (3619 nt, uc002jzn.1) induced the *TMPRSS2-ERG* fusion transcript in LNCaP cells. The induction was observed at a physiologically relevant concentration (40 nM) of DHT (Fig. 6A, lane 4). Furthermore, expression of exon16-17 of *AZI1*, a short 220 nt segment containing an imperfect sequence antisense to *TMPRSS2* and *ERG*, was sufficient to induce *TMPRSS2-ERG* fusion (Fig. 6B, lane 2). This result suggests that the induction of gene fusion is mediated by an RNA sequence that resides in exon16-17 and requires no AZI1 protein. Consistent with previous observations that sense input RNAs are ineffective for the fusion process, the expression of exon16-17 in the antiparallel orientation also failed to induce *TMPRSS2-ERG* fusion (Fig. 6B, lane 3).

**Fig 6.**
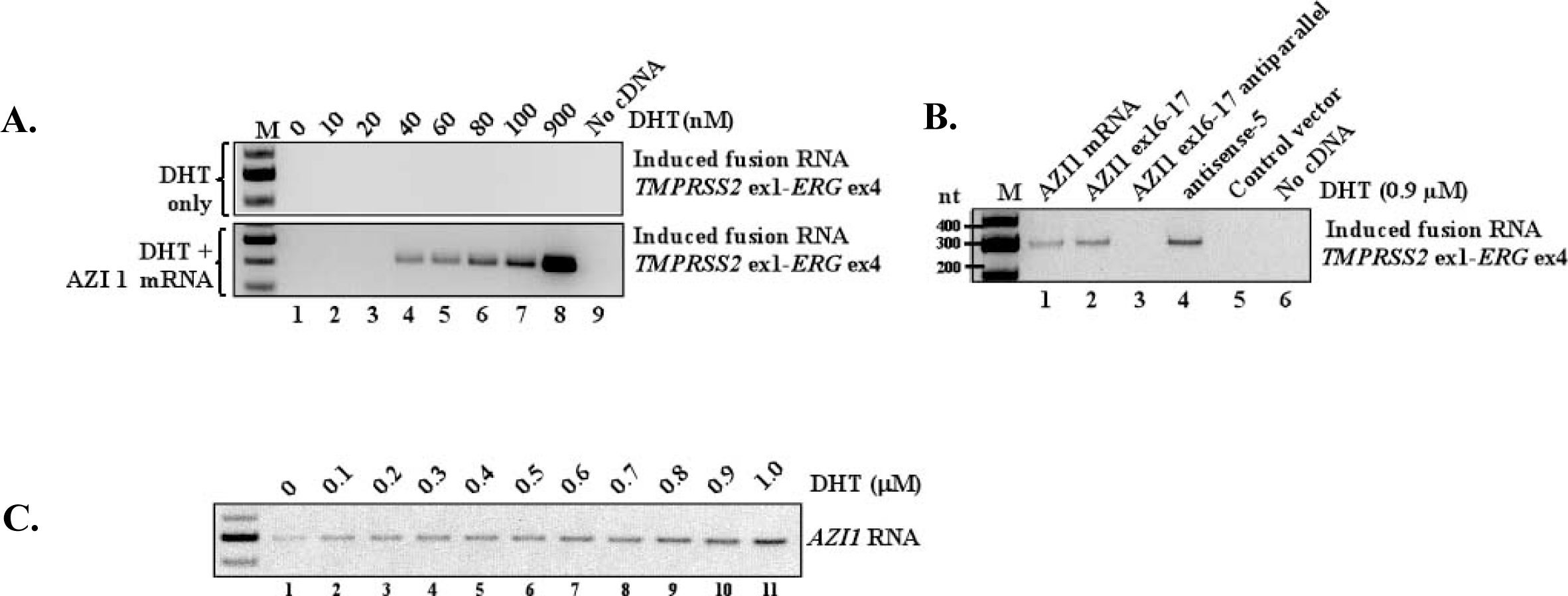
Specific endogenous RNA acts as the ‘initiator’ RNA to mediate gene fusion. **A.** Expression of full-length *AZI1* mRNA for 3 days induced *TMPRSS2-ERG* fusion. The induction occurred at physiologically relevant concentrations of DHT as low as 40 nM (lane 4). Three-rounds of nested PCR were performed to reveal the lowest amount of DHT that permits *AZI1*-mediated fusion induction. **B**. Expression of *AZI1* exon 16-17 (lane 2), but not its antiparallel sequence (lane 3), led to *TMPRSS2-ERG* fusion. Inductions by full-length AZI1 (lane 1) and by antisense-5 (lane 4) were used as positive controls. **C.** Endogenous *AZI1* RNA is expressed in LNCaP cells at low level and the expression is upregulated by DHT treatment.

## Discussion

In summary, this report provides the first evidence that expression of a chimeric RNA can drive the formation of gene fusions in mammalian cells. Hence, we propose that “the cart before the horse” hypothesis concerning fusion gene causation is mechanistically plausible. Our data support a model (shown in Fig. 7) where the initiator RNA with chimeric sequence invades chromosomal DNA to stabilize a transient RNA/DNA duplex using DNA sequences located in two distant genes. Resolution of such an RNA/DNA duplex by DNA repair mechanisms might yield the final gene fusion through recombination in regions prone to DNA breaks (Fig. 7A). Such events were rare in the initial population of transfected cells (1 in 10^3^ or 10^4^ cells occurred within 3 days). However, the necessary machinery is clearly present in normal prostate epithelial cells prior to malignant transformation. If the resulting gene fusion (such as *TMPRSS2-ERG)* provides a growth advantage, a single affected cell among billions of cells in a normal prostate tissue may proliferate abnormally and eventually contribute to cancer formation. Identifying such initiator RNAs might provide novel insights into early disease mechanisms, as well as the discovery of new preventive and therapeutic strategies to combat cancer.

**Fig 7.**
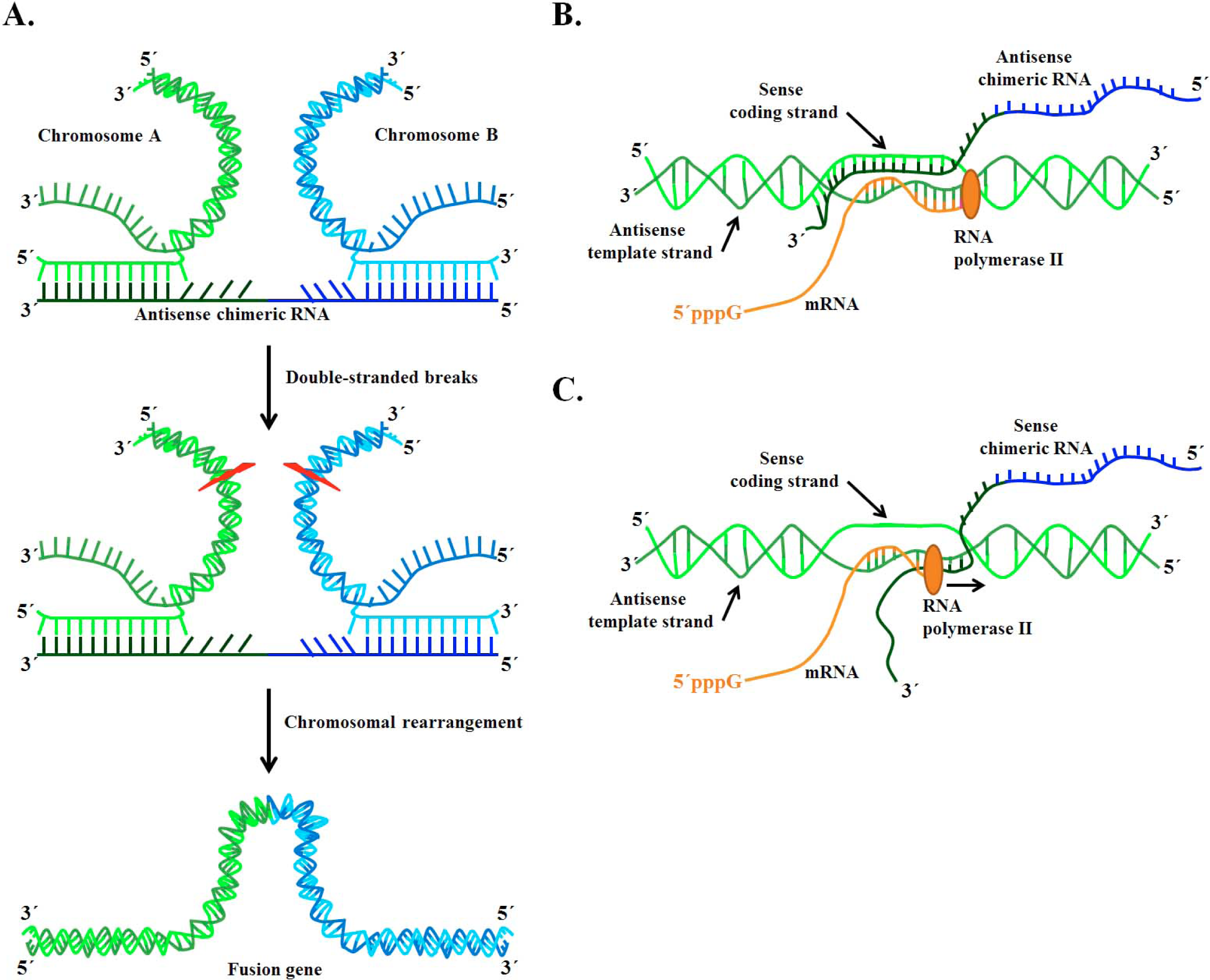
A model of RNA-mediated gene fusion in mammalian cells. **A.** Our data support a model where the initiator RNA with chimeric sequence invades chromosomal DNA to stabilize a transient RNA/DNA duplex using DNA sequences located in two distant genes. Resolution of such an RNA/DNA duplex by DNA break/repair mechanisms might yield the final gene fusion through recombination in regions prone to DNA breaks. Such events however are infrequent (1 in 10^3^ or 10^4^ cells in the experimental cell population), and are dependent of the presence of hormone DHT in the case of LNCaP cells. **B and C.** The data also indicate that it is the antisense rather than sense chimeric RNAs that effectively drive gene fusion, and that this disparity can be explained by transcriptional conflict produced by the transcriptional activity of parental genes. The sense chimeric RNAs forming DNA/RNA hybrids with antisense strands of genomic DNA (the template strand used for transcription) are likely be frequently “bumped” off by RNA polymerase and unable to stabilize the structures required for initiating genomic arrangements. The proposed RNA-driven model may provide a mechanism that can ‘specify’ gene fusion partners in early disease stages, and could have fundamental implications in the biology of mammalian genome stability, as well as gene editing technology via mechanisms native to mammalian cells.

Contrary to the previous “cart before the horse” model (Rowley and Blumenthal, 2008; Zaphiropoulos, 2011), our results do not support the postulation that a sense fusion mRNA derived from trans-splicing between two pre-mRNAs effectively directs gene fusion. Expressing sense input RNAs mirroring the trans-spliced mRNA failed to induce fusion in LNCaP cells (Fig. 1A and S2). Of ten antisense RNAs that were demonstrated to be capable of inducing fusion (Fig. 1A, 1B, 2E, 4A, 6B), all of their corresponding sense RNAs failed to induce fusion (Fig. 1A, 1C, 2G, 4B, 6B). This occurred even though the sense RNAs could, in theory, anneal to the same genomic sites targeted by their antisense counterparts and form similar DNA/RNA hybrids when paired with the antisense strand of genomic DNA. As demonstrated in Fig. 5, this antisense versus sense disparity can be explained by transcriptional conflict (Fig. 7B and 7C). Because the *TMPRSS2* promoter is highly active in LNCaP cells, sense chimeric RNAs forming DNA/RNA hybrids with antisense strands of genomic DNA (the template strand used for transcription) would be frequently “bumped” off and unable to stabilize the structures required for initiating genomic arrangements. The mechanistic basis for the antisense versus sense RNA, and whether these phenomena can be generalized, remain to be investigated.

Our results also do not support the hypotheses that antisense input RNAs, acting as a docking station, mediate trans-splicing by base-pairing with both endogenous sense parental pre-mRNAs, or by bringing the parental genes in close proximity thus facilitating trans-splicing of parental pre-mRNAs transcribed from two genomic loci. Both mechanisms would require the continuous presence of antisense input RNAs to sustain the expression of induced fusion transcripts. Yet we showed that the induced fusion expression has a permanent nature and requires no continuous presence of input RNAs (Fig. 3B and Fig. 4D). Furthermore, in the case of *TMPRSS2-ERG* there is no detectable *ERG* parental RNA as raw material (Fig. 3A) to account for the trans-splicing models. Moreover, sense input RNAs, which are not complimentary to the sense parental pre-mRNAs thus cannot act as their docking station, are able to induce the fusion transcripts after a brief period of transcriptional inhibition, again arguing against the docking model. On the contrary, the genomic breakpoints identified by genomic PCR and chromosomal co-localization provided by FISH, strongly support that the induced expression of fusion transcript is largely the consequence of gene fusion resulting from chromosomal translocation. While prior works have shown that infrequent *TMPRSS2-ERG* fusions can by induced through genotoxic stress such as gamma radiation in the presence of DHT that increases double-stranded DNA breaks (Lin et al., 2009; Mani et al., 2009), such mechanisms of general genotoxicity fail to account for the specificity of gene fusion partners found in cancer. This report is the *first* to demonstrate RNA-mediated gene fusion in mammalian cells, and provides an RNA-driven mechanism that can account for the ‘specificity’ of gene fusion partners that were selected to undergo gene fusion in early disease stages. The results may represent a pathological example in a broad spectrum of potential RNA-mediated genome rearrangements and could have fundamental implications in the biology of mammalian genome stability, as well as gene editing technology via mechanisms native to mammalian cells.

## Author contributions

SKG and LL conducted the experiments, SKG and LY designed the experiments, SKG and LY conceived the project and wrote the manuscript.

## Acknowledgments

We thank Richard Kelley, Michael Ittmann, Richard Sifers and Tom Cooper for critical suggestions, Christine Shiang for cloning AZI1 ex16-17, Olga Dakhova, Jianghua Wang, and Michael Ittmann for providing the cell lines. Sachin Kumar Gupta has been supported by CPRIT training grant RP160283. Laising Yen has been supported by Duncan Cancer Center Pilot Grant, CPRIT HIHRRA RP160795, and NIH R01EB013584. We would also like to thank Radhika Dandekar, Fabio Stossi, and Michael Mancini for assisting with microscopy, with the support by the Integrated Microscopy Core at Baylor College of Medicine with funding from the NIH (DK56338, and CA125123), CPRIT (RP150578) the Dan L. Duncan Comprehensive Cancer Center, and the John S. Dunn Gulf Coast Consortium for Chemical Genomics.

## Supplementary Information

**Fig. S1.**
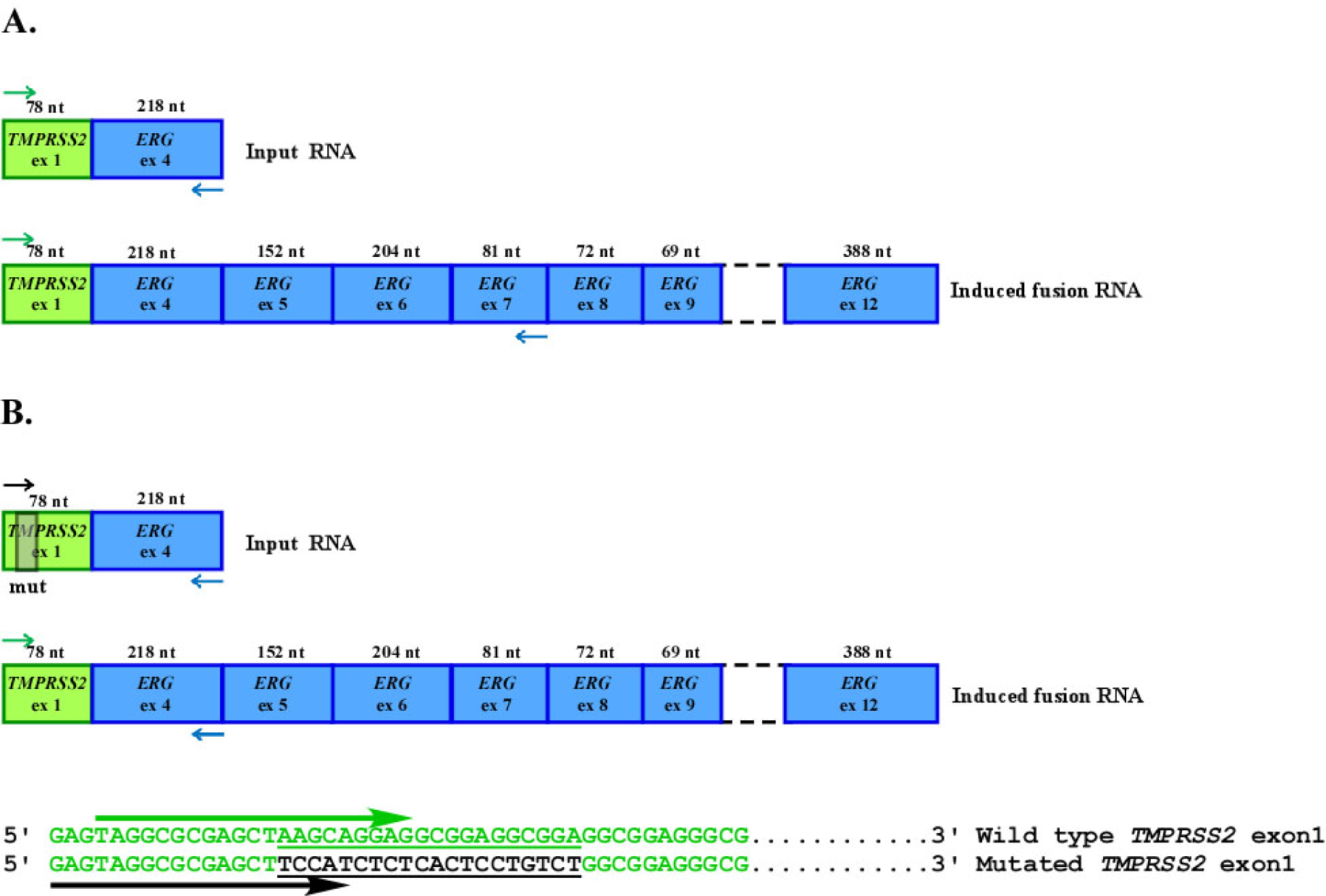
Primer design used to distinguish induced endogenous fusion RNAs from exogenous input RNAs. **A.** The endogenous full-length *TMPRSS2-ERG* fusion RNA most commonly found in prostate cancer consisting of *TMPRSS2* exon 1 (78 nt, uc002yzj.3) spliced to *ERG* exons 4-12 (1289 nt, uc021wjd.1). In Fig. 1A in the main text, the exogenous input RNAs expressed from plasmids consists of *TMPRSS2* exon-1 (78 nt) fused to *ERG* exon-4 (218 nt). The RT-PCR results shown in Fig. 1A and 1B (middle panel) in the main text were performed using a forward primer on *TMPRSS2* exon-1 paired with a reverse primer on *ERG* exon-7, thereby specifically amplifying the endogenous fusion RNA transcribed from the *TMPRSS2-ERG* fusion gene, but not from transfected plasmids or input RNAs because both lack *ERG* exon-7. **B.** Because RNA-induced fusion is a low frequency event, we developed a more efficient PCR method to detect induced endogenous fusion RNA using a primer pair targeting *TMPRSS2* exon-1 and *ERG* exon-4. To distinguish induced endogenous fusion RNAs from exogenous input RNAs, mutations were introduced at nucleotide positions 16-35 of the input RNAs (gray box in *TMPRSS2* exon-1), allowing specific amplifications of induced endogenous fusion RNAs or exogenous input RNA using specific forward primers. The lower panel shows the wild type *TMPRSS2* exon-1 sequence in green. The mutated area in input RNA is shown in black and underlined. The primer used to specifically recognize induced fusion RNA is shown as the green arrow and was used in most of the figures in the main text. The primer for input RNA is denoted by the black arrow and was used in Fig. 1B, 1C, 1D, 1E, Fig. 2B, and Fig. 3B in the main text.

**Fig. S2.**
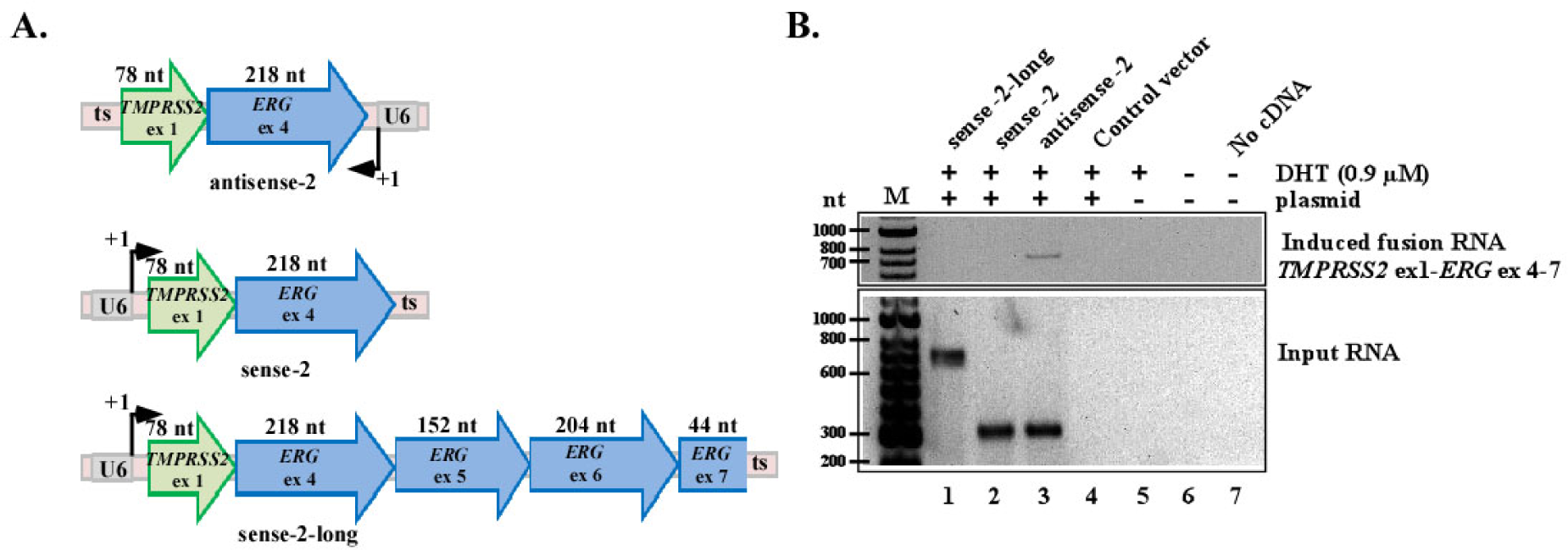
Sense input RNAs that mimic trans-spliced products failed to induce fusion transcripts. **A.** Schematics of the designed input RNAs consisting of *TMPRSS2* (uc002yzj.3) and *ERG* (uc021wjd.1) exons. Upper panels: antisense-2 and sense-2 input RNA consisting of *TMPRSS2* exon-1 (78 nt) and *ERG* exon-4 (216 nt). Lower panel: sense-2-long RNA consisting of *TMPRSS2* exon-1 (78 nt) with *ERG* exons-4, -5, -6 and partial -7 (618 nt). ts: transcriptional stop “TTTTTT” for the U6 promoter. **B.** RT-PCR detection of induced fusion transcript (top gel) and input RNA (bottom gel). For induced fusion transcript, RT-PCR was performed using a forward primer on *TMPRSS2* exon-1 paired with a reverse primer on the 3’ end of *ERG* exon-7 (see Fig. S1A), thereby specifically amplifying the endogenous fusion RNA transcribed from the *TMPRSS2-ERG* fusion gene, but not from transfected plasmids or input RNAs because both lack the 3’ end of *ERG* exon-7. Control vector lacking input RNA sequence (lane 4), DHT treatment only (lane 5), and PCR reactions lacking cDNA (lane 7) were used as negative controls. Antisense RNA is capable of inducing the fusion transcript (lane 3), whereas sense RNAs failed to induce fusion transcripts regardless of their lengths (lanes 1 and 2).

**Fig. S3.**
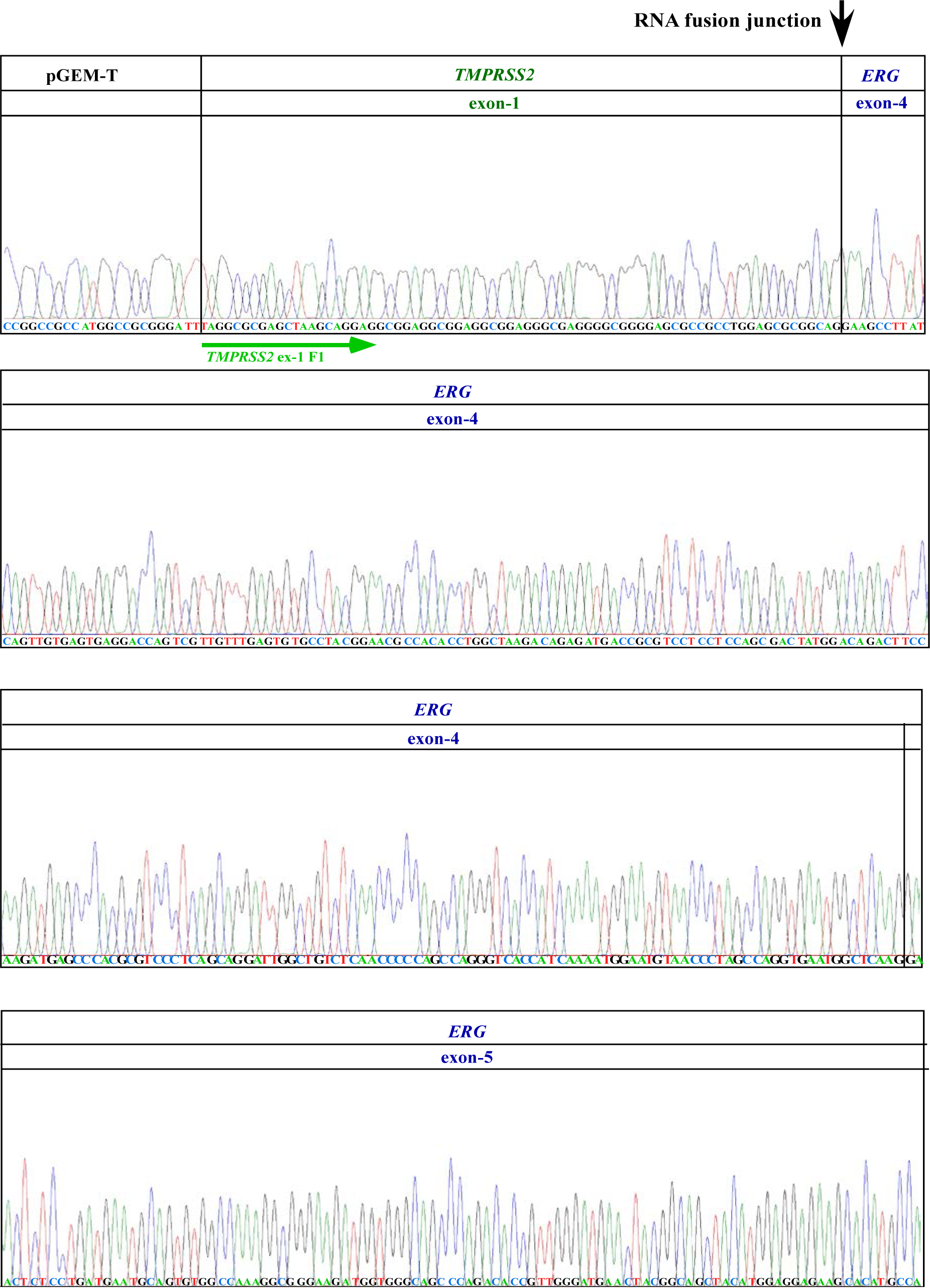
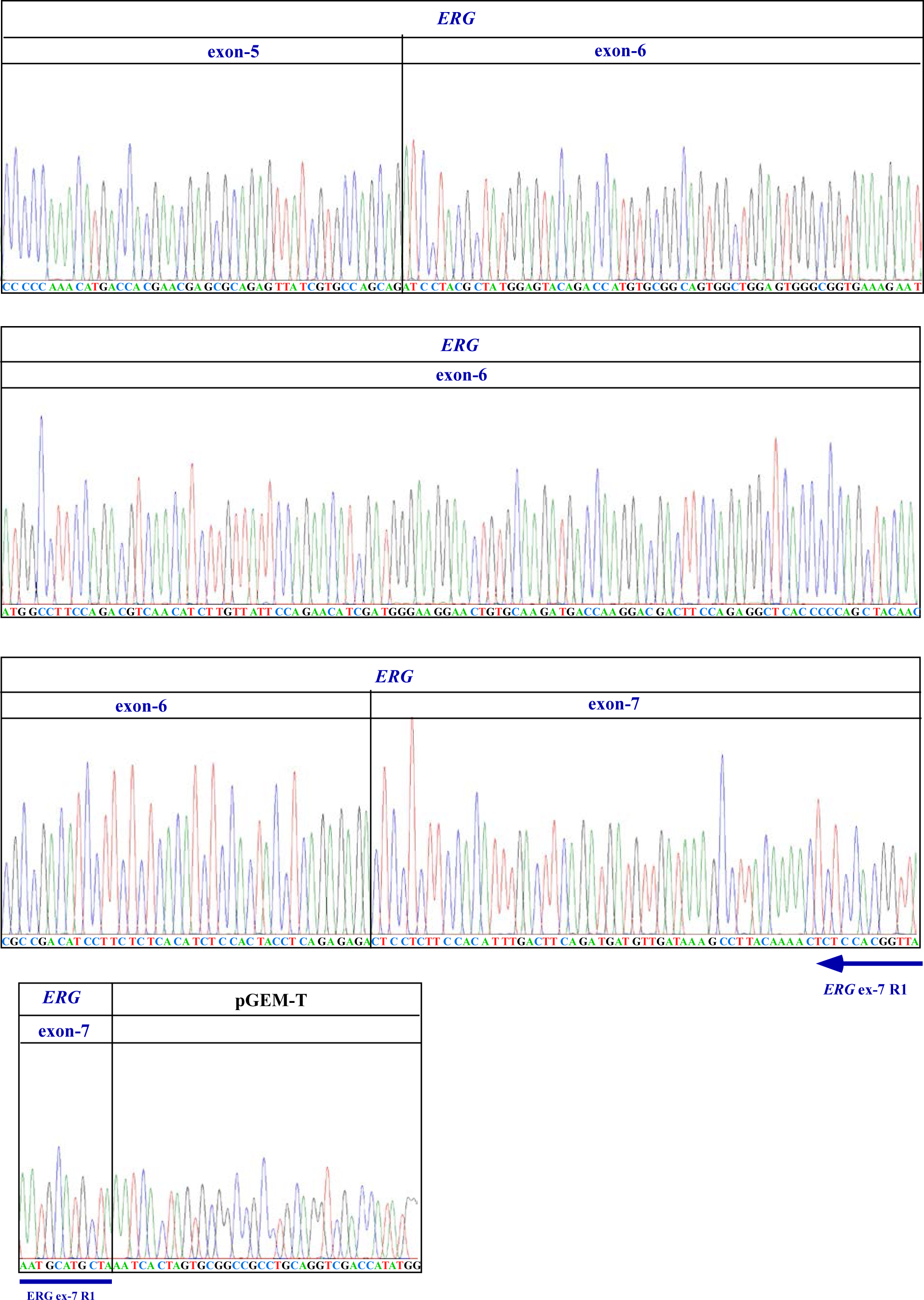
Sanger sequencing confirmation of the induced *TMPRSS2-ERG* fusion transcript in LNCaP cells mediated by antisense-2 input RNA. Induced fusion transcripts were converted to cDNA using oligo dT primers and PCR was performed using primers targeting *TMPRSS2* exon-1 and *ERG* exon-7 as described in Fig. S1. The RT-PCR assay relies on the primer targeting *ERG* exon-7 to specifically amplify the induced fusion transcript but not the input RNA, which lacks *ERG* exon-7 sequence. The locations of the primers used to amplify the PCR product are shown as the green and blue arrows for *TMPRSS2* exon-1 and *ERG* exon-7, respectively. The RNA fusion junction between *TMPRSS2* exon-1 and *ERG* exon-4 is indicated by the black arrow. The pGEM-T plasmid was used for cloning the cDNA inserts. Sanger sequencing chromatograms confirmed that the induced fusion transcript contains *TMPRSS2* exon-1 fused to *ERG* exons-4, -5, -6, and -7, and that these exons were joined by splicing at the annotated splice sites as would be expected of mature endogenous *TMPRSS2-ERG* fusion mRNA. This result confirms that expression of an input RNA can lead to specific fusion transcript in human cells.

**Fig. S4.**
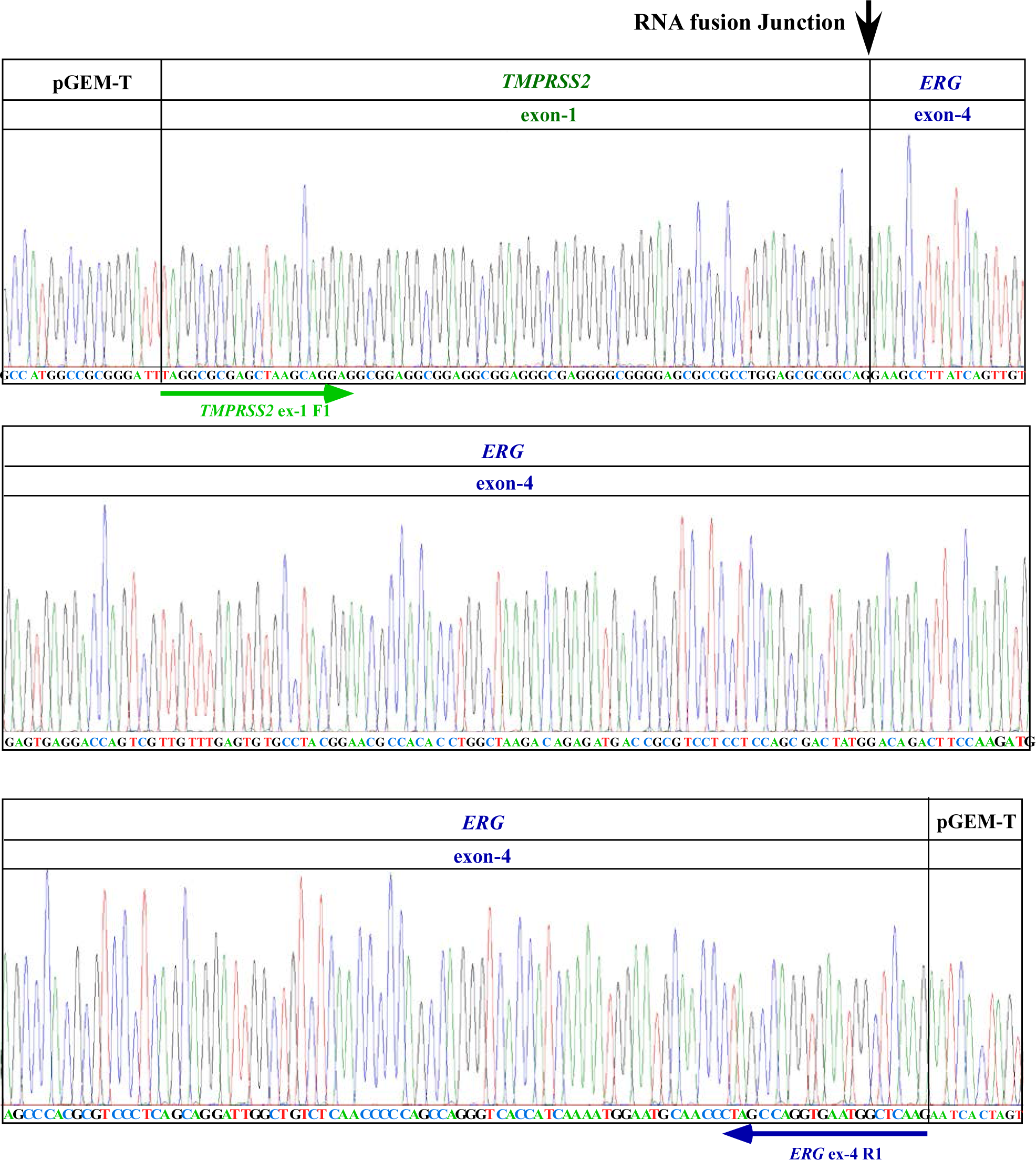
Sanger sequencing confirmation of the induced *TMPRSS2-ERG* fusion transcripts mediated by antisense-5 input RNA and amplified using an optimized primer pair. Induced fusion transcripts were converted to cDNA using oligo dT primers. RT-PCR was performed with primers targeting *TMPRSS2* exon-1 and *ERG* exon-4, which led to higher amplification efficiency. This RT-PCR assay relies on the primer targeting *TMPRSS2* exon-1 to specifically amplify the induced fusion transcript but not the input RNA, which contains the mutated region described in Fig S1B. The locations of the primers are shown as green and blue arrows for *TMPRSS2* exon-1 and *ERG* exon-4, respectively. The RNA fusion junction between *TMPRSS2* exon-1 and *ERG* exon-4 is indicated by the black arrow. The pGEM-T plasmid was used for cloning the cDNA inserts. Sanger sequencing chromatograms confirmed that the induced fusion transcript contained *TMPRSS2* exon-1 fused to *ERG* exon-4, as would be expected of mature endogenous *TMPRSS2-ERG* fusion mRNA. We developed this primer pair to optimize the efficiency of detecting low levels of induced endogenous *TMPRSS2-ERG* fusion mRNA, and these primers were used for most of the RT-PCR assays.

**Fig. S5.**
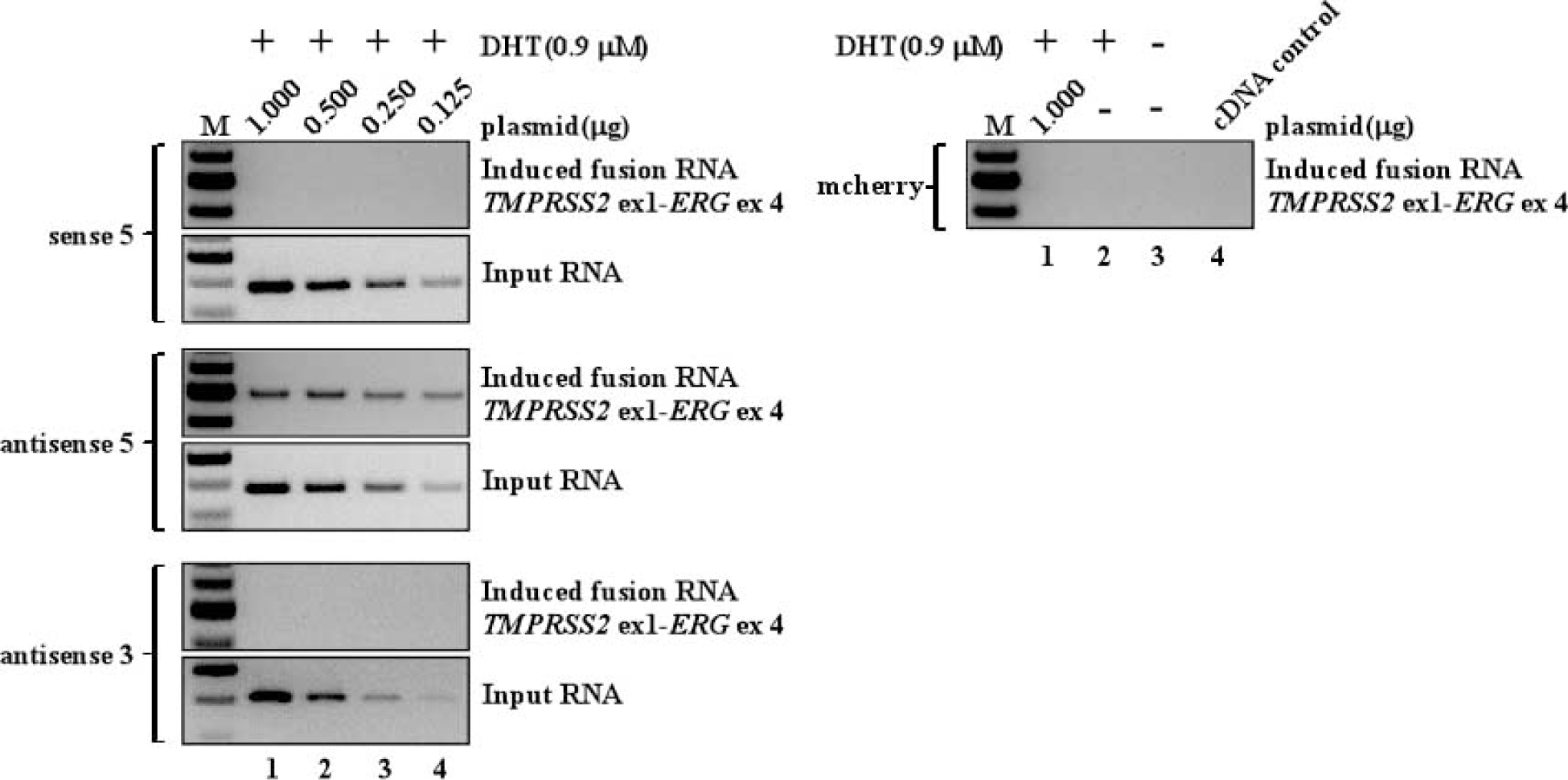
Antisense-5 but not its corresponding sense-5 induces *TMPRSS2-ERG* gene-fusion. Different amount of plasmids (1.000μg, 0.500μg, 0.250μg and 0.125μg) expressing either sense-5 or antisense-5 were transfected to produce different amount of corresponding input RNAs in LNCaP cells. To maintain transfection efficiency, mCherry control plasmid was added to each transfection to make the final amount of plasmid to 1.0 μg. Induction was done for three days in the presence of 0.9 μM DHT. RT-PCR was performed to detect the levels of both induced *TMPRSS2-ERG* transcript and input RNA. RT-PCR results shows that even very low amount of antisense-5 input RNA is sufficient to induce *TMPRSS2-ERG* gene fusion (middle panel, lane 4) whereas very high level of sense-5 input RNA fails to induce *TMPRSS2-ERG* gene fusion (upper panel, lane 1). Thus, it is not the amount of the input RNA but the orientation of the input RNA (antisense vs. sense) that is important for fusion induction. Antisense-3, which is unable to induce fusion, was used as a negative control. mCherry alone was also transfected as a negative control. Additional controls (right panel): a vector expressing mCherry lacking input RNA sequence (lane 1), DHT treatment only (lane 2), and PCR reaction lacking cDNA (lane 4).

**Fig. S6.**
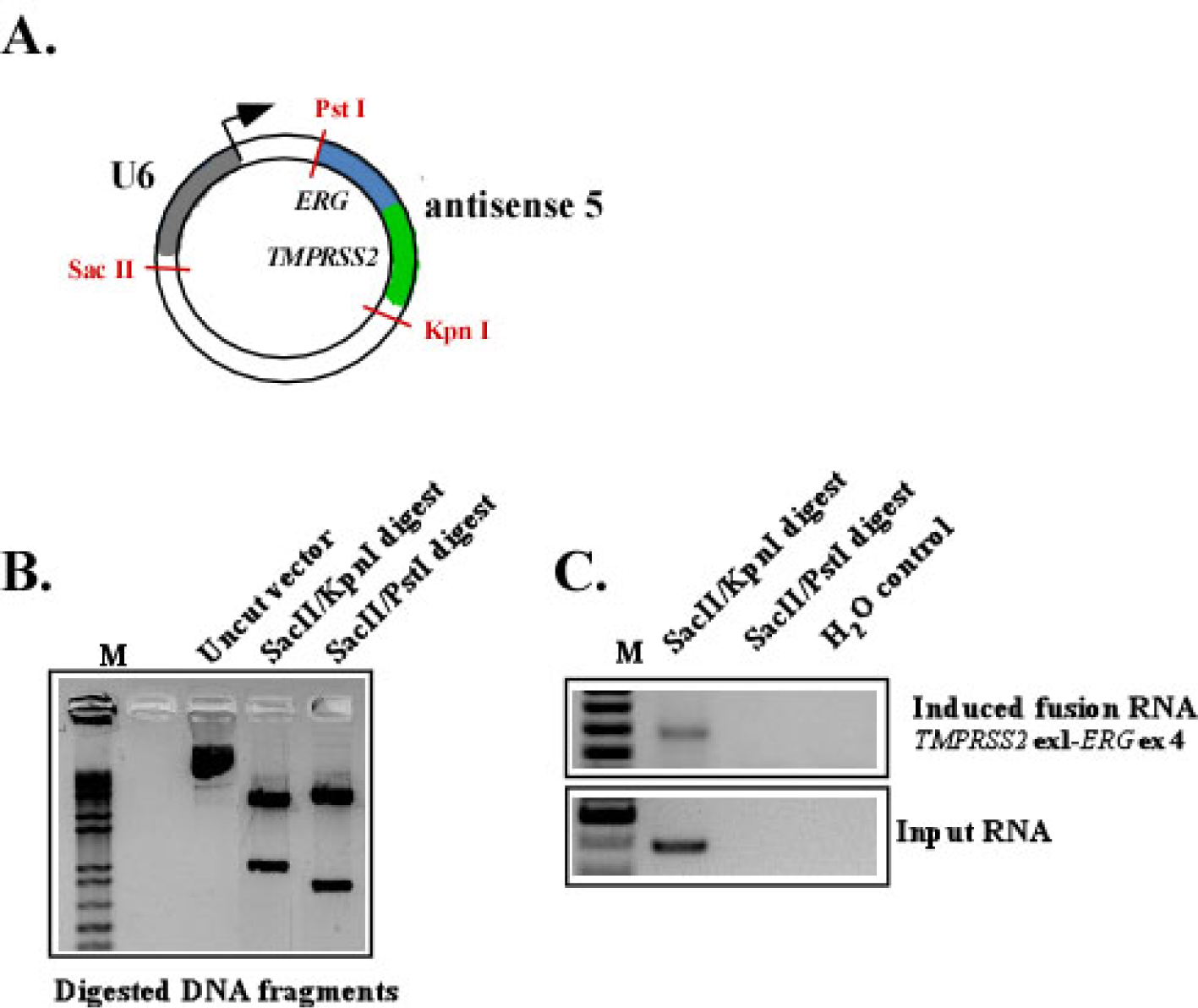
Expressed input RNA, but not the DNA plasmid, induces fusion transcript. **A.** Schematic of the expression plasmid containing a U6 promoter and antisense-5 sequence. Unique restriction sites are shown in red. The arrow indicates the transcriptional start. The plasmid was digested by SacII/KpnI, leaving U6 with antisense-5, or with SacII/PstI, separating U6 from antisense-5. **B.** Confirmation of complete digestion was visualized by agarose gel. Uncut plasmid vector was used as control. **C.** Induction of *TMPRSS2-ERG* fusion after transfection of digested plasmid fragments. The *TMPRSS2-ERG* fusion transcript was induced only when the U6 promoter was attached to antisense-5 sequence, indicating that it is the input RNA expressed by the U6 promoter, not the transfected DNA plasmid per se, that leads to fusion induction.

**Fig. S7.**
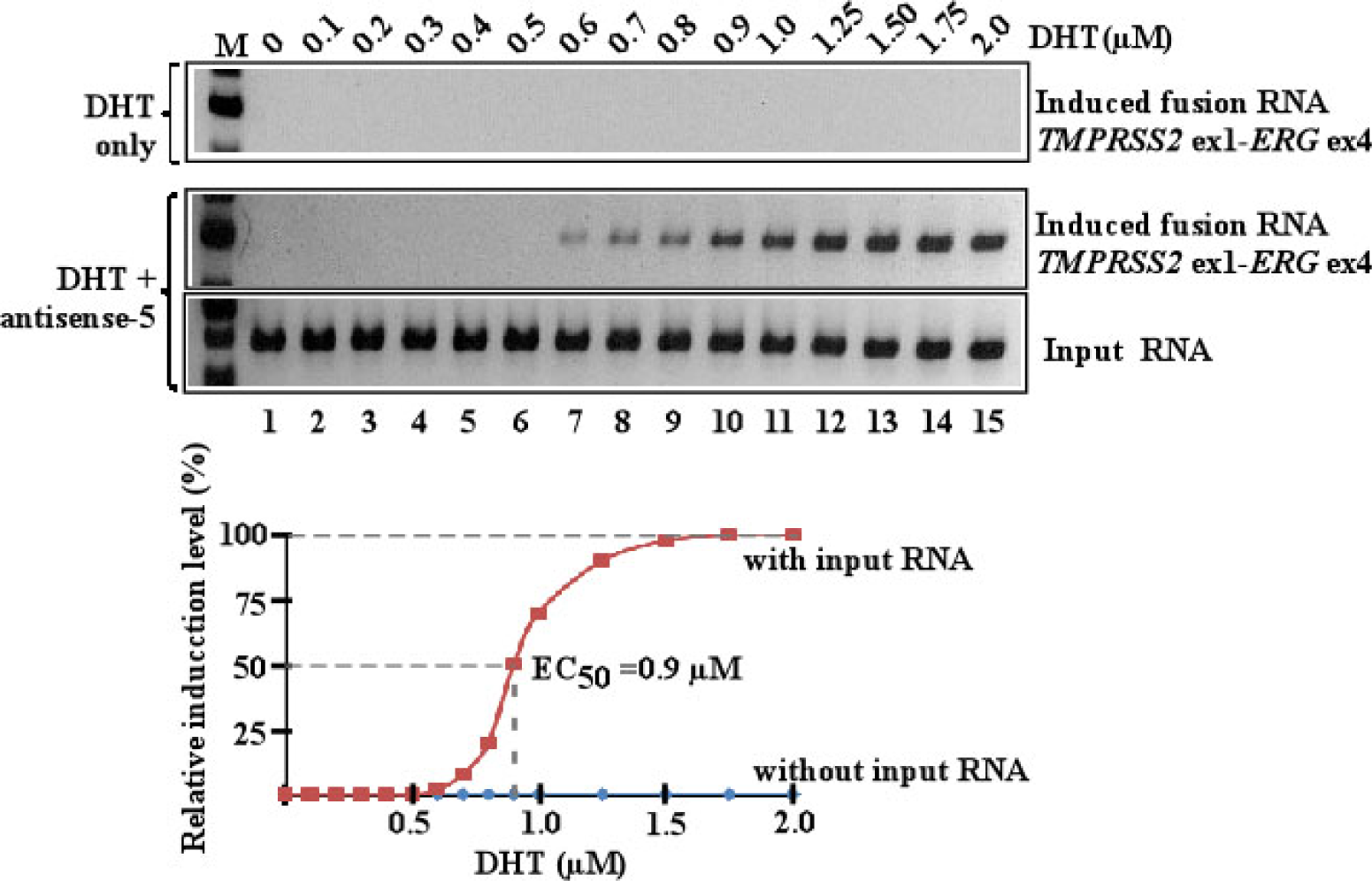
DHT dosage effect in facilitating RNA-mediated gene fusion as assayed by one round RT-PCR. To determine the DHT concentrations that facilitate RNA-mediated gene fusion detectable by one round RT-PCR, we generated a dose-response curve using LNCaP cells transfected with antisense-5 and treated with different DHT concentrations for 3 days. Relative RT-PCR intensities of induced fusion transcript bands as generated by one round RT-PCR were plotted against DHT concentrations. In the presence of antisense-5, fusion induction is detectable at ~0.6 μM DHT, reaches its maximum at 1.5 μM DHT, with the effective concentration that yields 50% of maximal induction (EC_50_) at ~0.9 μM. In the absence of antisense-5, no induction was observed in the entire DHT range tested up to 2.0 μM (upper panel), indicating that DHT alone is ineffective in inducing *TMPRSS2*-*ERG* fusion. To streamline our experimental procedures, we used 0.9 μM DHT (the EC_50_ concentration) for 3 days followed by one round of RT-PCR as our standard assay in the main text. However, it is important to note that all key experiments were also performed under physiologically relevant DHT concentrations (<100 nM), and in those cases three-rounds of nested PCR were used to reveal the lowest amount of DHT required. As shown in Fig. 1D and Fig. 6A in the main text, gene fusion events induced by antisense-5 occurred at physiologically relevant DHT concentrations as low as 20 nM, and gene fusion events induced by endogenous *AZI1* mRNA occurred at as low as 40 nM DHT.

**Fig. S8.**
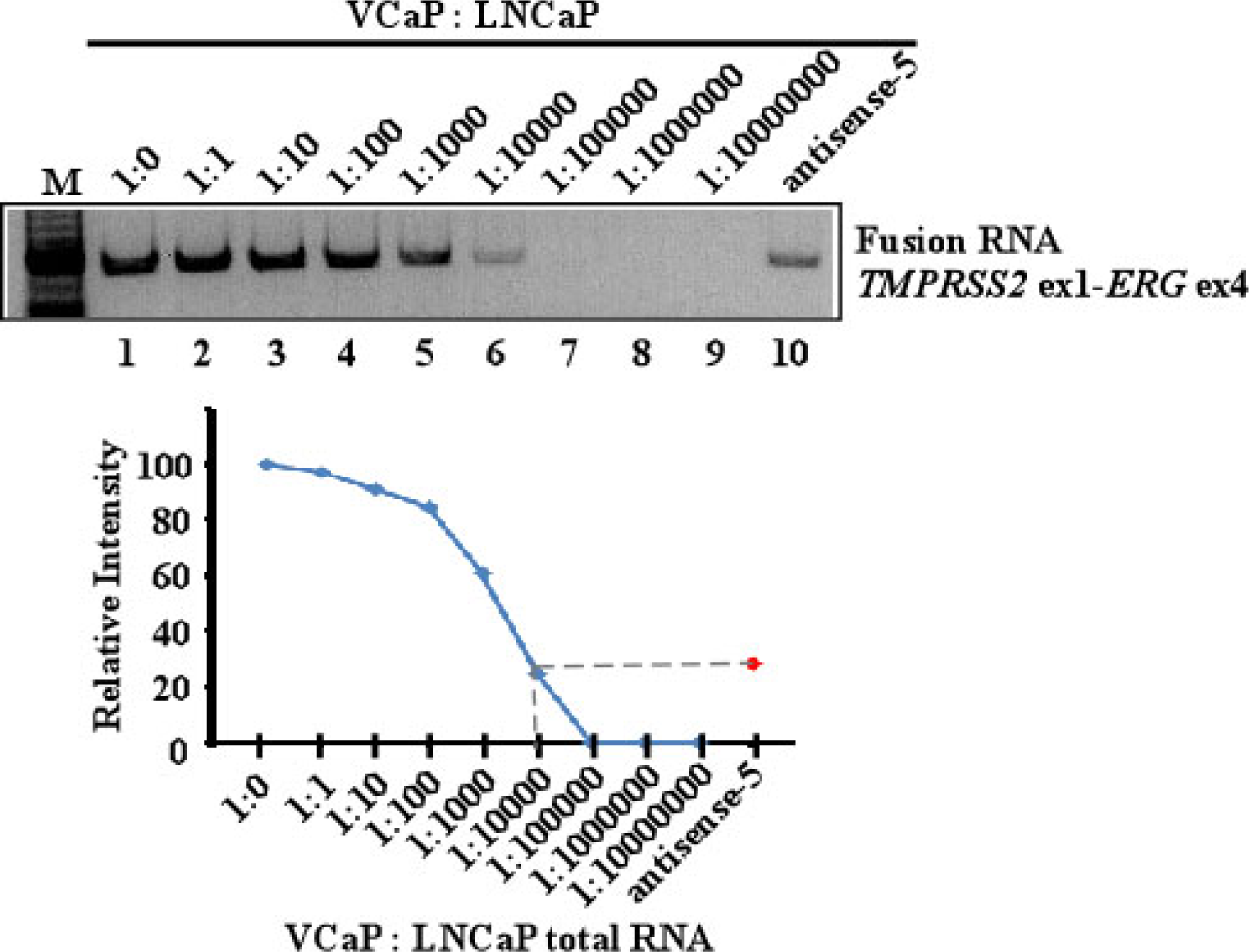
Estimation of the percentage of LNCaP cells transfected with antisense-5 that carry the induced *TMPRSS2-ERG* fusion. To estimate the percentage of LNCaP cells that underwent *TMPRSS2*-*ERG* gene fusion mediated by antisense-5 at 0.9 μM DHT (the EC_50_), we mixed various amounts of total RNA from VCaP cells that express *TMPRSS2*-*ERG* (Mertz et al., 2007; Teles Alves et al., 2013) with the total RNA from untransfected LNCaP cells in ratios of 1:1 up to 1:10^7^. A standard dilution curve was determined using the relative RT-PCR intensity of the *TMPRSS2-ERG* fusion transcript. The relative intensity of induction by antisense-5 in LNCaP cells, as assayed under the same RT-PCR conditions, is shown as a red dot that corresponds to an equivalent dilution of 1:10^3^ to 1:10^4^, suggesting that approximately 1 in 10^3^ or 10^4^ LNCaP cells is positive of *TMPRSS2-ERG* fusion.

**Fig. S9.**
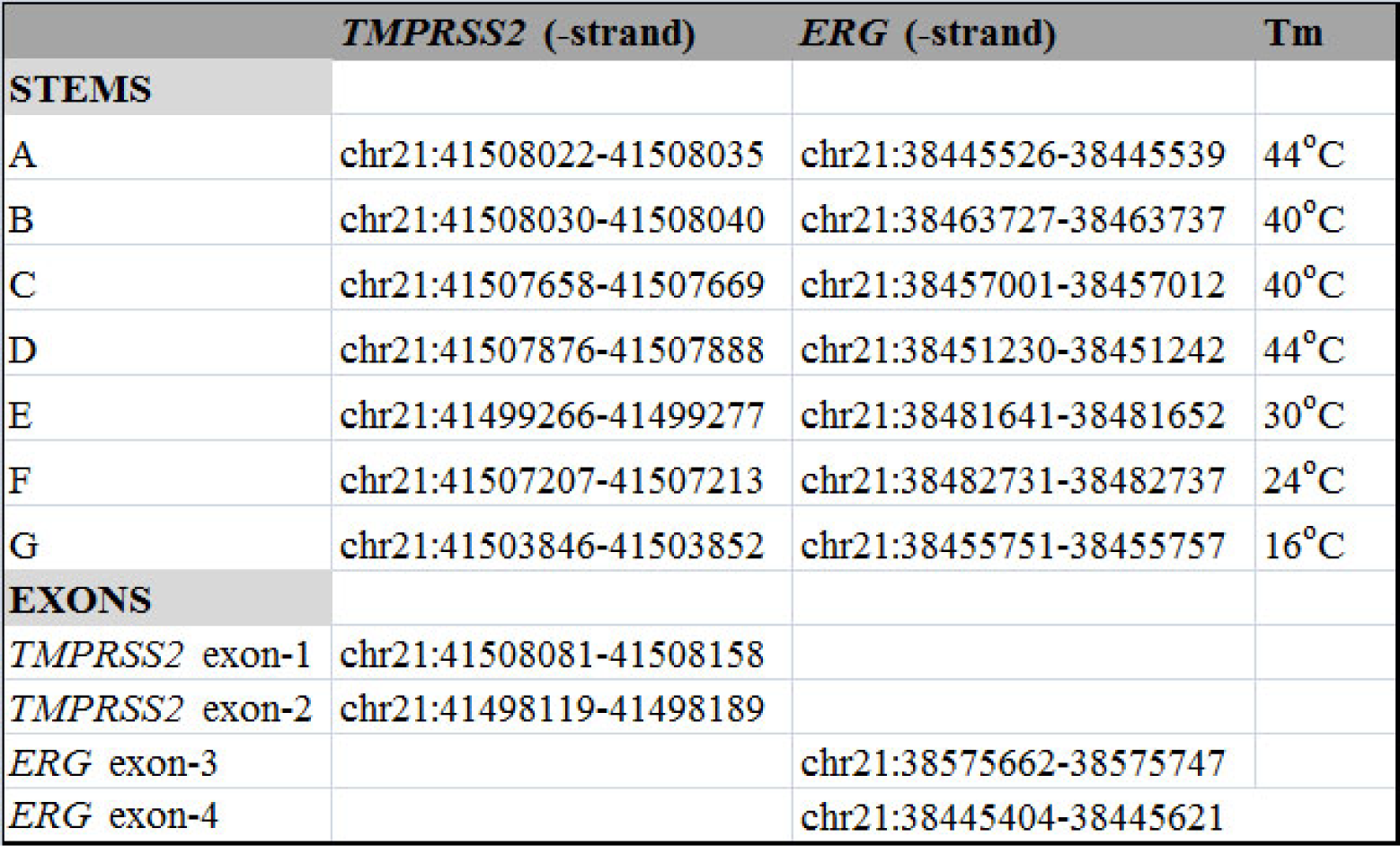
Genome coordinates and Tm for the putative genomic stems described in Fig. 2C and 2D of the main text. Genome coordinates are based on the GRCh38/hg38 version. Coordinates of the exons (*TMPRSS2* exon-1 and -2; *ERG* exon-3 and -4) are also shown.

**Fig. S10.**
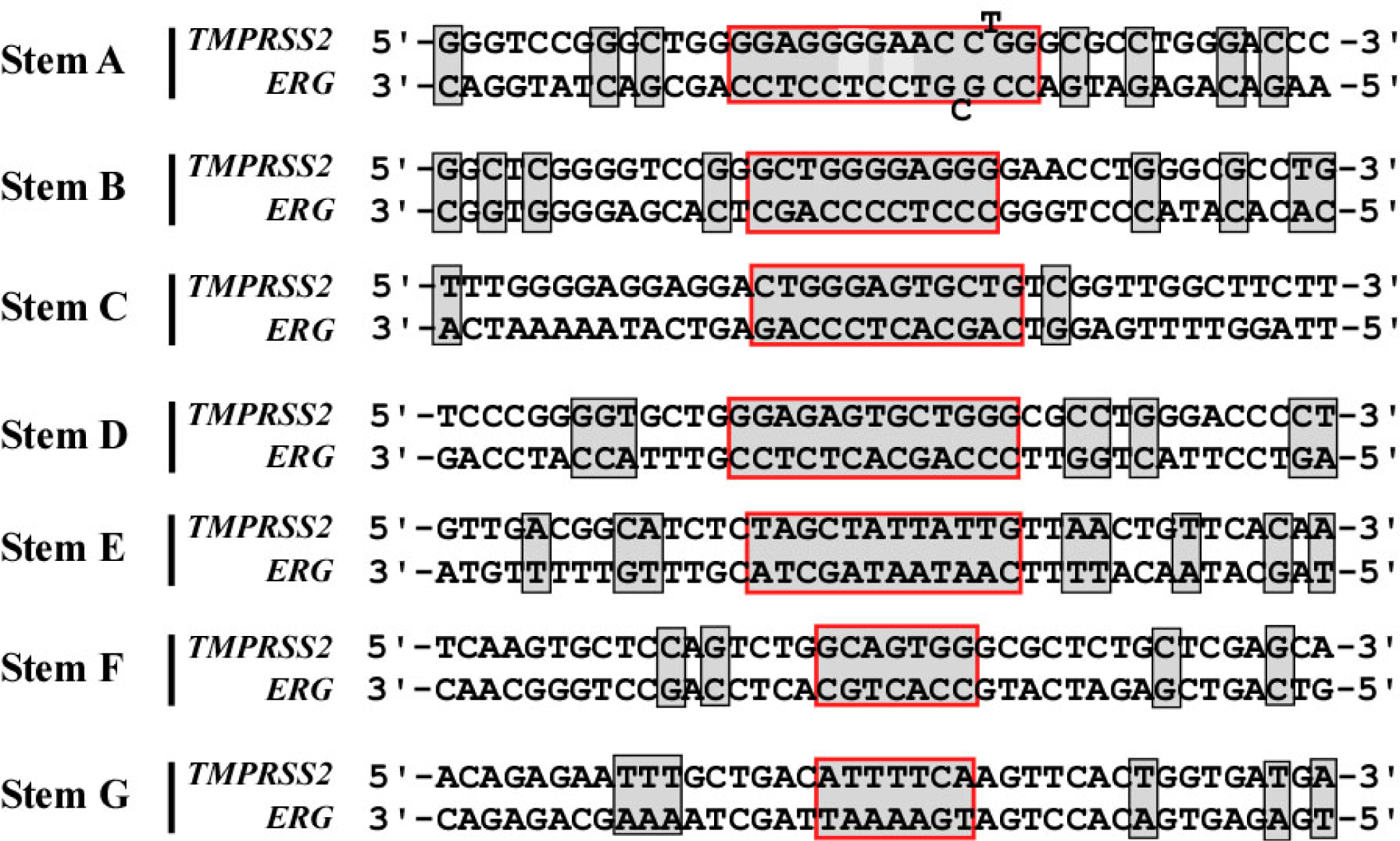
Complementary base pairing of the genomic DNA sequences comprising the stems and their flanking sequences. The sense strand of genomic sequences from *TMPRSS2* and *ERG* that form the putative stems (in red boxes) with various degrees of stability as identified by BLAST analyses. Regions showing perfect complementary base pairing (A-T and G-C) are shaded in gray.

**Fig. S11.**
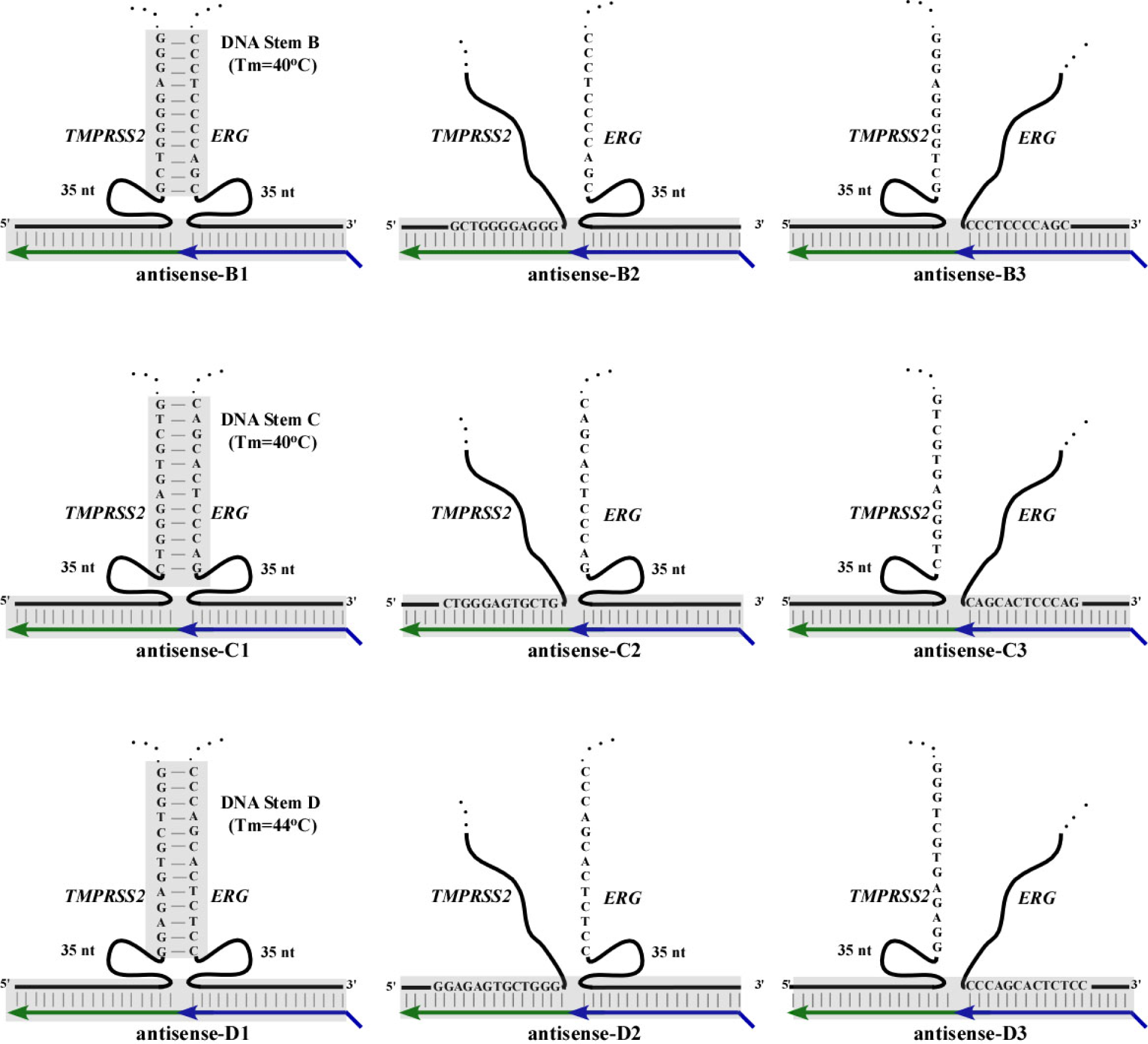
Antisense input RNAs designed to disrupt the genomic stems B, C, and D. Stem B (upper row). Stem C (middle row). Stem D (lower row). Left column: Illustrations show the three-way junction formed by genomic DNA and corresponding antisense input RNA (antisense-B1, C1, and D1). Middle column: Disruption of stems via the *TMPRSS2* side using tailor-made input RNAs (antisense-B2, C2, and D2) that directly hybridize to the *TMPRSS2* stem sequence. Right column: Disruption of stems via the *ERG* side using tailor-made input RNAs (antisense-B3, C3, and D3) that directly hybridize to the *ERG* stem sequence. Shaded regions indicate base pairing. The *TMPRSS2* portion of the input RNA is shown in green and the *ERG* portion is in blue.

**Fig. S12.**
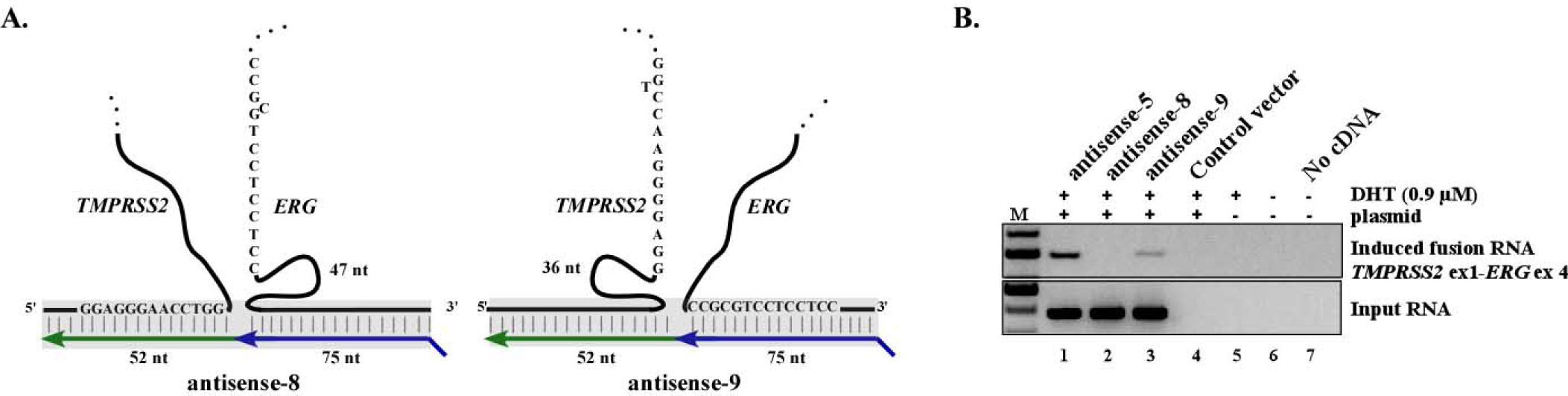
Antisense input RNAs designed to disrupt the genomic stems A. **A.** Disruption of the putative genomic stem using tailor-made input RNAs that directly hybridize to one side of the stem A. **B.** RT-PCR assays of fusion transcripts induced by the input RNAs illustrated in A. Interfering with three-way junction formation eliminated (lane 2) or significantly reduced the induction (lane 3). Controls include: vector lacking input RNA sequence (lane 4), DHT treatment only (lane 5), and PCR reaction lacking cDNA (lane 7).

**Fig. S13.**
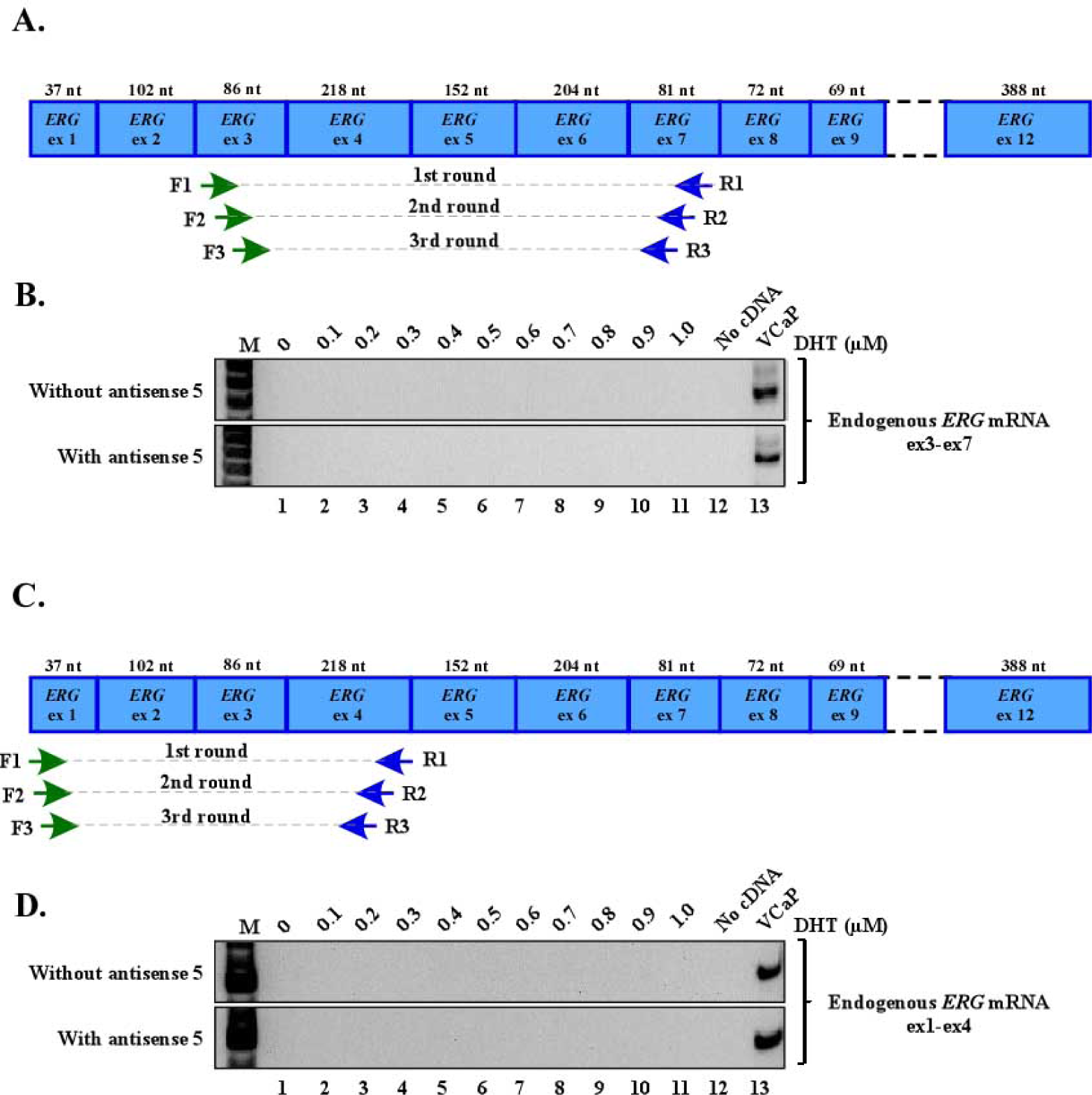
Endogenous *ERG* mRNA is not detected in LNCaP cells by three-rounds of nested RT-PCR. **A.** Nested primer pairs used to detect low levels of endogenous *ERG* mRNA from LNCaP cells. Forward primers for *ERG* exon-3 are shown in green whereas reverse primers for *ERG* exon-7 are shown in blue (primer sequences are listed in the Materials and Methods). **B.** Endogenous *ERG* mRNA was not detected in LNCaP cells in the presence or absence of DHT or antisense-5 (lane 1 to 11) by three-rounds of nested RT-PCR. cDNA prepared from VCaP cells which express *ERG* mRNA was used as positive control (lane 13). **C** and **D.** Similarly, using forward primers on *ERG* exon-1 and reverse primers on *ERG* exon-4 also failed to detect endogenous *ERG* mRNA by three-rounds of nested RT-PCR (lane 1 to 11, primer sequences are listed in the Materials and Methods). These primer sets were chosen because they specifically amply endogenous *ERG* mRNA but not *TMPRSS2-ERG* fusion transcript that contains *ERG* exon-3 to exon-12.

**Fig. S14.**
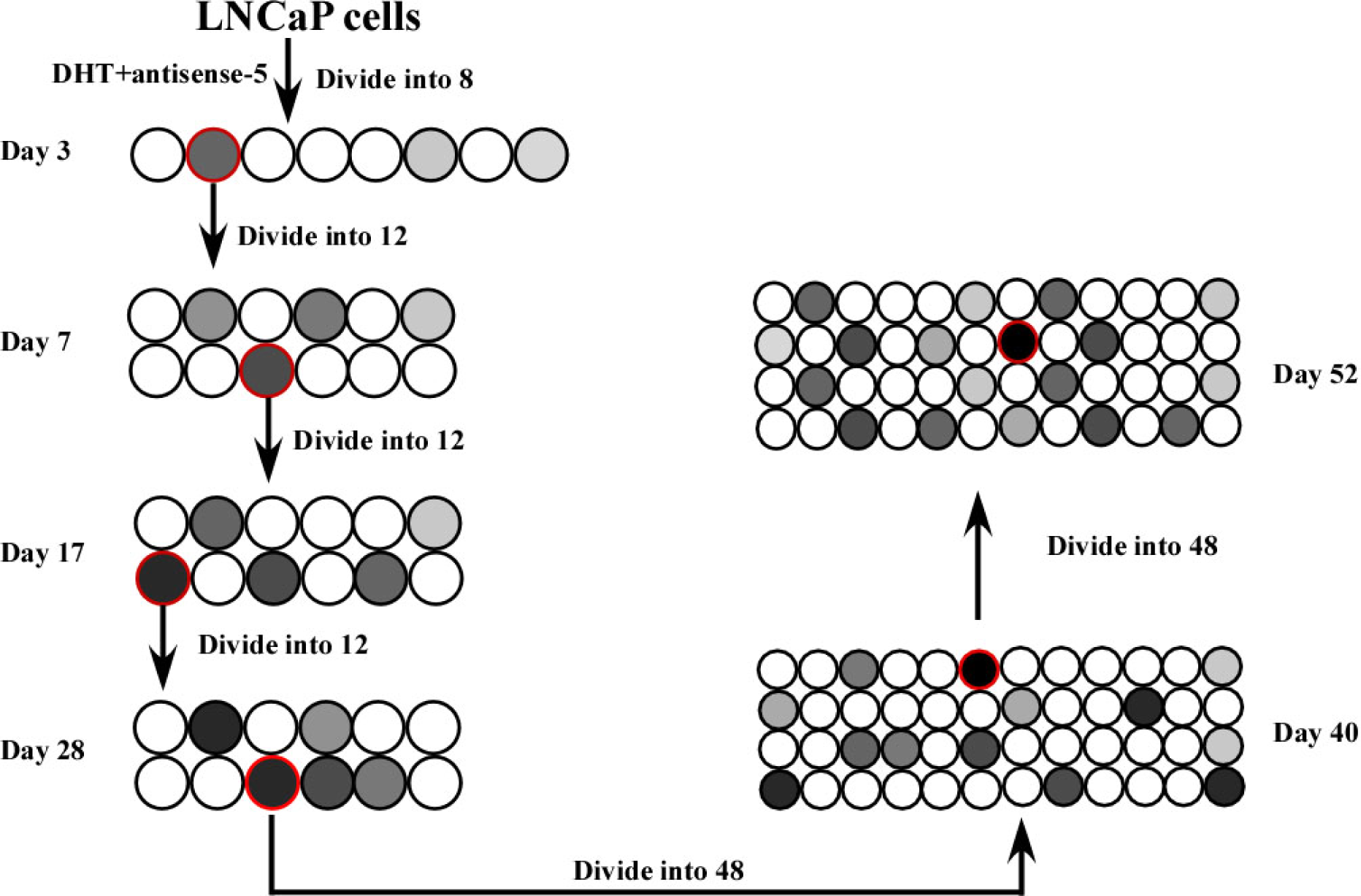
Procedure used to propagate and enrich the induced LNCaP population for 52 days in the absence of DHT. LNCaP cells were transfected on day 1 with antisense-5 and incubated for 3 days with DHT. Cells were then divided into sub-populations. Half of the cells from each sub-population were harvested for RT-PCR assays to detect the induced fusion transcript whereas the other half was used to propagate the population. The sub-population containing the highest intensity of induced fusion transcripts (circled in red) was selected for continuous propagation for the next round of division and RT-PCR assays. The process was continued for 52 days. RT-PCR was performed to detect both induced fusion transcripts and antisense input RNA during each division as shown in Fig. 3B.

**Fig. S15.**
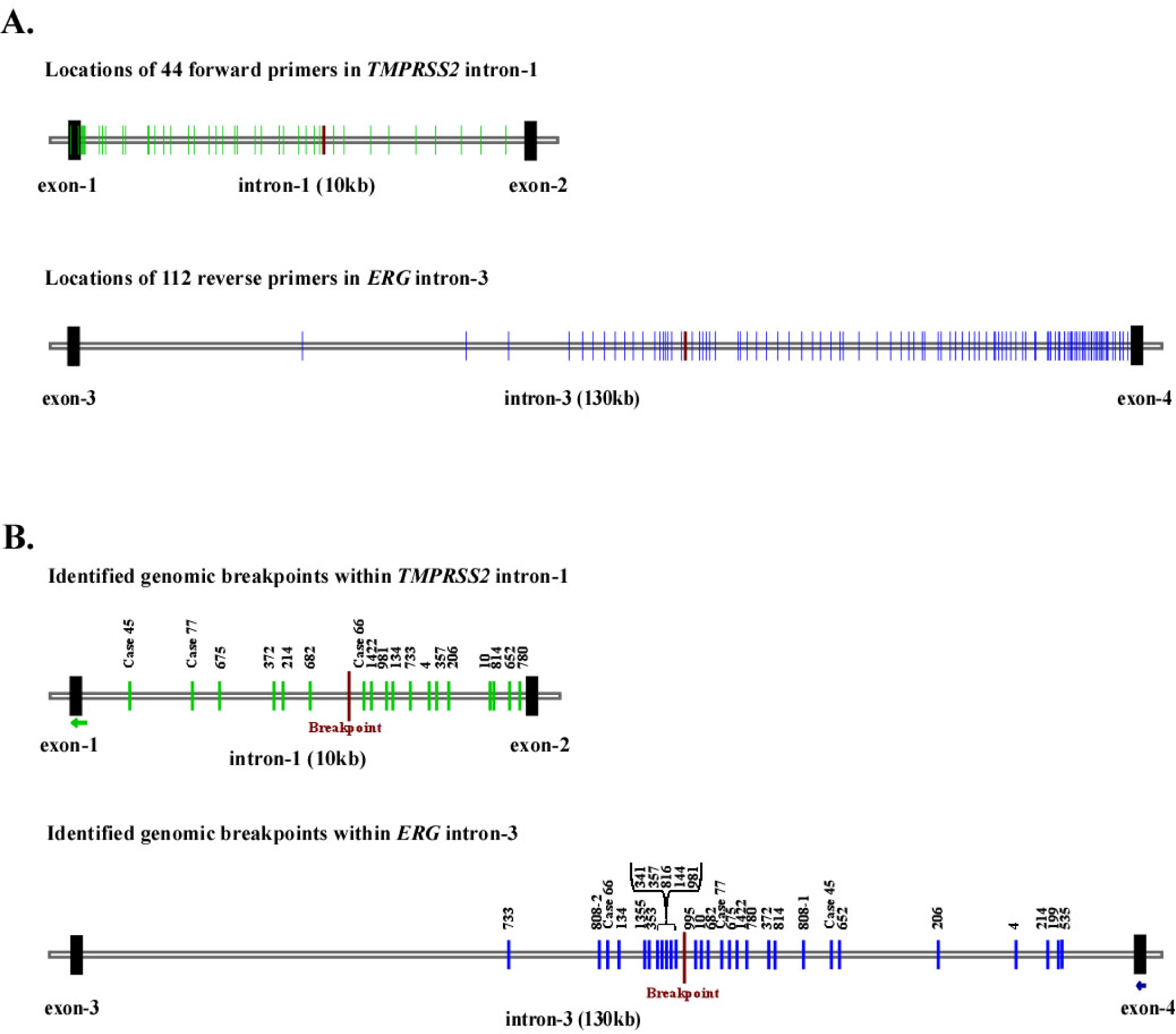
Locations targeted by primers for mapping the genomic breakpoint and locations of identified *TMPRSS2-ERG* genomic breakpoints. **A.** The locations of 44 forward primers (green) spacing across *TMPRSS2* intron-1 (~10 kb) and 112 reverse primers (blue) spacing across *ERG* intron-3 (~130 kb). *ERG* intron-3 is significantly larger than that of *TMPRSS2* intron-1. Each vertical line represents a target location by a primer or a set of nested primers. The primers targeting *ERG* intron-3 are designed to concentrate in the region near exon-4 and in the hot spot regions previously identified from prostate cancer patients. The specific primers that amplify the genomic breakpoint shown in Fig. 3D in the main text are labeled as red. **B.** Locations of identified *TMPRSS2-ERG* genomic breakpoints. Locations of previously identified *TMPRSS2-ERG* genomic breakpoints from prostate cancer patients (Weier et al.) are shown in green in *TMPRSS2* intron-1 and blue in *ERG* intron-3. The number on top of each location is the ID number of a patient. The genomic breakpoint identified in our study is shown in red. Short arrows indicate sites targeted by antisense-5.

**Fig. S16.**
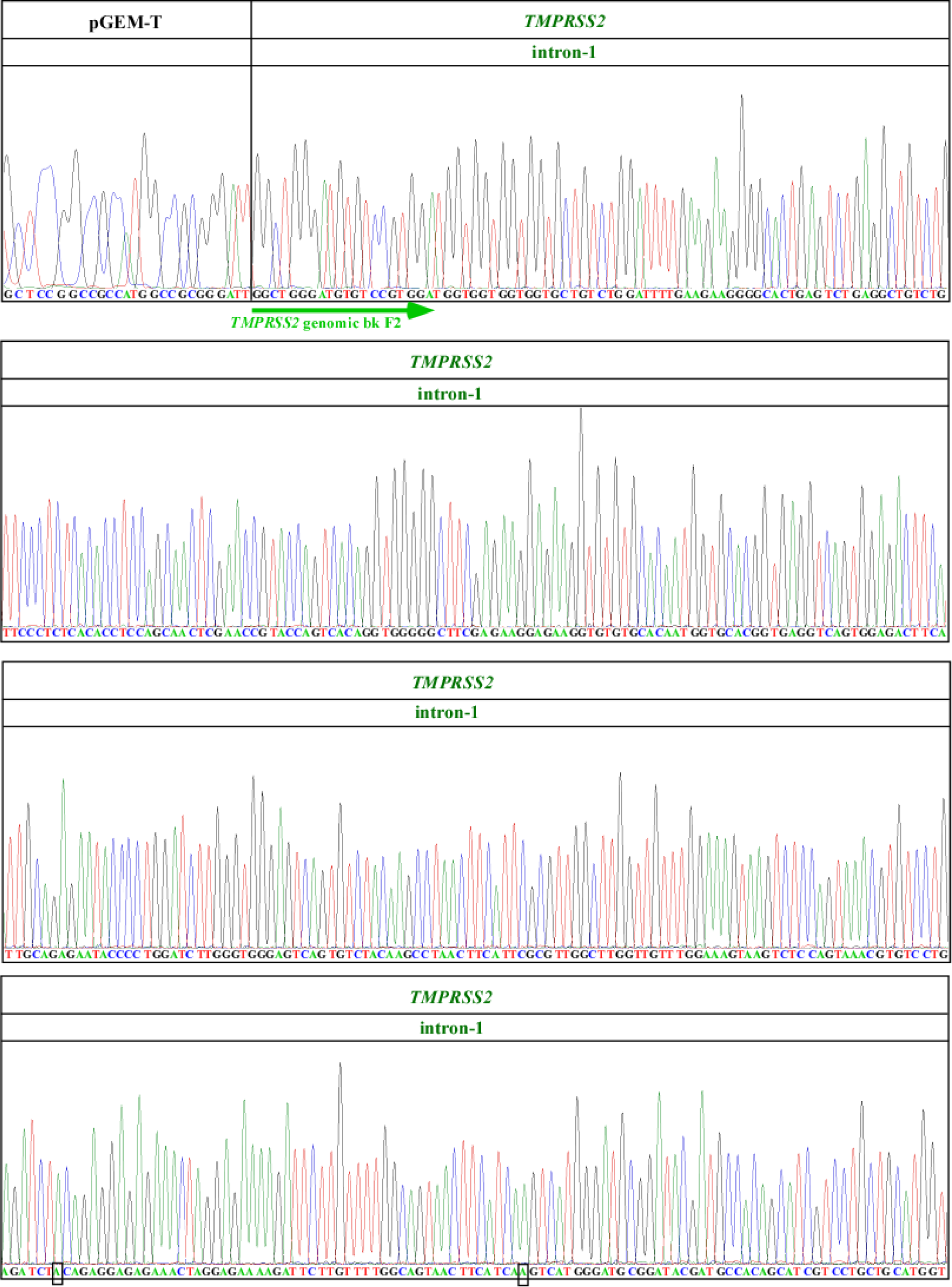
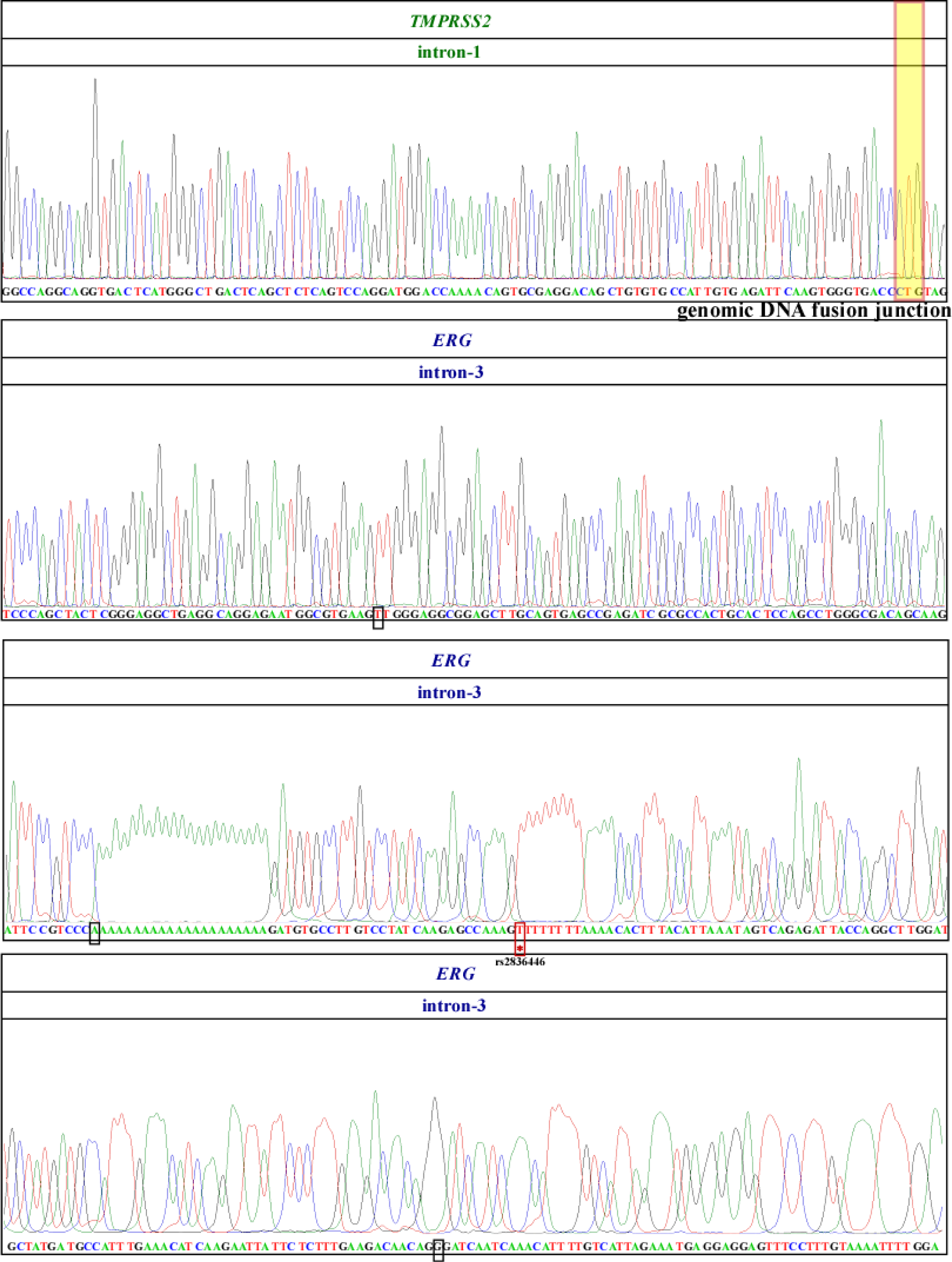
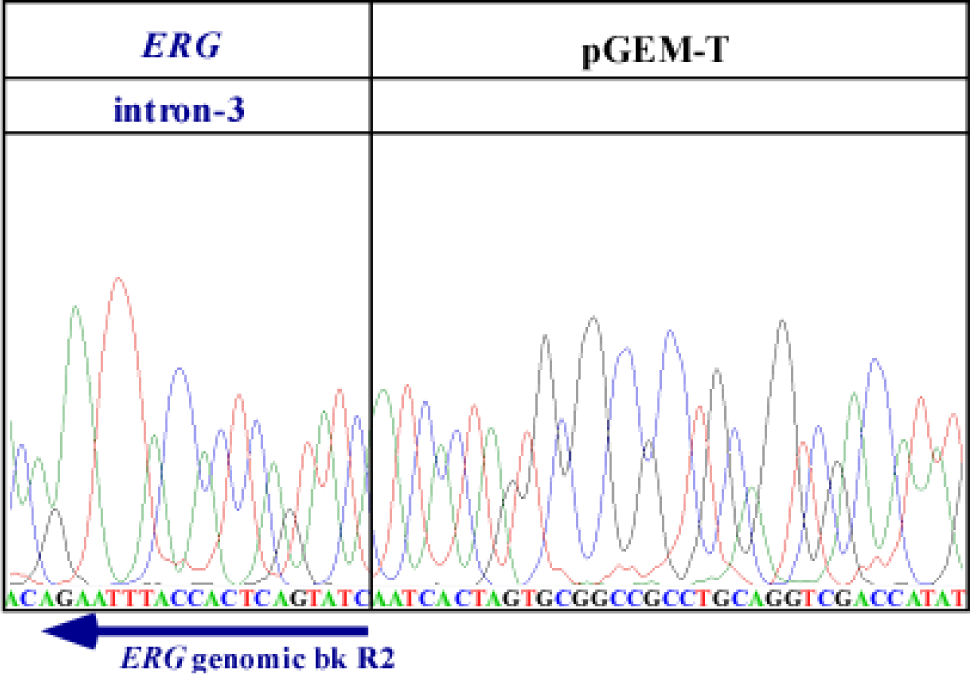
Sanger sequencing confirmed that the induced *TMPRSS2-ERG* fusion is due to gene fusion resulting from genomic arrangement. LNCaP cells transfected with antisense-5 were enriched for the *TMPRSS2-ERG* fusion transcript as described in Fig. S14. Genomic DNA PCR was performed on the enriched LNCaP population which yielded a gene fusion band of ~862 bp as shown in Fig. 3D in the main text. Sanger sequencing chromatograms confirmed that the 862 bp band contains ~500 bp of *TMPRSS2* intron-1 joined to ~362 bp of *ERG* intron-3. The locations of the primers used to amplify the PCR product are shown as the green and blue arrows for *TMPRSS2* intron-1 and *ERG* intron-3, respectively under the chromatograms. The genomic breakpoint between *TMPRSS2* intron-1 and *ERG* intron-3 is indicated by the yellow shaded region that contains a three nucleotide ‘CTG’ microhomology. The pGEM-T plasmid was used for cloning the genomic DNA PCR product. Five mutations shown in black boxes are PCR artifacts because they are absent in other clones that were sequenced in parallel. The mutation denoted by the asterisk (*) in the red box is present in all sequenced clones and is a known Single Nucleotide Polymorphism (rs2836446).

**Fig. S17.**
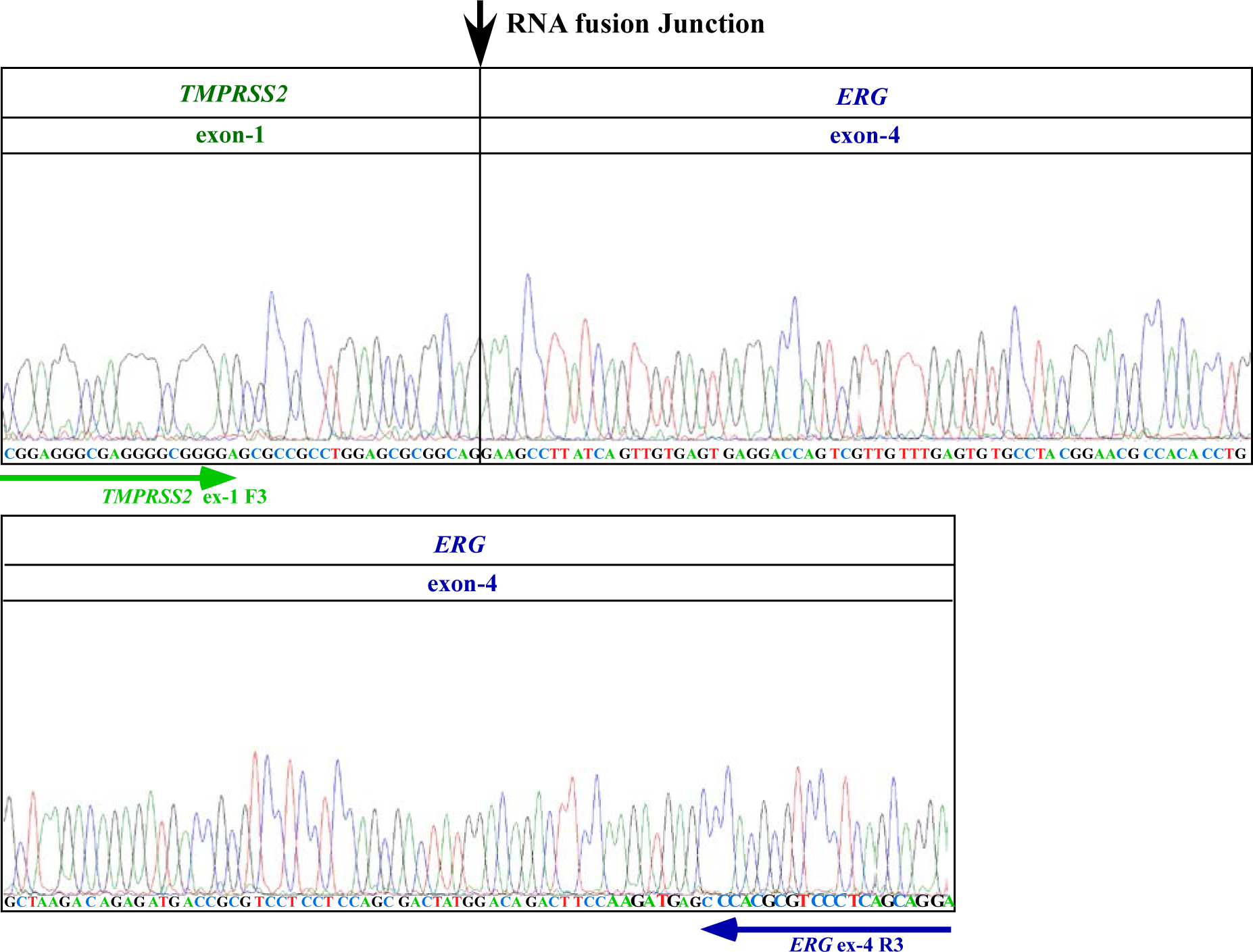
Sanger sequencing confirmation of the induced *TMPRSS2-ERG* fusion transcript in PNT1A cells mediated by antisense-5 RNA. Sanger sequencing confirmed that the induced fusion RNA in normal prostate epithelium cells (as shown in Fig. 3F in the main text) is the same fusion transcript containing *TMPRSS2* exon-1 fused to *ERG* exon-4 as would be expected of mature endogenous *TMPRSS2-ERG* fusion mRNA. This indicates that induction of the *TMPRSS2-ERG* fusion transcript by input RNA is permissible in normal prostate epithelial cells prior to malignant transformation. The locations of the primers used to amplify the PCR product are shown as the green and blue arrows for *TMPRSS2* exon-1 and *ERG* exon-4, respectively. The RNA fusion junction between *TMPRSS2* exon-1 and *ERG* exon-4 is indicated by the black arrow.

**Fig. S18.**
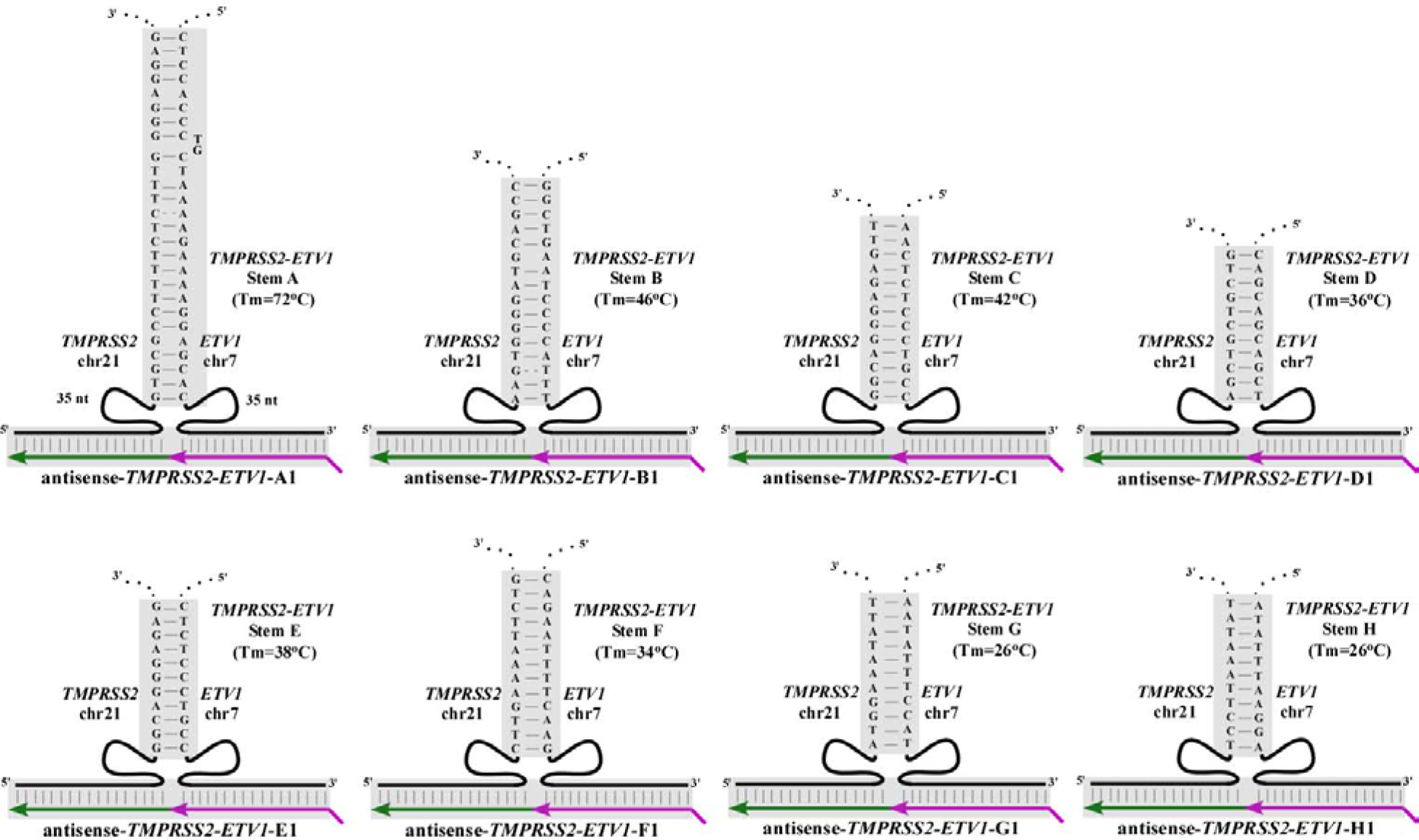
Schematics of the putative three-way junctions formed between *TMPRSS2* and *ETV1* genomic DNA and the corresponding antisense input RNA. We used BLAST alignment to identify eight stems that could be formed with varying degrees of stability by the sense genomic *TMPRSS2* sequence paired with the sense genomic *ETV1* sequence. Matching antisense RNAs were then designed to forge the three-way junction with these putative intron stems. Shaded regions indicate base pairing between antisense input RNA (in green and purple) and genomic DNA (in black). *TMPRSS2* is on chromosome 21 whereas *ETV1* is on chromosome 7. For both, the genomic minus strand is the sense strand.

**Fig. S19.**
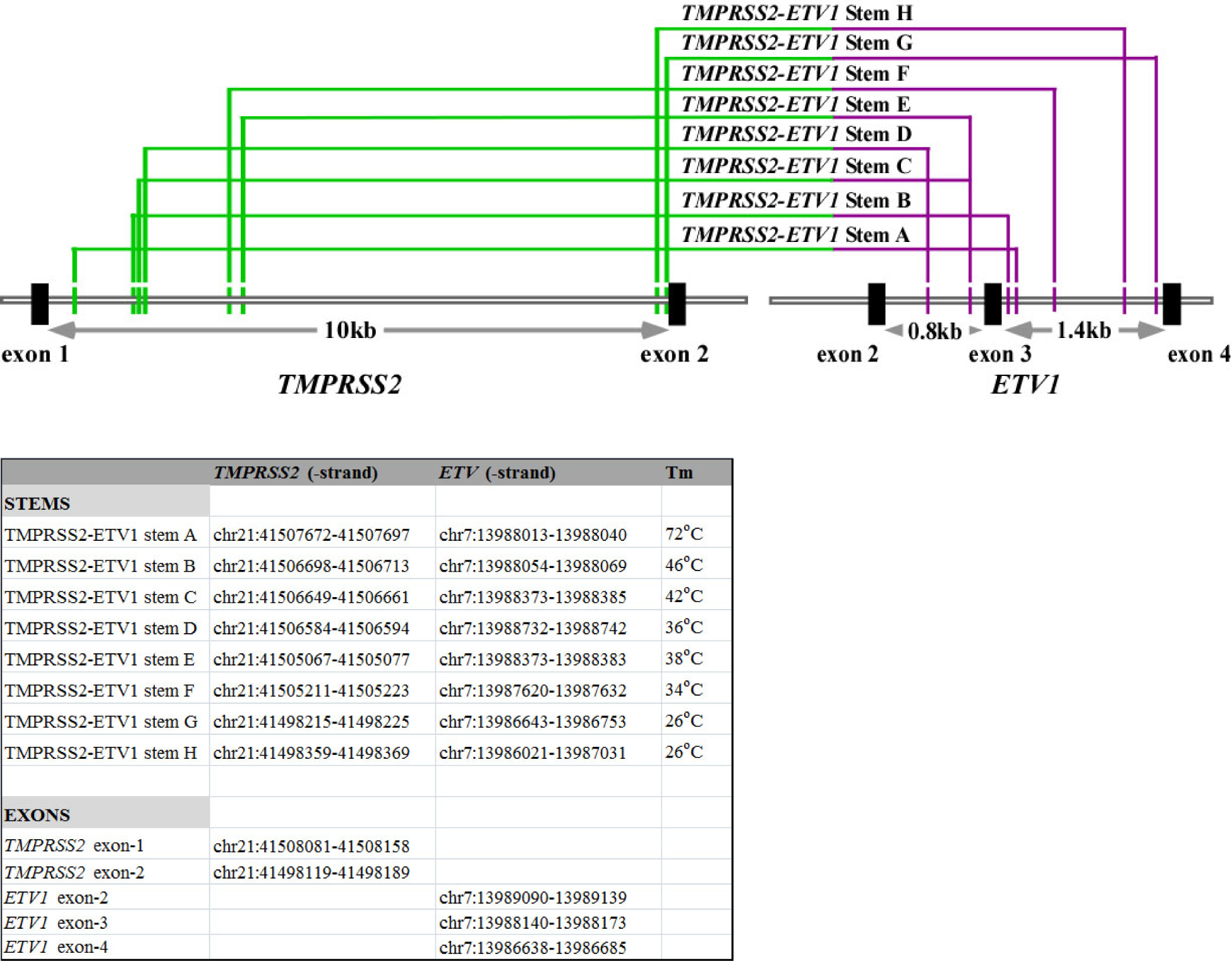
Genome coordinates and Tm for the putative *TMPRSS2-ETV1* genomic stems described in S18. Upper panel: Locations of putative *TMPRSS2-ETV1* stems A, B, C, D, E, F, G, and H identified by BLAST alignment. All stem sequences are located in *TMPRSS2* intron-1 (chromosome 21) paired with sequences in *ETV1* intron-2 or -3 (chromosome 7). Genome coordinates are based on GRCh38/hg38 version. Coordinates of the exons (*TMPRSS2* exons-1 and -2; *ETV1* exons-2, -3, and -4) are also shown.

**Fig. S20.**
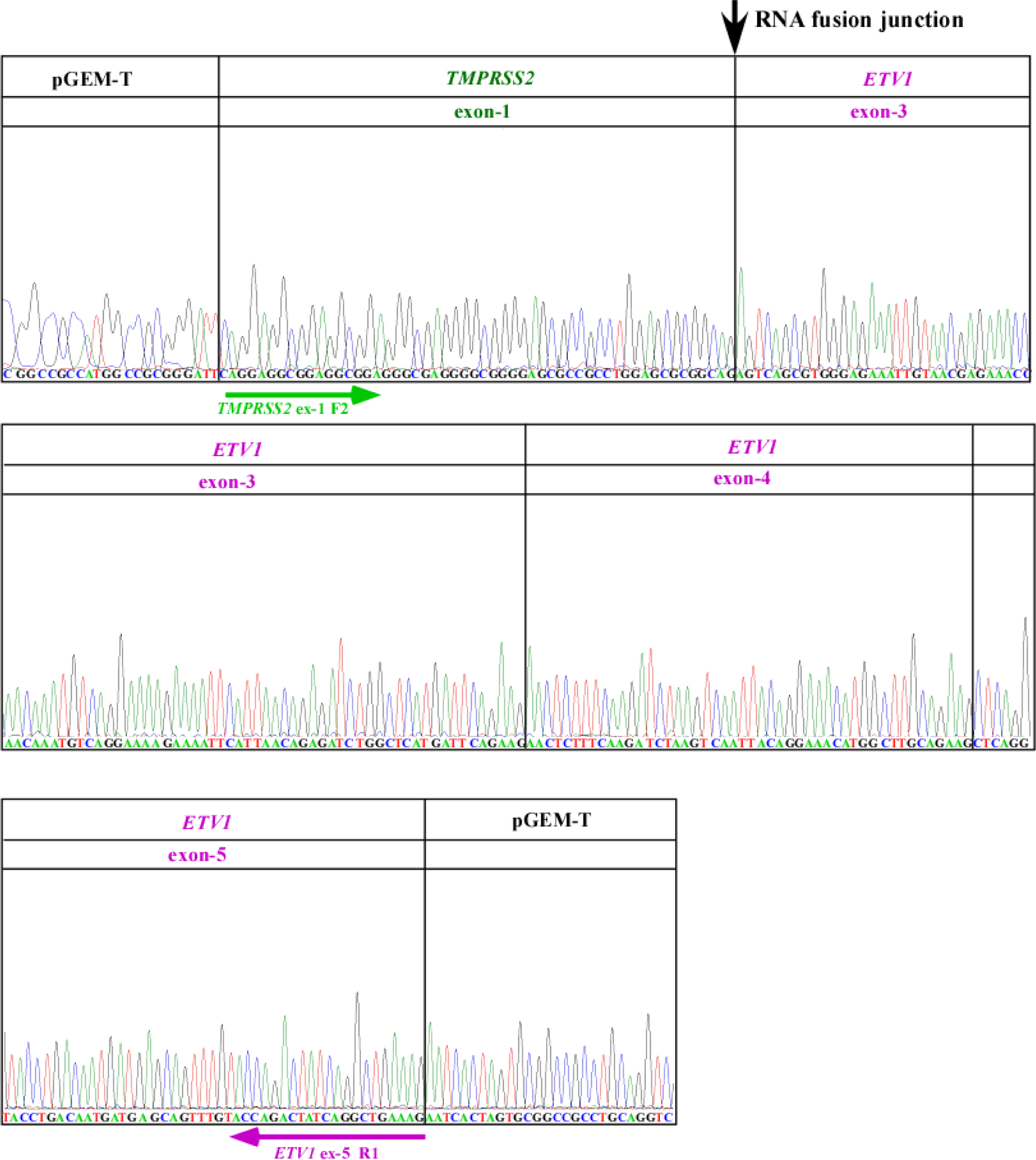
Sanger sequencing confirmed the induced *TMPRSS2-ETV1* fusion transcripts in LNCaP cells. Sanger sequencing chromatograms confirmed that the induced RNA contains *TMPRSS2* exon-1 fused to *ETV1* exon-3 as would be expected of mature endogenous *TMPRSS2-ETV1* fusion mRNA. These results suggest that RNA-mediated gene fusion in mammalian cells is not restricted to *TMPRSS2* and *ERG* but appears to be generally permissible regardless of whether it is an intra-chromosomal (*TMPRSS2-ERG)* or inter-chromosomal (*TMPRSS2-ETV1)* fusion. The induced fusion transcript was converted to cDNA using oligo dT primers and PCR was performed using primers targeting *TMPRSS2* exon-1 and *ETV1* exon-5. The locations of the primers are shown as the green and purple arrow for *TMPRSS2* exon-1 and *ETV1* exon-5, respectively. The RNA fusion junction between *TMPRSS2* exon-1 and *ETV1* exon-3 is indicated by the black arrow.

**Fig. S21.**
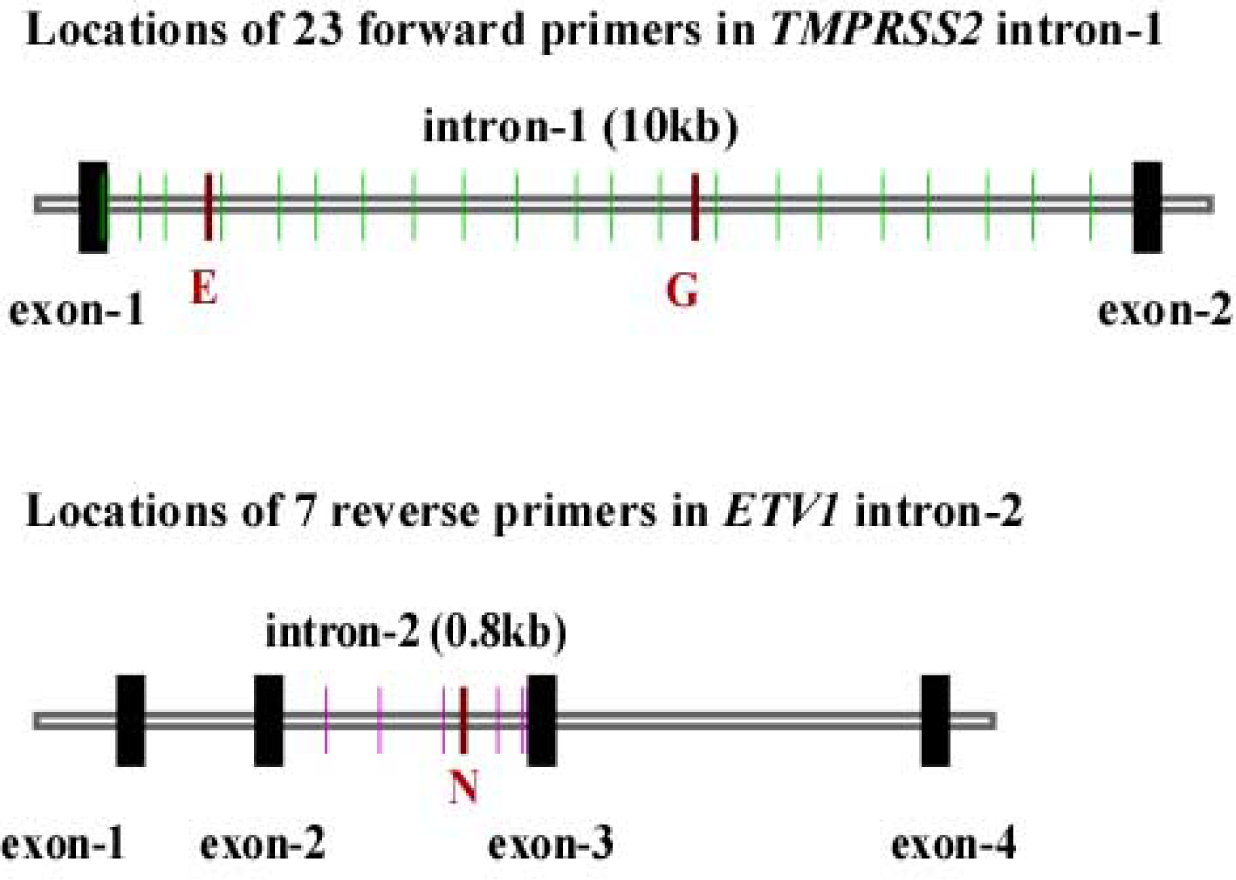
Locations targeted by primers for mapping the *TMPRSS2-ETV1* genomic breakpoints. The locations of 23 forward primers (green) spacing across *TMPRSS2* intron-1 (~10 kb) and 7 reverse primers (purple) spacing across *ETV1* intron-2 (~0.8 kb). Each vertical line represents a target location by a primer or a set of nested primers. The specific primers that amplify the genomic breakpoint shown in Fig. 4E in the main text are labeled as red.

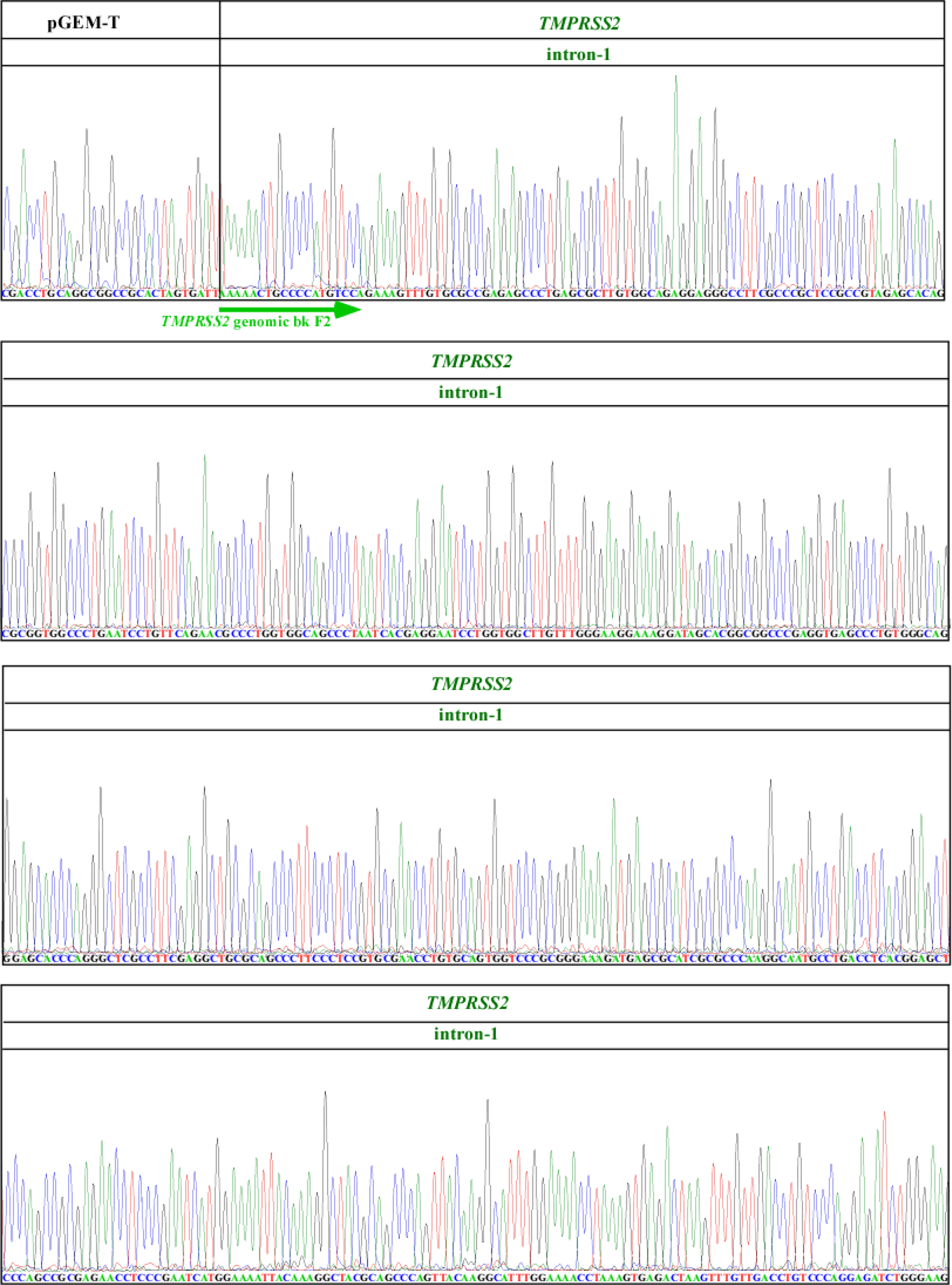

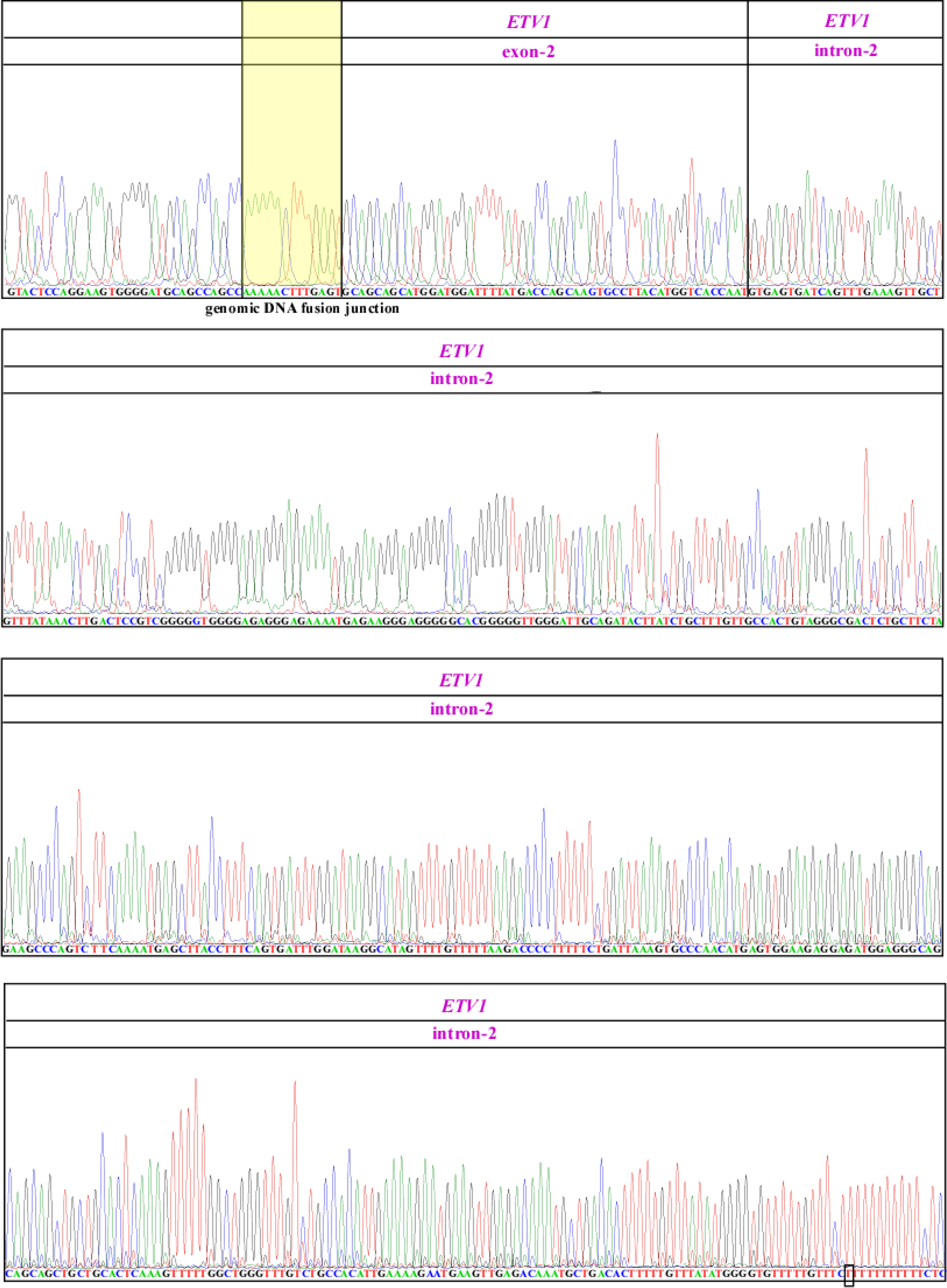

**Fig. S22.**
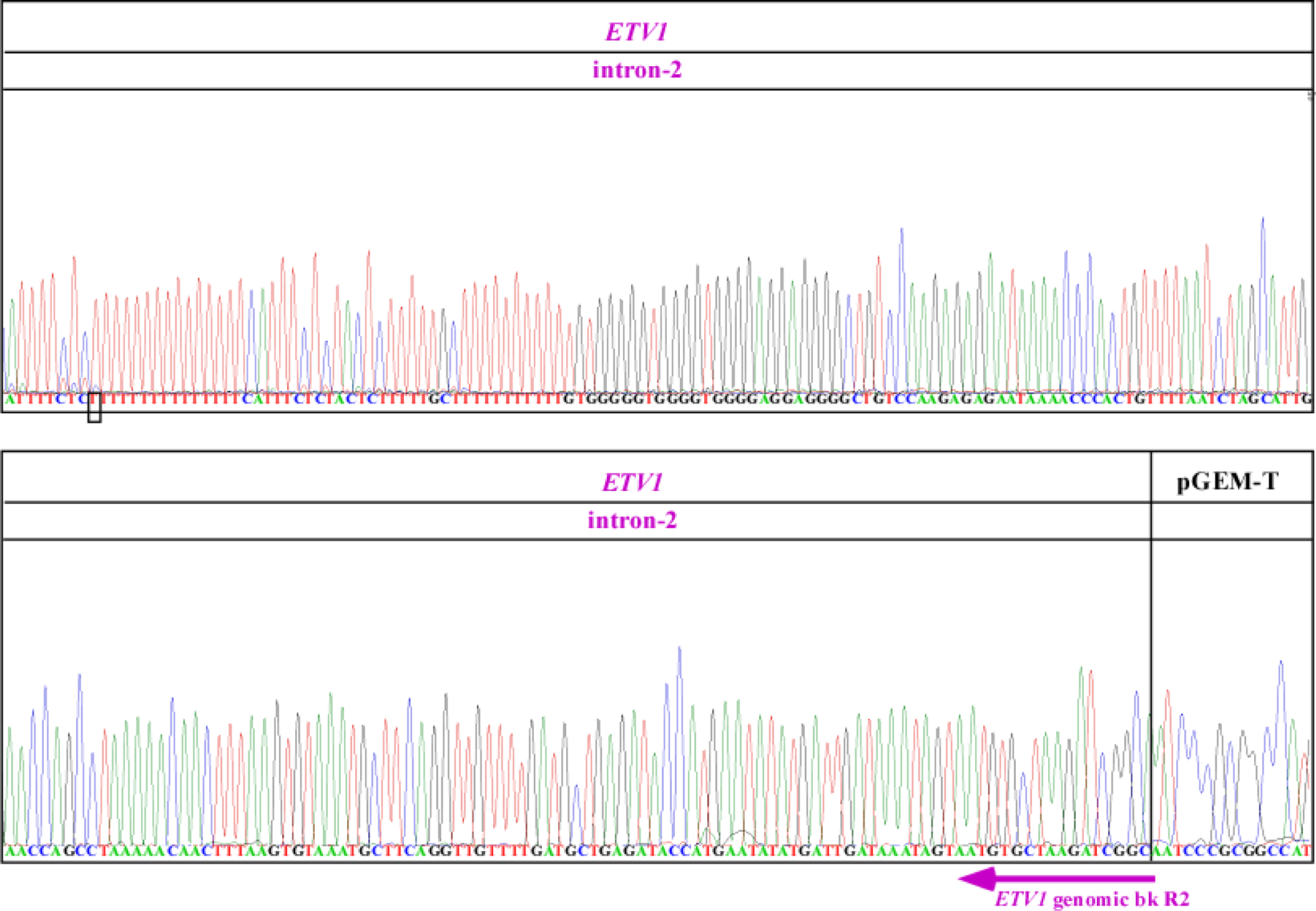
Sanger sequencing confirmed that the induced *TMPRSS2-ETV1* band x is due to gene fusion resulting from genomic arrangement. LNCaP cells transfected with antisense-TMPRSS2-ETV1-A1 were enriched for the *TMPRSS2-ETV1* fusion transcript using a procedure similar to that described in Fig. S14. Genomic DNA PCR was performed on the enriched LNCaP population which yielded three gene fusion bands of ~1150 bp (band x), ~1044 bp (band y), ~1043 bp (band z), as shown in Fig. 4E in the main text. Sanger sequencing chromatograms confirmed that the 1150 bp band x contains ~485 bp of *TMPRSS2* intron-1 joined to ~665 bp of *ETV1* intron-3. The locations of the primers used to amplify the PCR product are shown as the green and purple arrows for *TMPRSS2* intron-1 and *ETV1* intron-3, respectively. Region of microhomology at the breakpoints are boxed by solid lines. The pGEM-T plasmid was used for cloning the genomic DNA PCR product. Mutations shown in black boxes are PCR artifacts because they are absent in other clones that were sequenced in parallel.

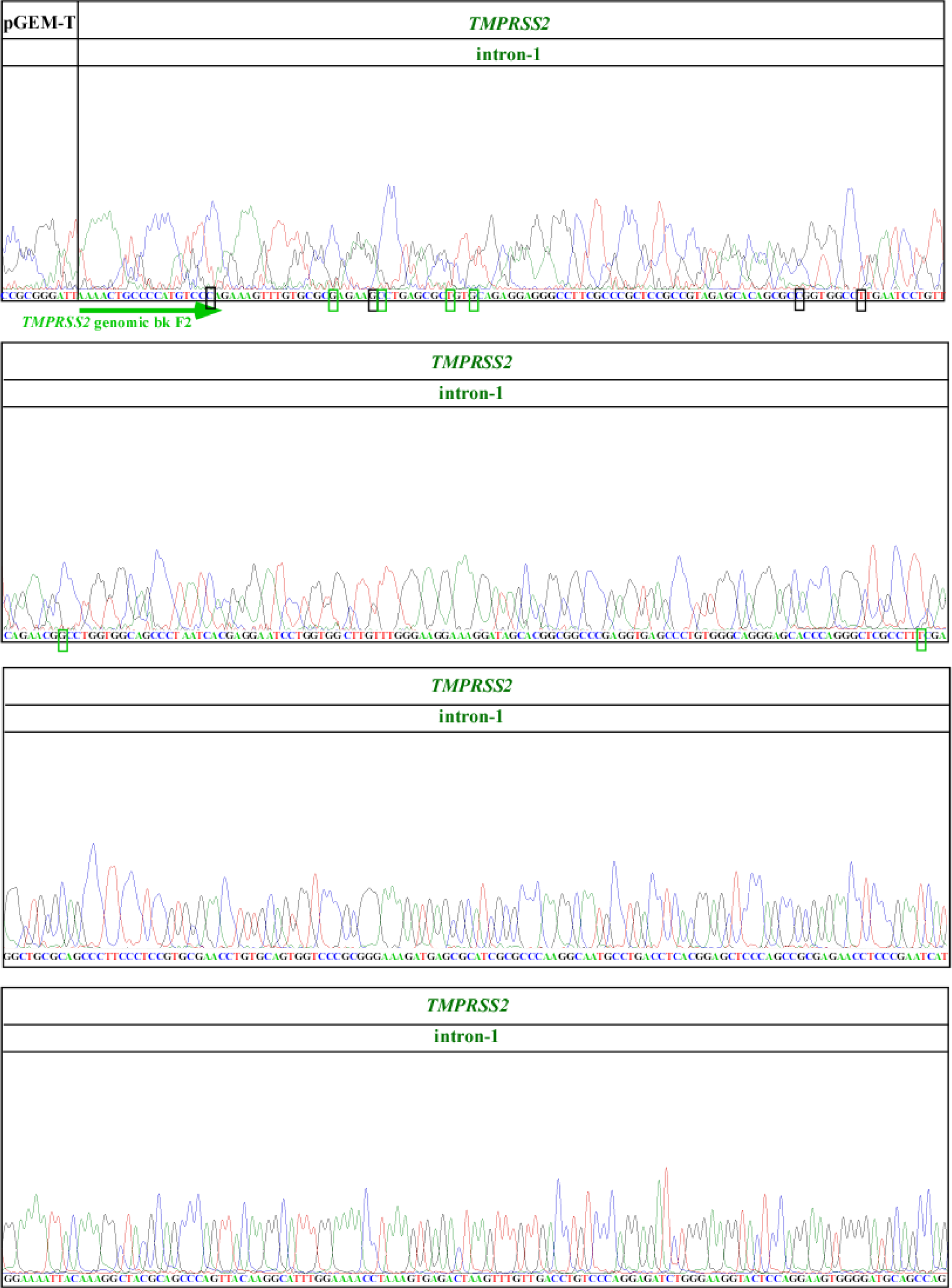

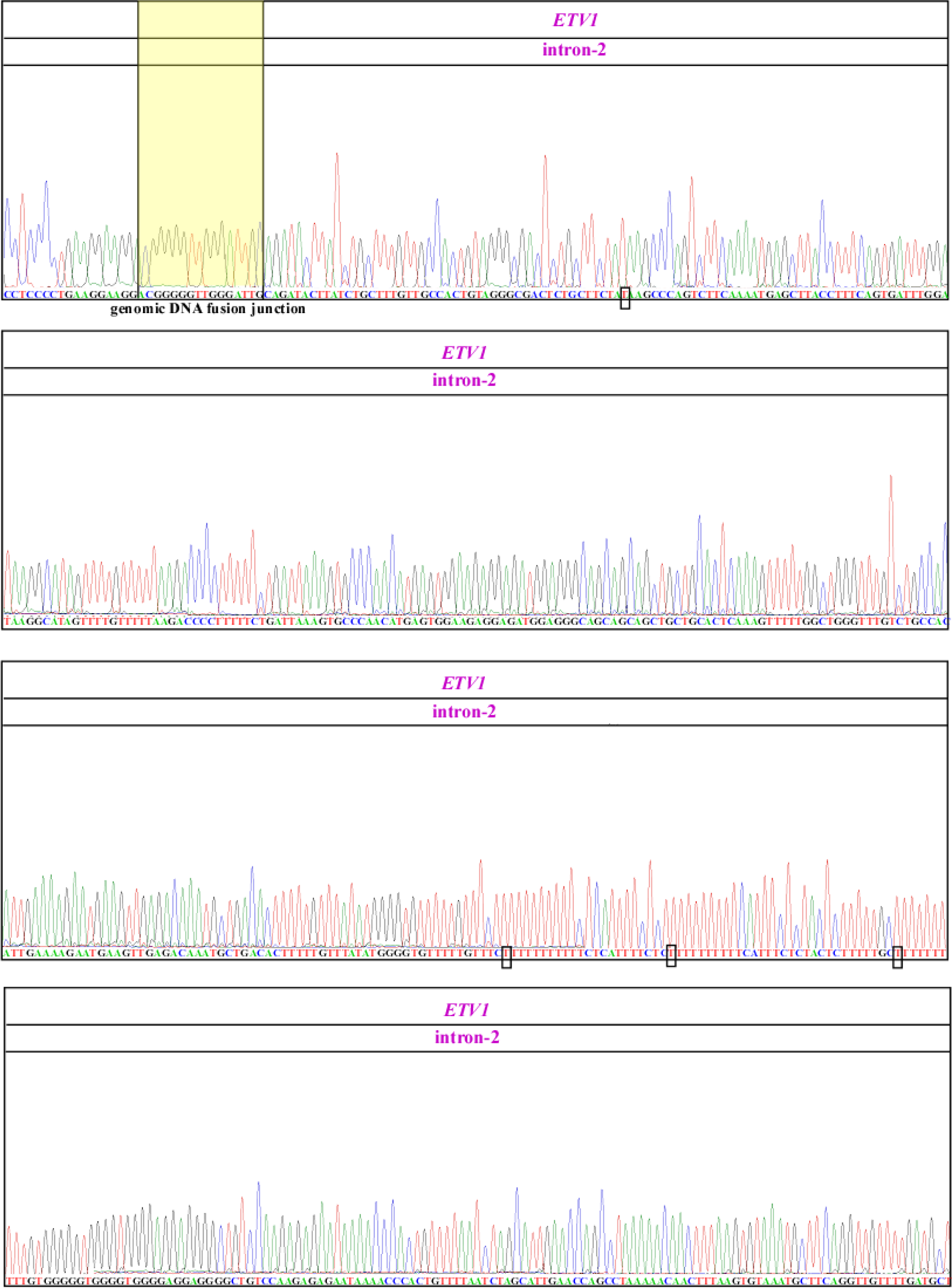

**Fig. S23.**
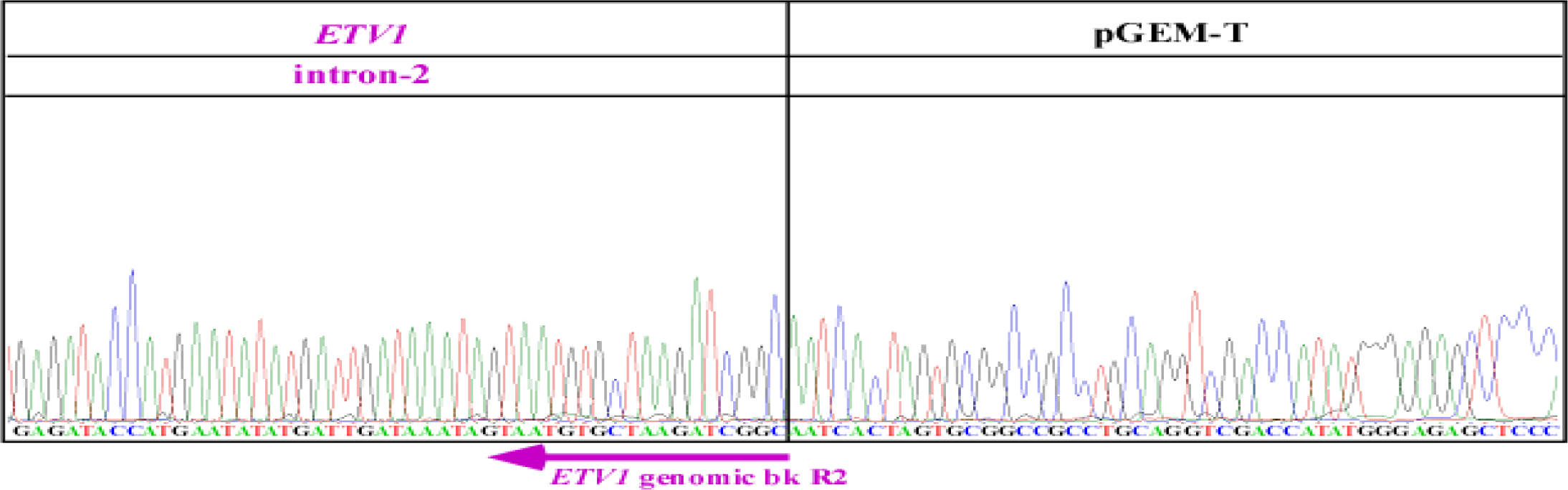
Sanger sequencing confirmed that the induced *TMPRSS2-ETV1* band y is due to gene fusion resulting from genomic arrangement. Similarly, Sanger sequencing was performed on the genomic PCR fusion bands ~1044 bp (band y) shown in Fig. 4E in the main text. The resulting chromatograms confirmed that the 1044 bp band y contains ~516 bp of *TMPRSS2* intron-1 joined to ~528 bp of *ETV1* intron-3. The locations of the primers used to amplify the PCR product are shown as the green and purple arrows for *TMPRSS2* intron-1 and *ETV1* intron-3, respectively. Region of microhomology and indels at the breakpoints are boxed by solid lines. The pGEM-T plasmid was used for cloning the genomic DNA PCR product. Mutations shown in small black boxes are PCR artifacts because they are absent in other clones that were sequenced in parallel, whereas mutations shown in small green boxes are either PCR introduced or not verified because of poor sequencing quality in other clones that were sequenced in parallel.

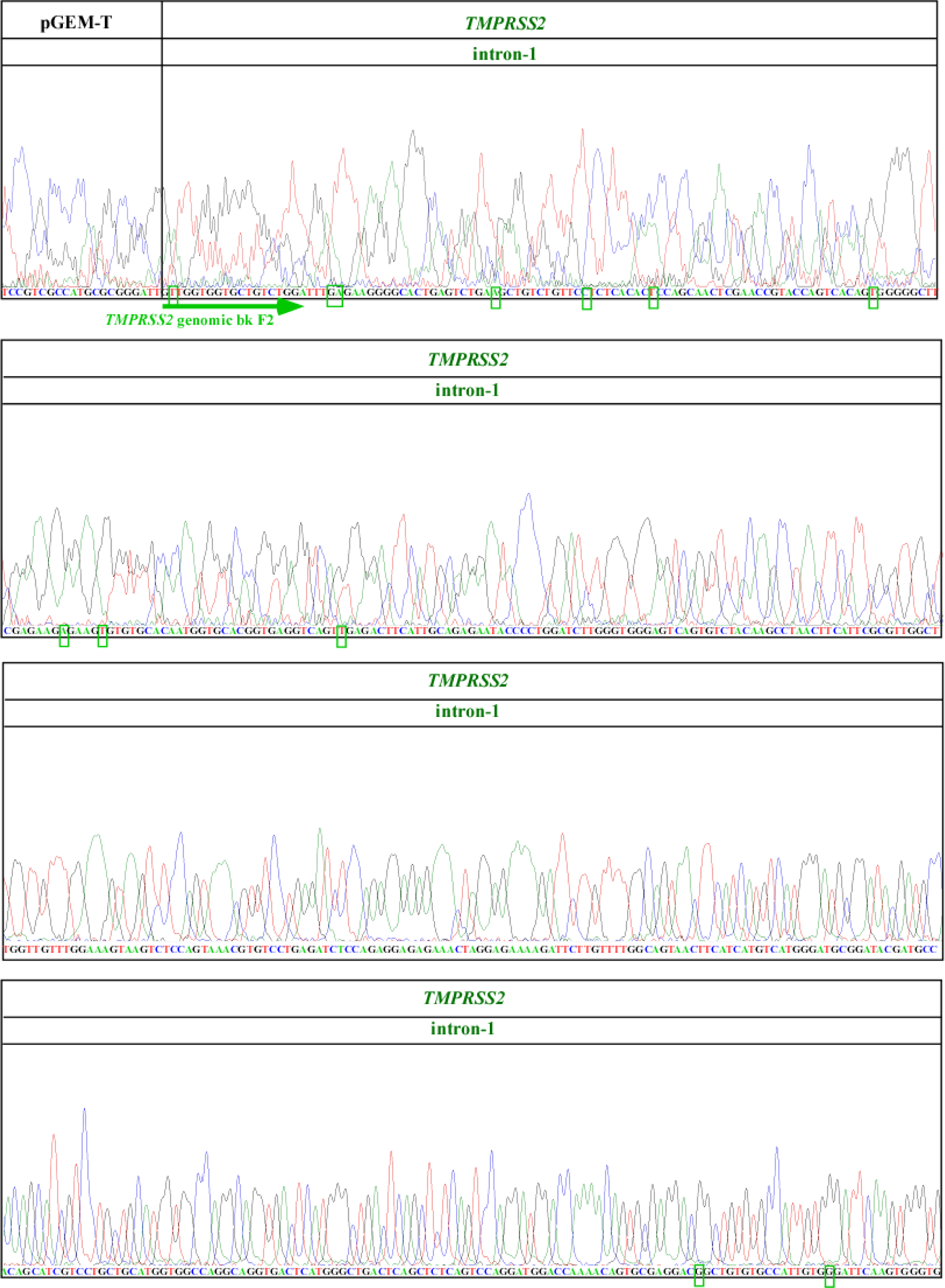

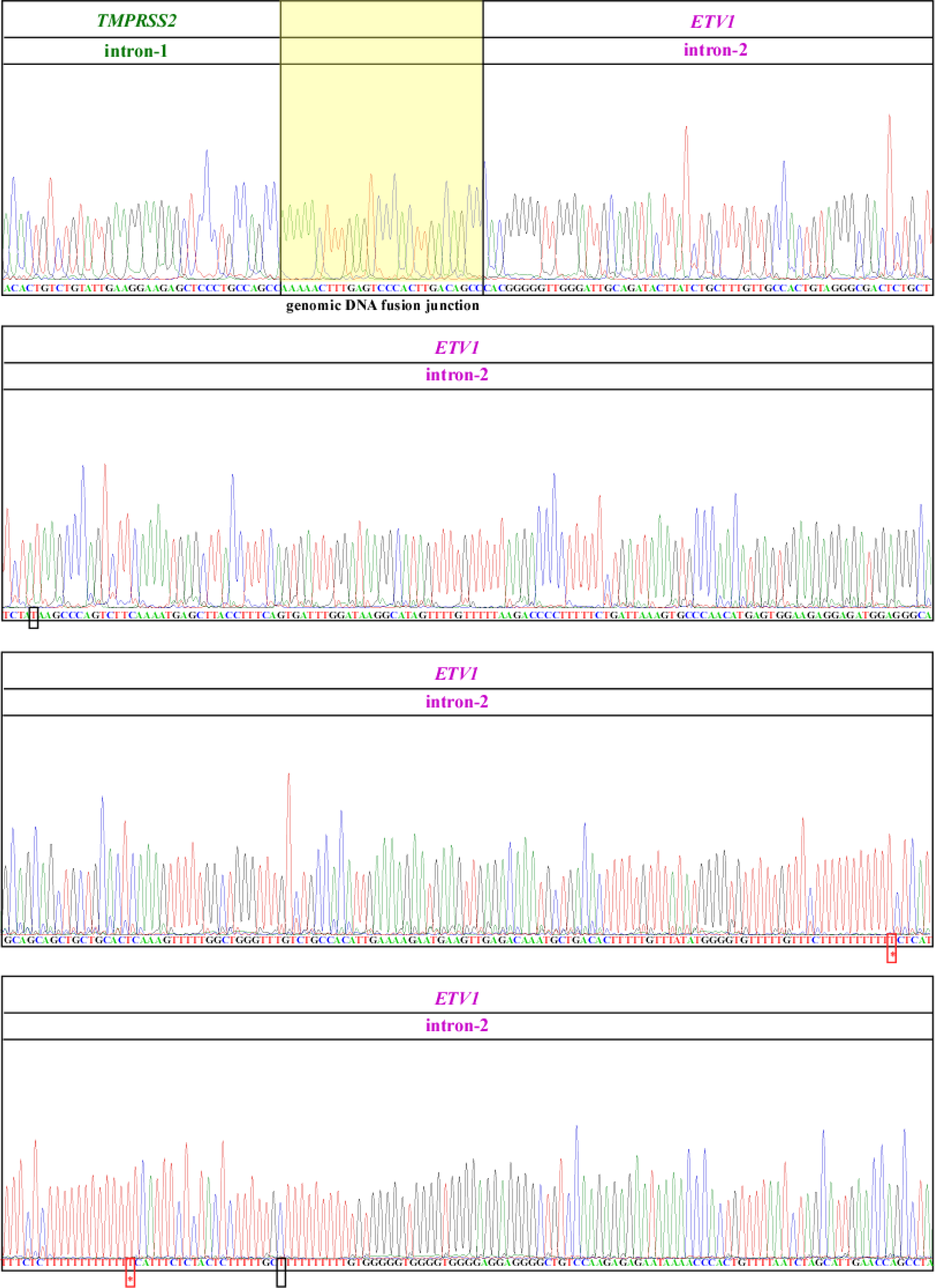

**Fig. S24.**
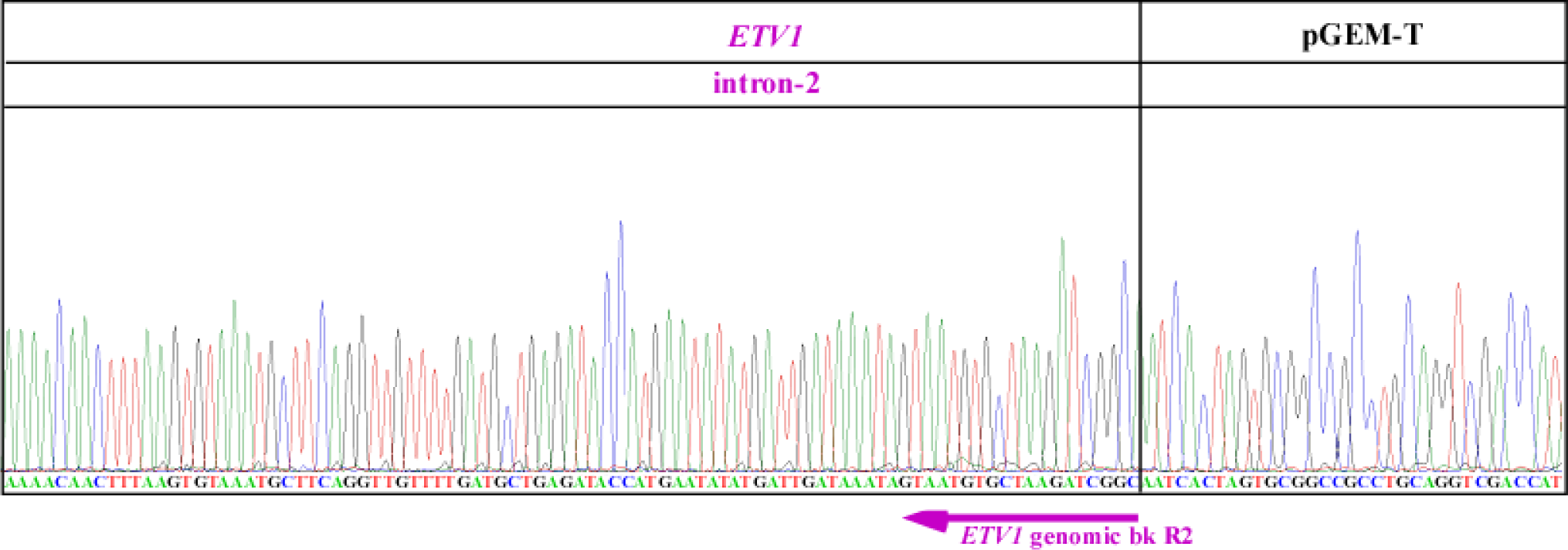
Sanger sequencing confirmed that the induced *TMPRSS2-ETV1* band z is due to gene fusion resulting from genomic arrangement. Similarly, Sanger sequencing was performed on the genomic PCR fusion bands ~1043 bp (band z) shown in Fig. 4E in the main text. The resulting chromatograms confirmed that the 1043 bp band z contains ~514 bp of *TMPRSS2* intron-1 joined to ~529 bp of *ETV1* intron-3. The locations of the primers used to amplify the PCR product are shown as the green and purple arrows for *TMPRSS2* intron-1 and *ETV1* intron-3, respectively. Region of microhomology and indels at the breakpoints are boxed by solid lines. The pGEM-T plasmid was used for cloning the genomic DNA PCR product. Mutations shown in small black boxes are PCR artifacts because they are absent in other clones that were sequenced in parallel, whereas mutations shown in small green boxes are either PCR introduced or not verified because of poor sequencing quality in other clones that were sequenced in parallel.

**Movies S25a & b. Examples of *TMPRSS2-ETV1* fusion gene identified by 3D-microscopy FISH image reconstruction.**

The enriched *TMPRSS2-ETV1* LNCaP cell population was fixed with formaldehyde followed by hybridization with FISH probes against *TMPRSS2* gene (red) and *ETV1* gene (green). Images were acquired on a GE Healthcare DeltaVision image restoration microscope with an Olympus UPLS Apo 100x/1.4 NA and 2k x 2k EDGE/sCMOS camera. Z stacks were acquired with spacing of 0.25 um. Images were deconvolved using conservative algorithm in Resolve3D SoftWorx (version 6.5.2). Max projection and volume visualization was also performed with the same software. S25a represents 3-D along x-axis and S25b along y-axis.

## Materials and Methods

### LNCaP cell culture

LNCaP cells were routinely cultured in RPMI 1640 medium (RPM1 1640, 1X, with L-glutamine, #10-040-CV, CORNING cellgro) containing 10% fetal bovine serum (premium grade FBS, #1500-500, Seradigm) and 1% penicillin/streptomycin (#15140-122, Gibco) in a 5% CO_2_ humidified incubator. For experiments involving the induction of fusion gene by input RNA, regular fetal bovine serum in the culture medium was replaced by Charcoal:Dextran stripped fetal bovine serum (catalog#100-119, Gemini Bioproducts) to remove hormones present in serum. LNCaP cells were cultured in this special medium for 24 hrs prior to plasmid transfection.

### PNT1A cell culture

PNT1A cells were routinely cultured in RPMI 1640 medium containing 10% fetal bovine serum (premium grade FBS, #1500-500, Seradigm) and 1% penicillin/streptomycin (#15140-122, Gibco) in a 5% CO_2_ humidified incubator.

### Transient transfection of plasmids for expressing the input RNAs

Twenty hours prior to transfection, LNCaP cells were seeded in 12-wells plate (BioLite 12 Well Multidish, #130185, Thermo Fisher Scientific) with a density of 5x10^5^cells/well and 1 ml/well of culture medium containing Charcoal:Dextran stripped fetal bovine as described above. Transfection was performed using Turbofect transfection reagent (Thermo Scientific, #R0531) according to manufacturer’s protocol. Briefly, 1μg of a particular plasmid was first diluted in 100μl of the serum-free DMEM followed by immediate mixing by pipetting. 4μl of the transfection reagent was then added to the diluted DNA followed by mixing and incubation for 20 min. The DNA/transfection reagent mixture was then added drop wise to a well containing LNCaP cells in 1ml medium.

For transfection in PNT1A cells, 5x10^5^ cells/well were plated in 12-wells plate in 1 ml/well of cultured medium 24 hrs prior to transfection. Transfection was performed using the same formula described for LNCaP cells. For repetitive transfections, initially transfected PNT1A cell population were split every three days, half was processed for RT-PCR assay and half was seeded again in a new well for the next transfection.

### DHT preparation and treatment

DHT (Dihydrotestosterone) was purchased from Sigma Adlrich (5α-Androstan-17β-ol-3-one, #A8380). Concentrated stock of 1500μM was prepared by dissolving 4.3566 mg of DHT powder in 10 ml of 100% ethanol (200 proof ethanol, Koptec, #V1016) and then aliquoted in 1ml tubes and stored at -80°C.

For treating cultured cells, concentrated DHT stock was diluted as 10x working solutions (for example, for 0.9 μM final concentration, 10x is prepared as 9.0 μM) with the appropriate complete culture medium and used immediately. Complete media for LNCaP cells: RPMI 1640 + 10% Charcoal:Dextran stripped fetal bovine serum + 1% penicillin/streptomycin.

Complete media for PNT1A: RPMI 1640 + 10% fetal bovine serum + 1% penicillin/streptomycin. Six hrs post transfection, 111μl of fresh 10x DHT working solutions was added to each well of 12-wells plate containing 1ml medium and transfected cells.

For long-term treatment, medium was changed with fresh DHT every three days.

### RNA isolation

Total RNA from cultured cells was extracted using High Pure RNA isolation Kit according to manufacturer’s instructions (#11828665001, Roche). Briefly, cells were suspended in 200μl of PBS buffer and were then lysed with 400μl of lysis buffer. The sample was then passed through the filter assembly resulting in the binding of the nucleic acids to the filter. The filter containing nucleic acids was then incubated with DNase I dissolved in DNase incubation buffer to degrade genomic and plasmid DNAs. The column was then rinsed with wash buffer and total RNA then eluted in a new tube for further analysis.

For detection of residual genomic and plasmid DNA, eluted RNA was subject to PCR reaction with primers specific to intron regions of house-keeping gene GAPDH, and with primers specific to plasmid transfected. Total RNA was converted to cDNA only if it is validated as free of DNA contamination.

### Reverse transcription reaction

1 μg of total RNA was used for each reverse transcription reaction according to manufacturer instruction (superscript III RT, # 18080-051, Invitrogen). RNA was converted to cDNA either with Oligo dT primer (for induced fusion transcripts) or with random hexamers (for input RNAs expressed by U6 promoter). After the addition of dNTPs, the mixture was denatured at 65°C for 5 minutes. This was followed by the addition of a master-mix containing 1× superscript buffer, 10 mM DTT, 5 mM Magnesium chloride, RNaseOUT and Superscript III reverse transcriptase. Reactions were carried out at 50°C for 50 minutes and then terminated by incubation at 85°C for 5 minutes. cDNA was then treated with RNase-H for 20 minutes at 37°C to degrade RNA in DNA/RNA hybrid. 1 μl of cDNA was used as template for each subsequent PCR reaction.

### RT-PCR for detecting induced fusion transcripts

The majority of induced fusion RNAs in this manuscript were detected using one-round RT-PCR. The following cases were assayed using three-round nested PCR: (1) the results of DHT treatment at physiological concentrations as shown in Fig. 1D and 6A, (2) the induction of *TMPRSS2-ERG* fusion transcript in non-malignant PNT1A cells as shown in Fig. 3F, (3) the specificity of input RNAs assayed in Fig. 4C, and (4) endogenous *ERG* level detection in LNCaP cells in Fig. S13. The following cases were assayed using two-round nested PCR: the induction of *TMPRSS2-ETV1* fusion RNA as shown in Fig. 4A and 4B, and the detection of *TMPRSS2-ETV1* fusion in the enriched population in Fig. 4D.

PCR was done with a standard three-step protocol using REDTaq DNA polymerase (#D5684-1KU, Sigma) according to manufacturer instruction.

Reaction was set as follows:

<u>PCR reaction</u>:

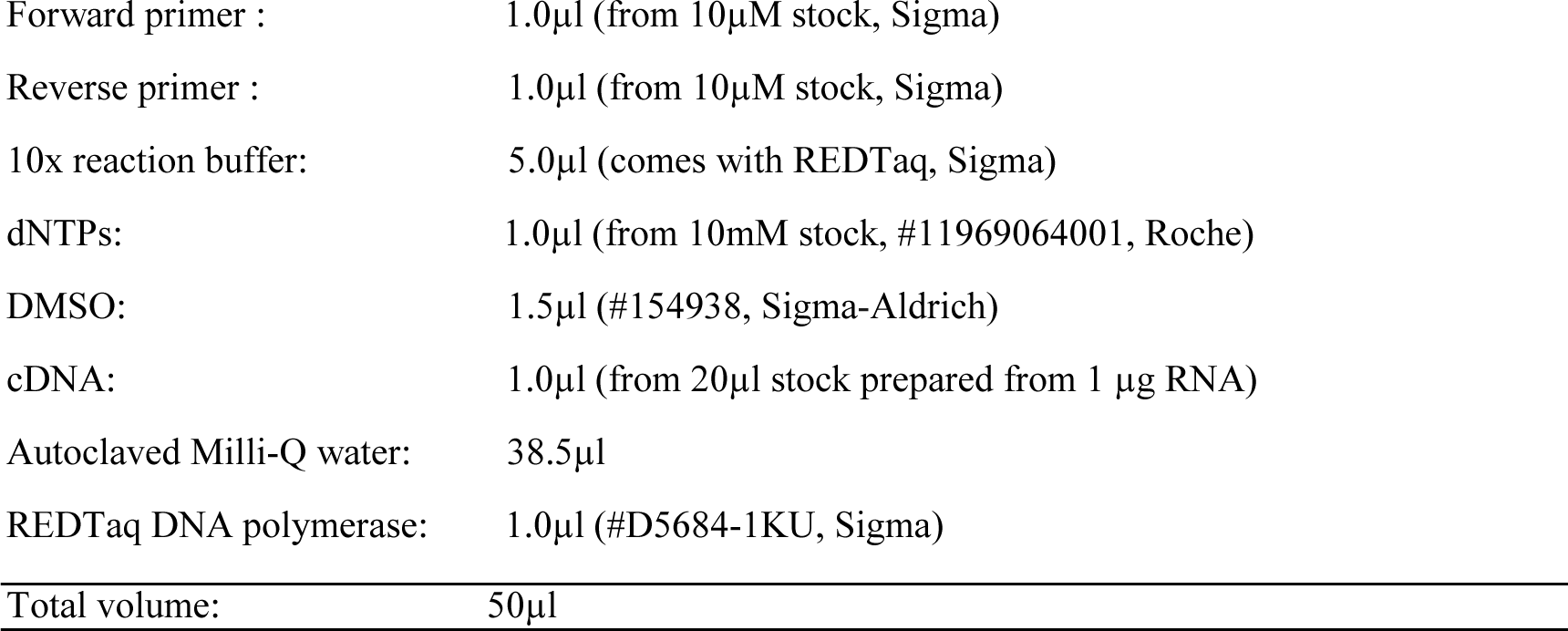

<u>Standard one-round PCR conditions for *TMPRSS2-ERG*</u>:

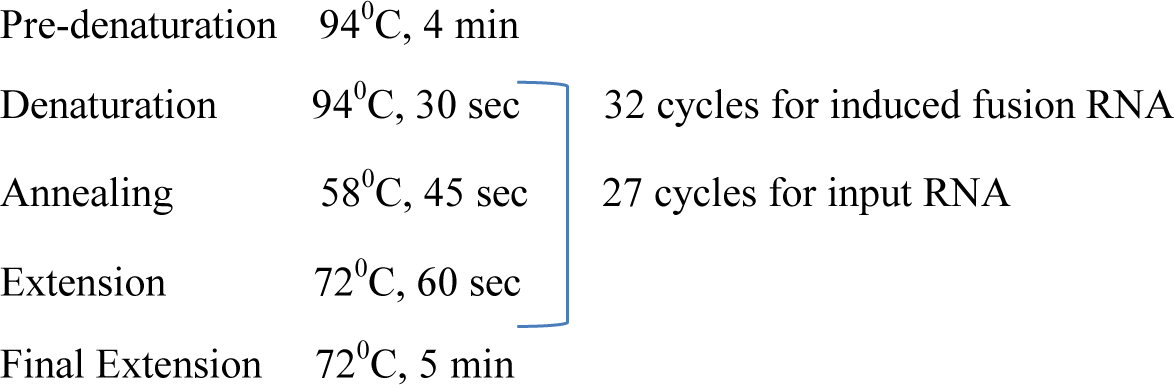

<u>PCR conditions for three-round nested PCR for *TMPRSS2-ERG*:</u>

**<u>1^st^ round</u> :** PCR with *TMPRSS2* ex-1 F1 and *ERG* ex-4 R1 on 1μl of cDNA

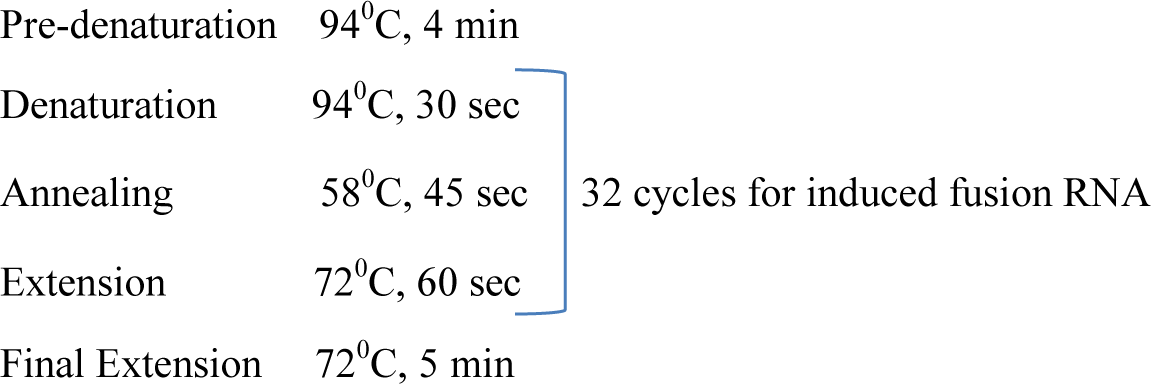

**<u>2^nd^ round</u> :** PCR with *TMPRSS2* ex-1 F2 and *ERG* ex-4 R2 on 1μl of 1^st^ round product, PCR conditions same as 1^st^ round.

**<u>3^rd^ round</u> :** PCR with *TMPRSS2* ex-1 F3 and *ERG* ex-4 R3 on 1μl of 2^nd^ round product, PCR conditions same as 1^st^ round.

<u>PCR conditions for two-round nested PCR for *TMPRSS2-ETV1*:</u>

**<u>1^st^ Round</u>**: Top down PCR with *TMPRSS2* ex-1 F1 and *ETV1* ex-6 R1

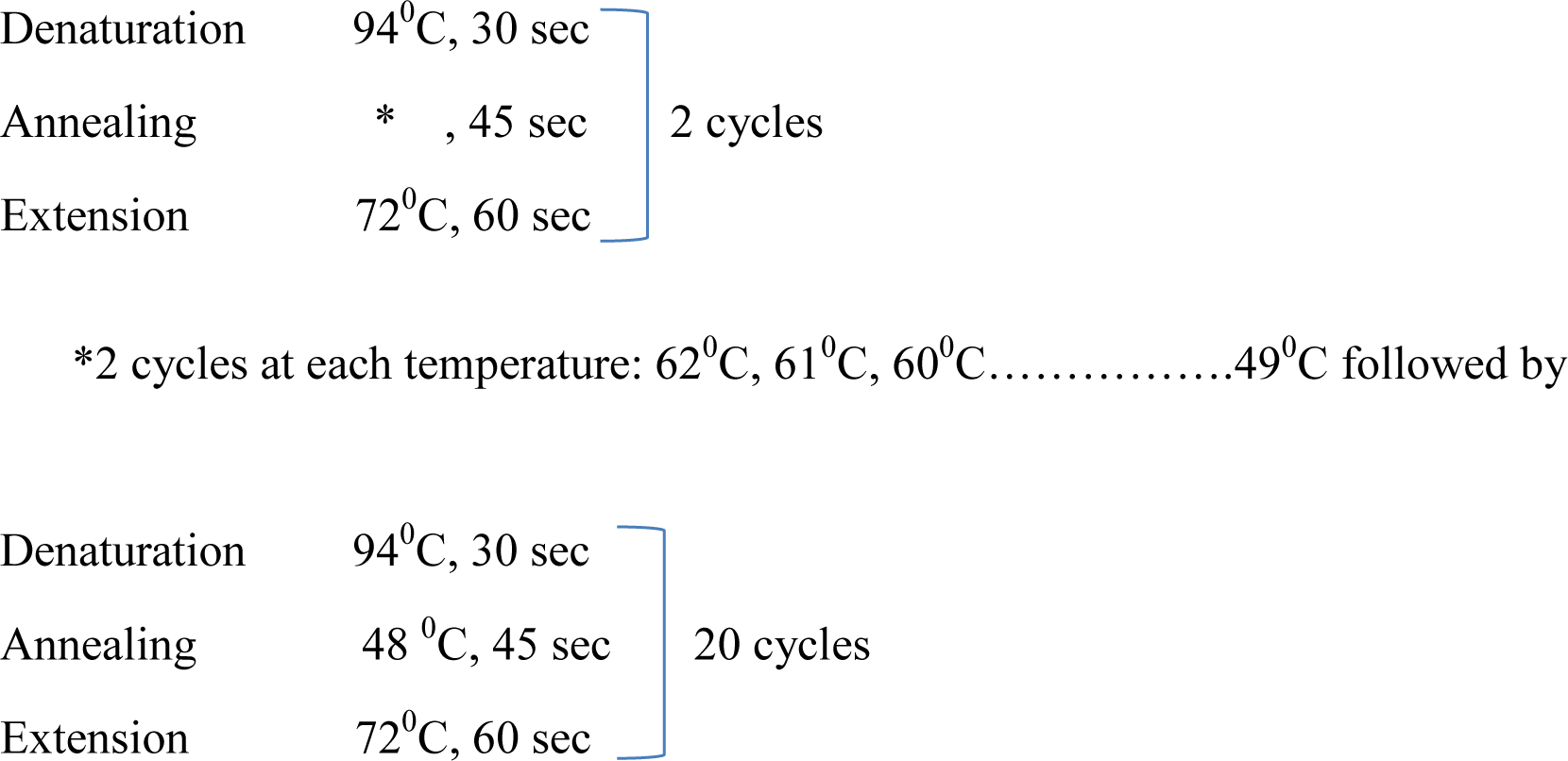

**<u>2^nd^ Round</u>**: PCR with *TMPRSS2* ex-1 F2 and *ETV1* ex-5 R1 on 1μl of 1^st^ round.

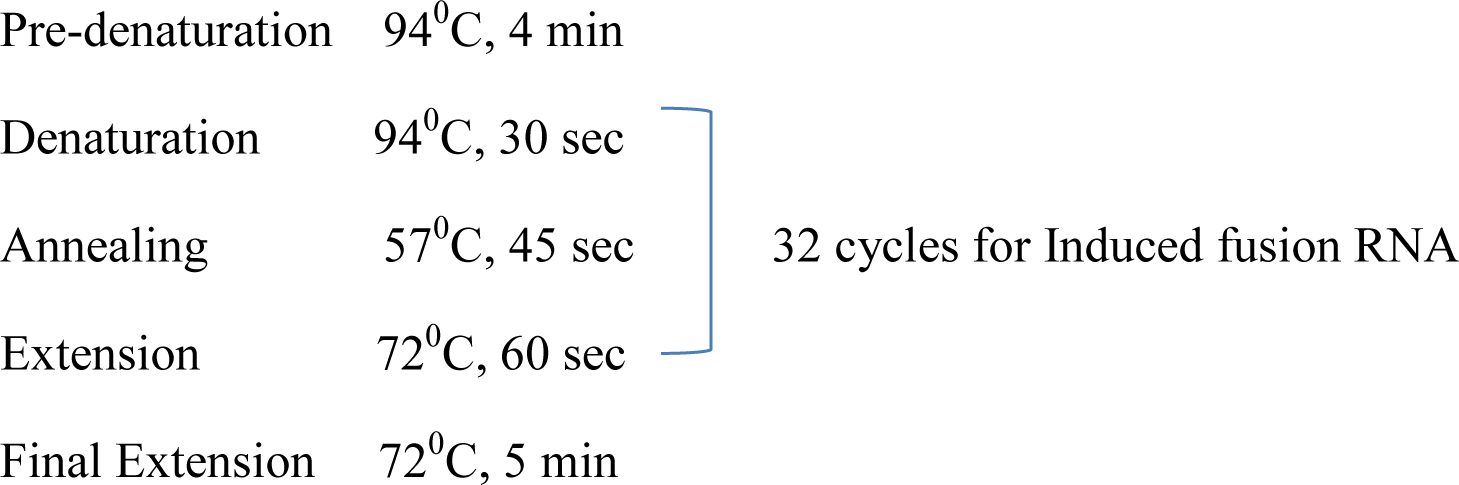

**<u>3^rd^ round</u> (**for Fig. 4D): PCR with *TMPRSS2* ex-1 F3 and *ETV1* ex-5 R2 on 1μl of 2^nd^ round, PCR conditions same as 2^nd^ round.

### Long range PCR for detecting genomic DNA fusion junction

Nested long-range PCRs according to the manufacturer’s protocols using LA PCR kit (Takara, # RR002M). 200 ng of genomic DNA was used in each reaction and PCR was performed with annealing and extension at 68°C for 20 minutes. 1μl from the above reaction (1st round PCR) was used as template for the 2nd round PCR.

For the genomic breakpoint identified in this manuscript, 1st round long range PCR was done using primers *TMPRSS2* genomic bk-F1 and *ERG* genomic bk-R1 shown in primer list below. 1μl from the above reaction (1st round PCR) was used as template for the 2nd round PCR using inner primers *TMPRSS2* genomic bk-F2 and *ERG* genomic bk-R2.

### Cloning and Sanger sequencing of induced fusion transcripts

PCR amplified cDNA bands were excised from the gel and eluted using QIAquick Gel Extraction Kit (#28706, Qiagen). The eluted bands were then cloned to pGEM-T vector (pGEM-T vector system I, # A3600) following manufacturer instruction. Sanger sequencing was performed using the service of Beckman Coulter Genomics.

### Tm calculations

Melting temperature (Tm) of putative genomic DNA stems were calculated using the following formula (Rychlik and Rhoads, 1989):

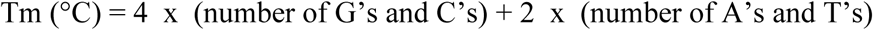

A high energy G·T and A·C wobble pair known to have Watson-Crick like geometry in DNA double helix (Kimsey and Al-Hashimi, 2014; Watson and Crick, 1953) are considered as having the same stability as an A·T pair.

### Fluorescent in situ hybridization (FISH)

Enriched population carrying *TMPRRS2-ETV1* fusion events were first grown on 18 mm round #1 coverglass in a 12-well cell culture plate at the initial density of 200-400k/well. Cells were then fixed with 4% (vol./vol.) formaldehyde followed by denaturation of DNA with 0.1 N HCl for 5 min and with 70% formamide at 85°C for 7 min. Hybridization of target DNA with probes were done at 37°C for 16hr in a humidified chamber. Cells were then washed, stained with DAPI and imaged with microscope. FISH probes for *TMPRSS2* (RP11-35C4, red) and *ETV1* (RP11-769K2, green) were purchased from Empire Genomics.

### α-Amanitin assay

Twenty hours prior to transfection, LNCaP cells were seeded in 12-wells plate (BioLite 12 Well Multidish, #130185, Thermo Fisher Scientific) with a density of 5x10^5^cells/well and transfection was performed using Turbofect transfection reagent (Thermo Scientific, #R0531) as described earlier. DHT was added at the final concentration of 0.9μM six hours post transfection. Following overnight incubation, cells were then treated with 4μg/ml α-amanitin for various time periods (0, 2, 6, 12 and 24 hours). Cells were then revived in fresh medium containing 0.9μM DHT without α-amanitin and RT-PCR was performed for either *TMPRSS2-ERG* or *TMPRSS2-ETV1* fusion.

### Input RNA sequences

#### Sense-1

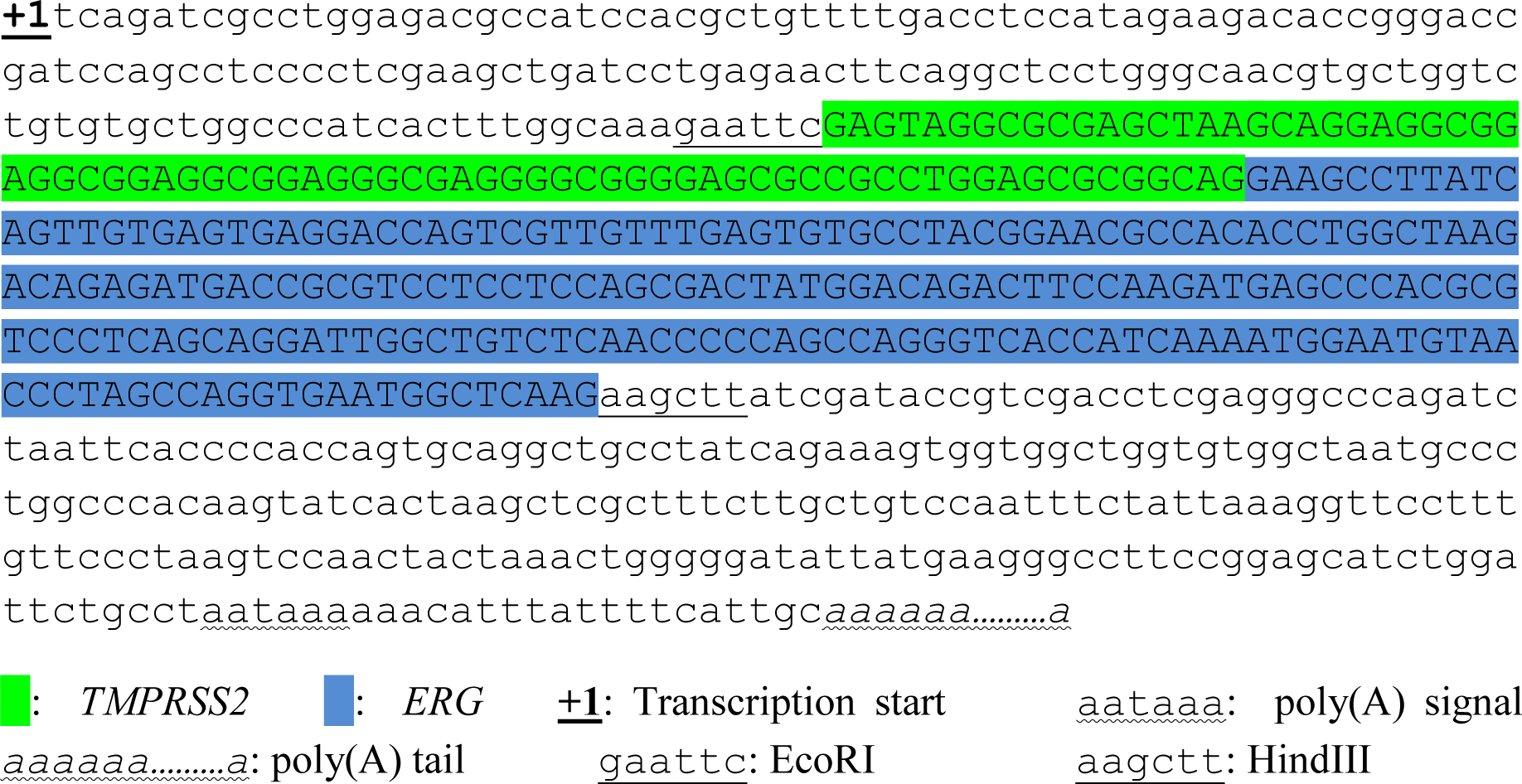

#### Sense-2

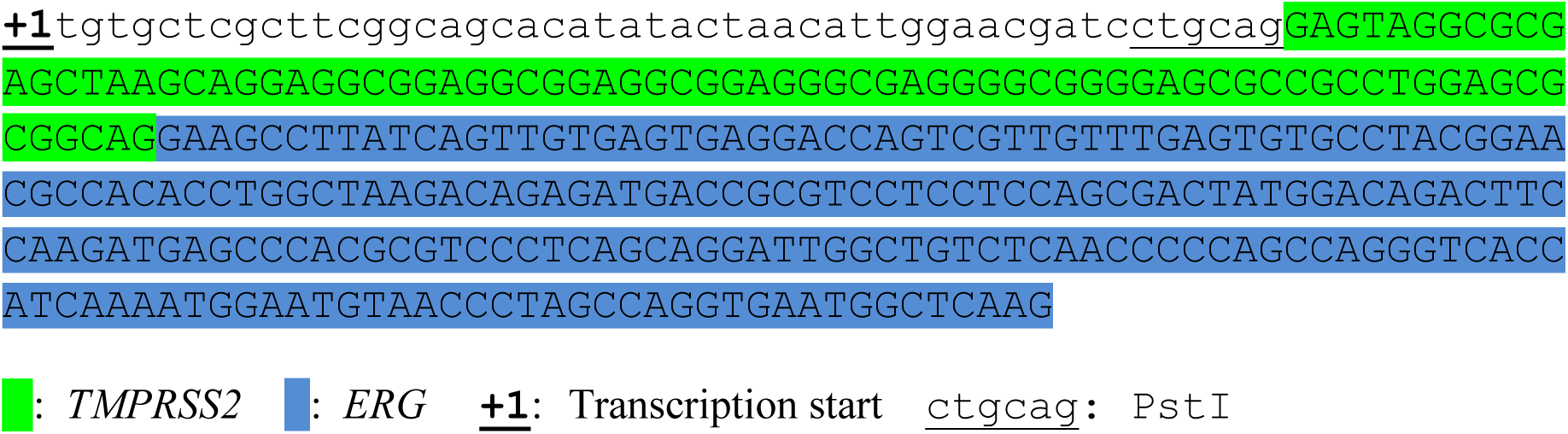

#### Sense-2 long

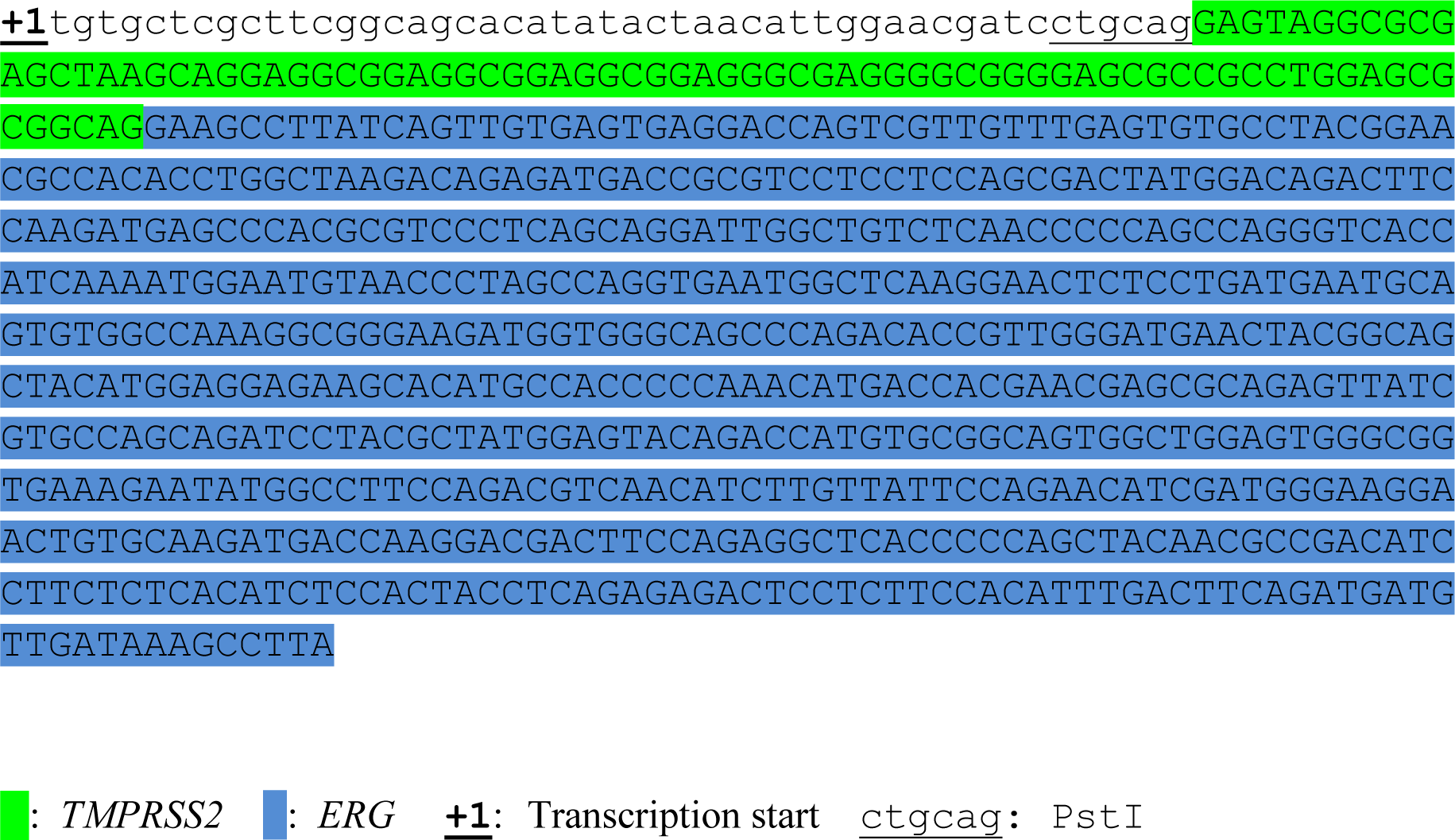

#### Antisense-1

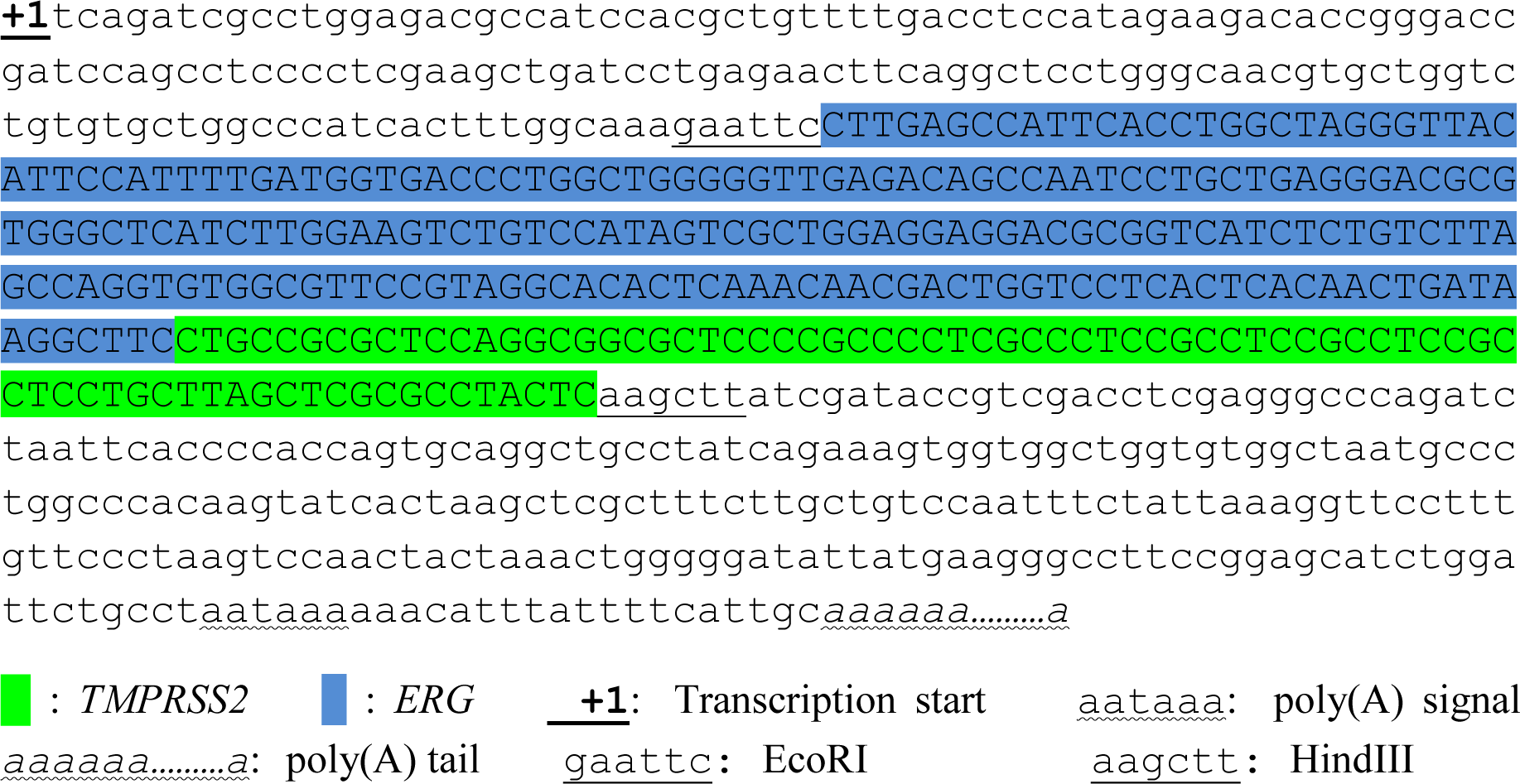

#### Antisense-2

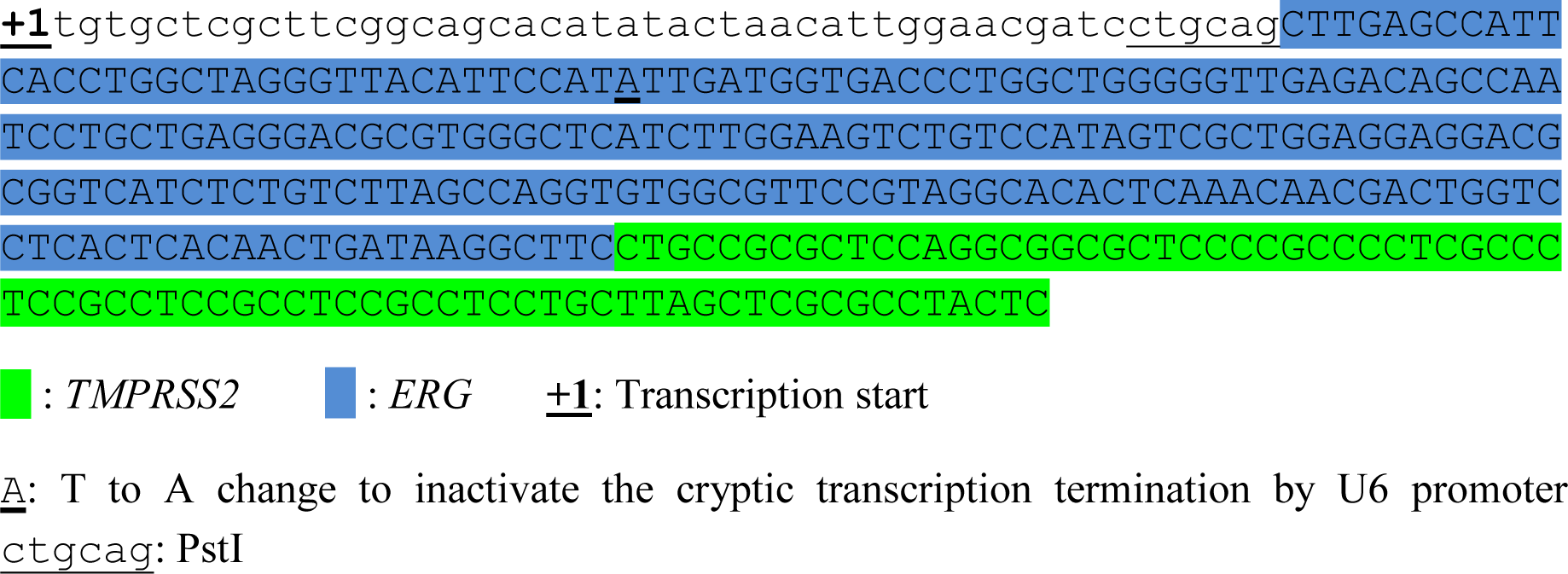

#### Antisense-3

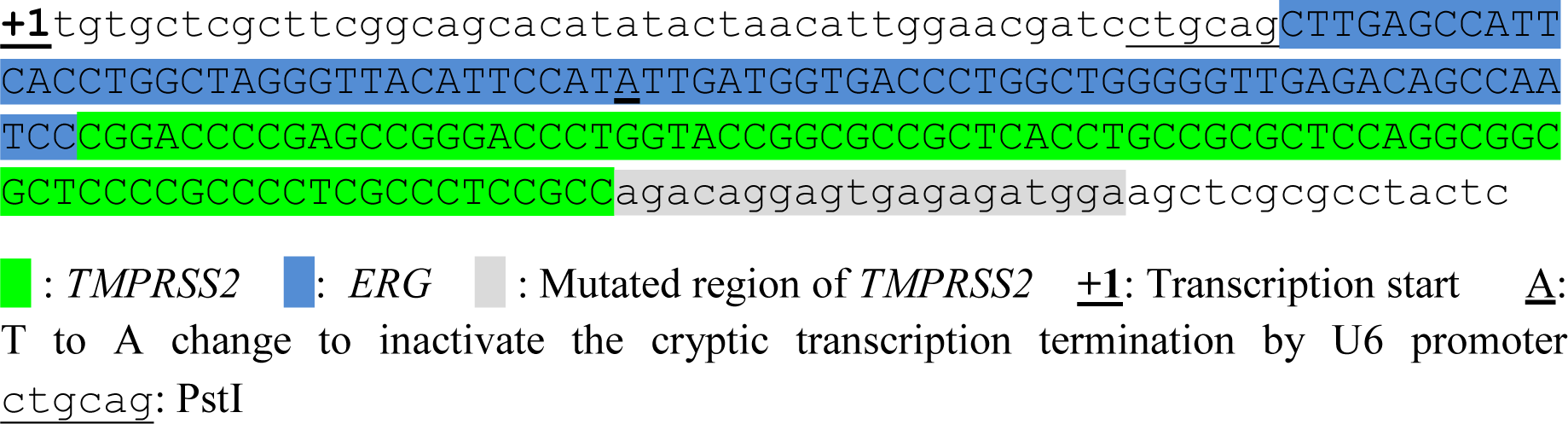

#### Antisense-4

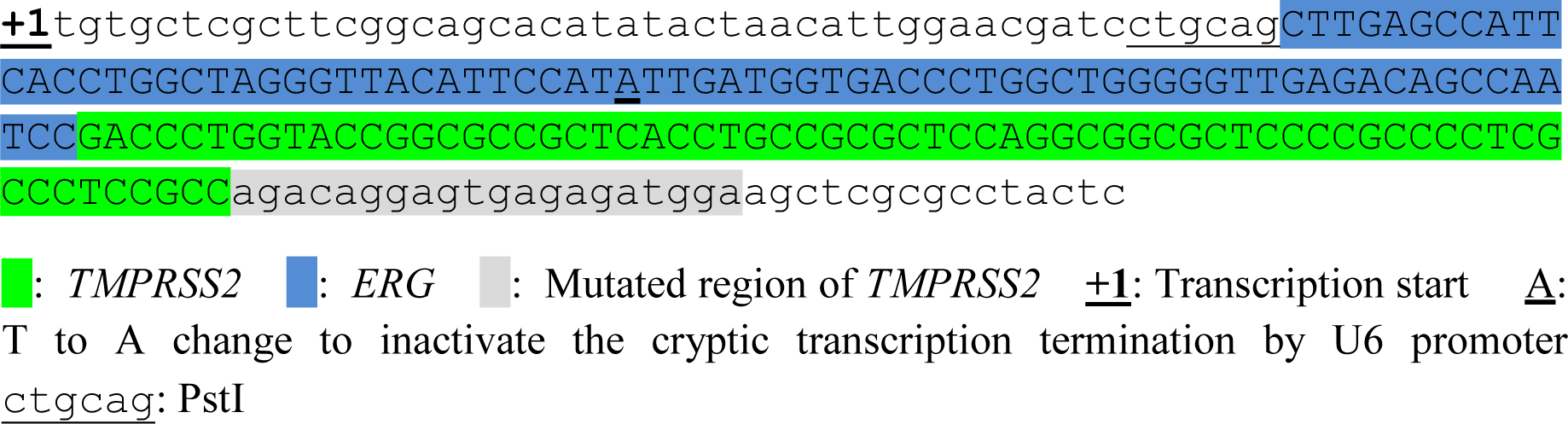

#### Antisense-5

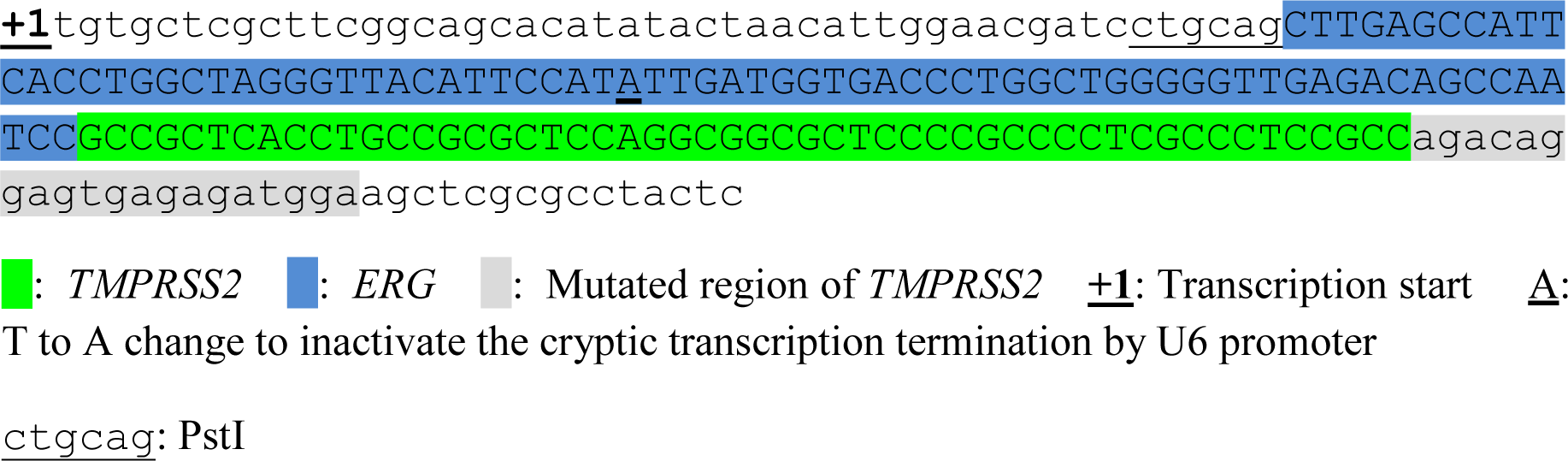

#### Antisense-6

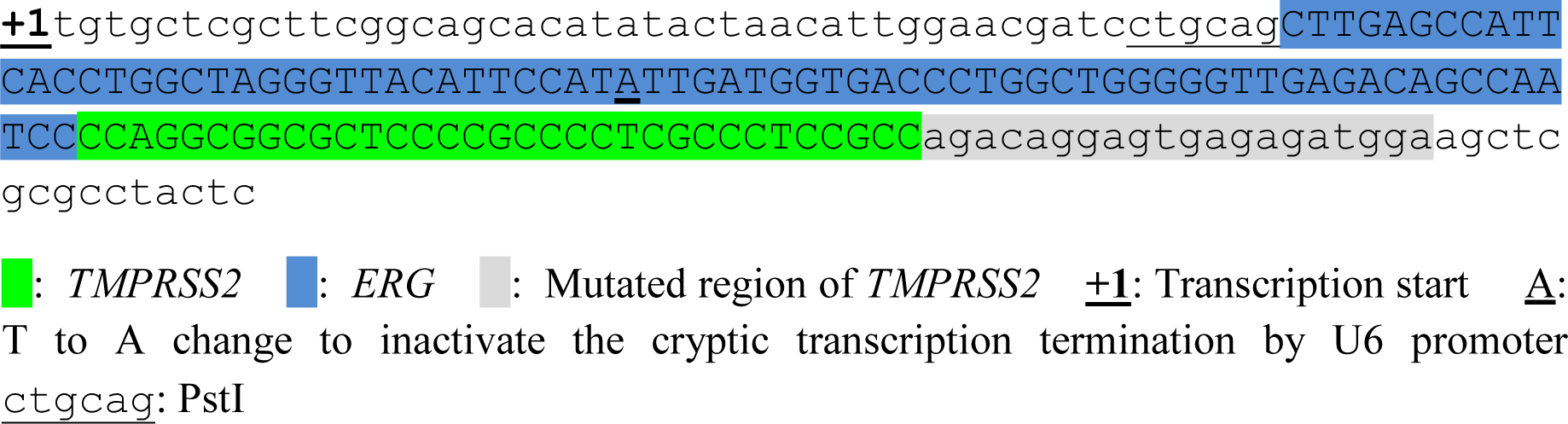

#### Antisense-7

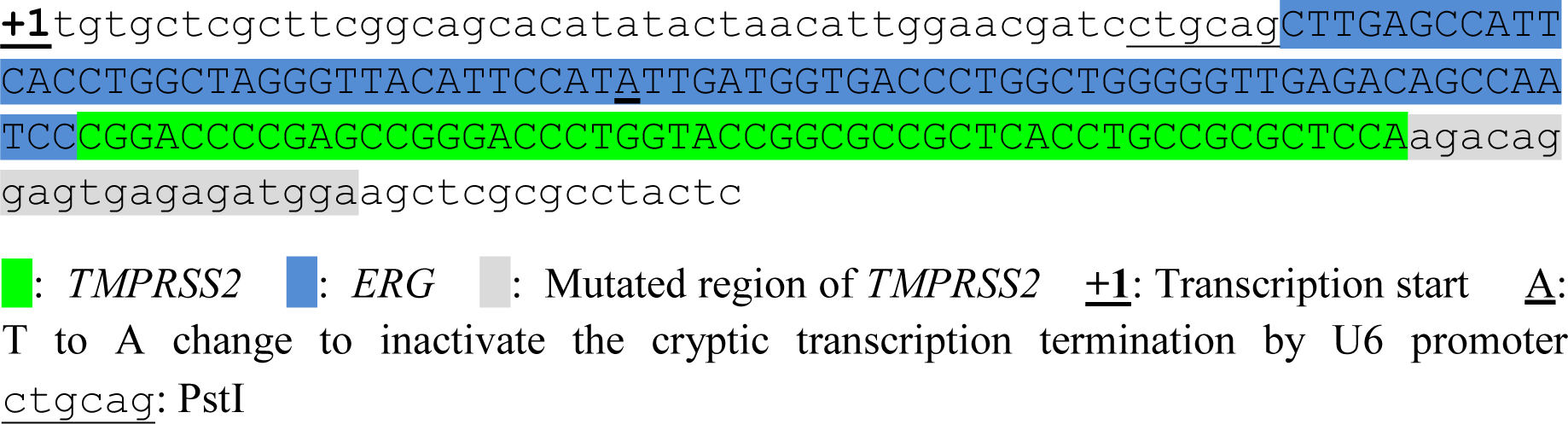

#### Antisense-8

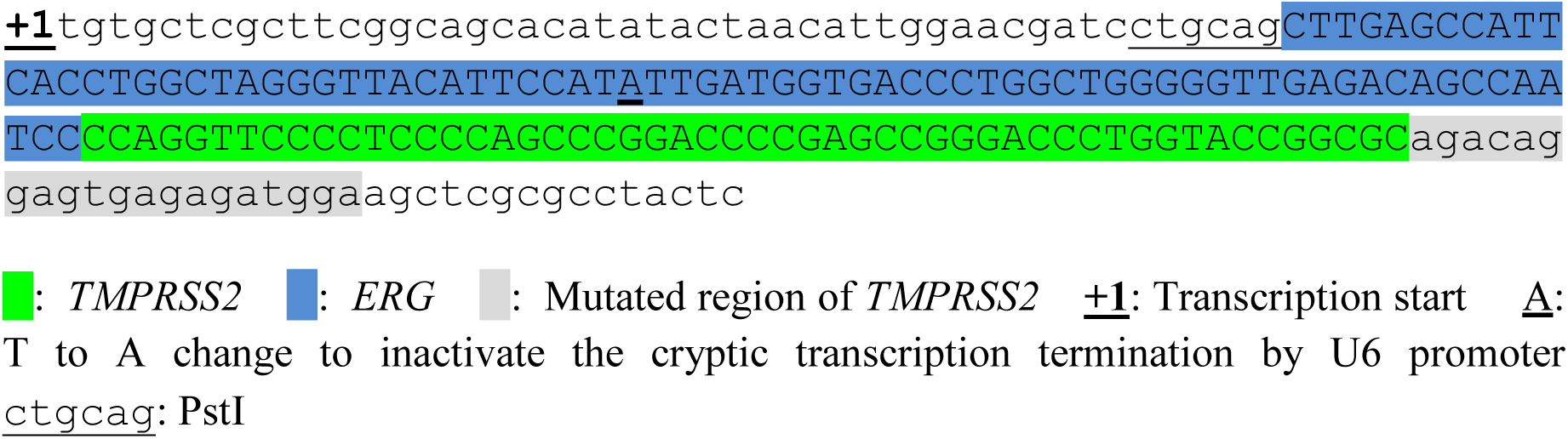

#### Antisense-9

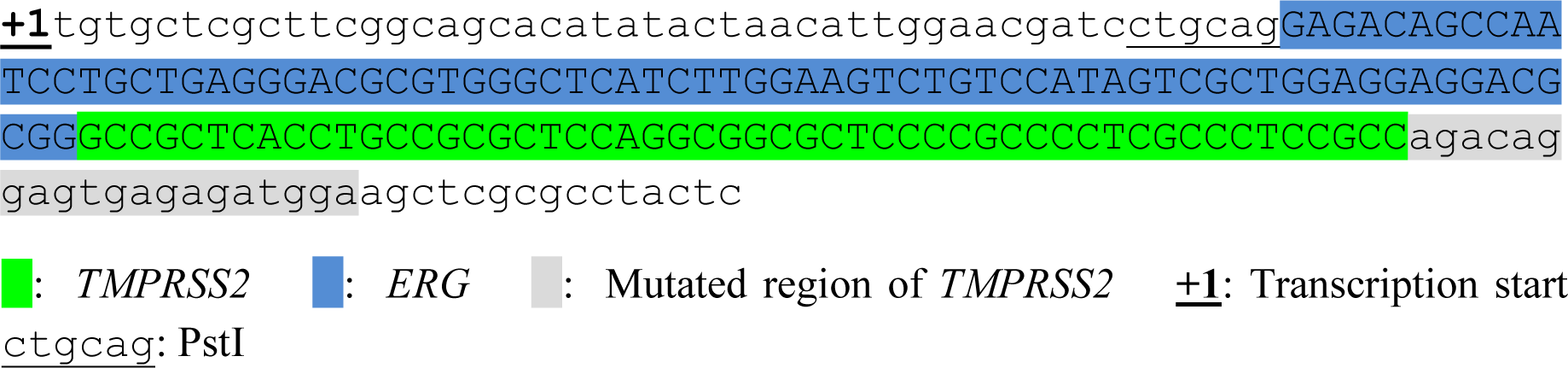

#### Antisense-5A

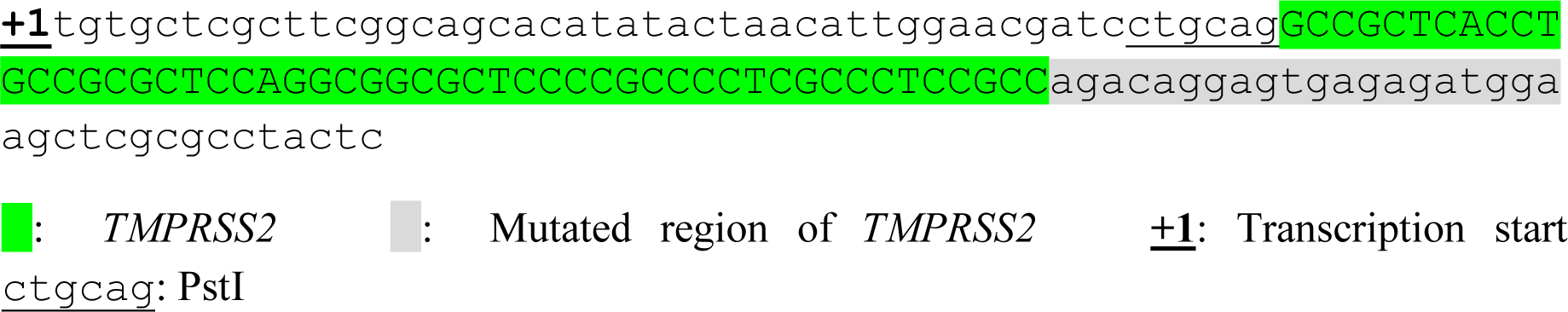

#### Antisense-5B

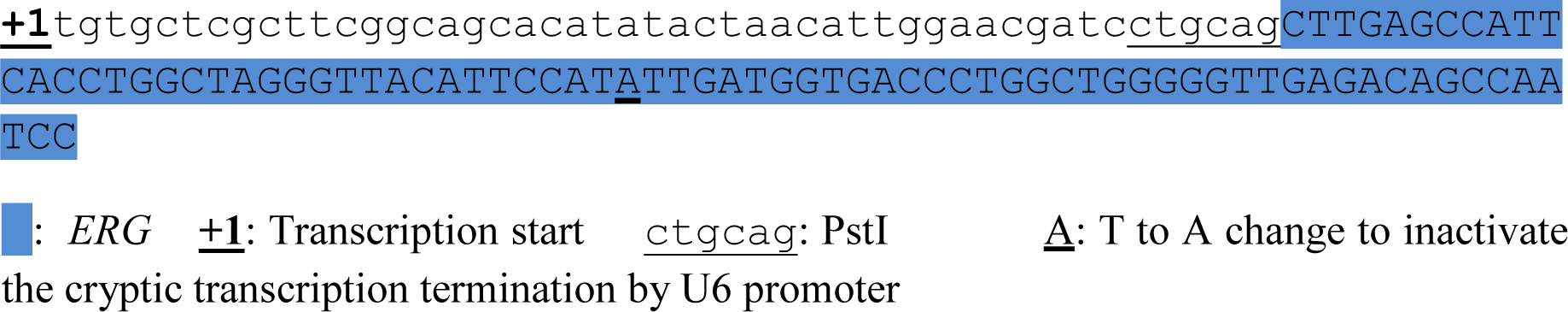

#### Antisense-B1

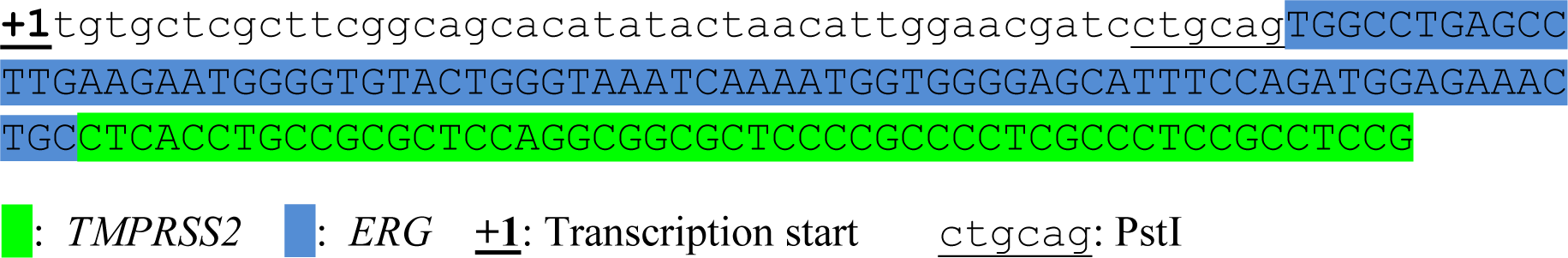

#### Antisense-B2

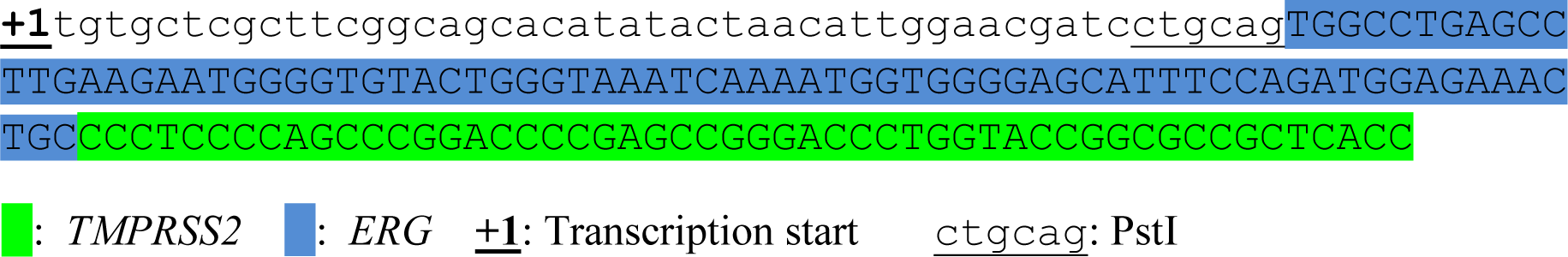

#### Antisense-B3

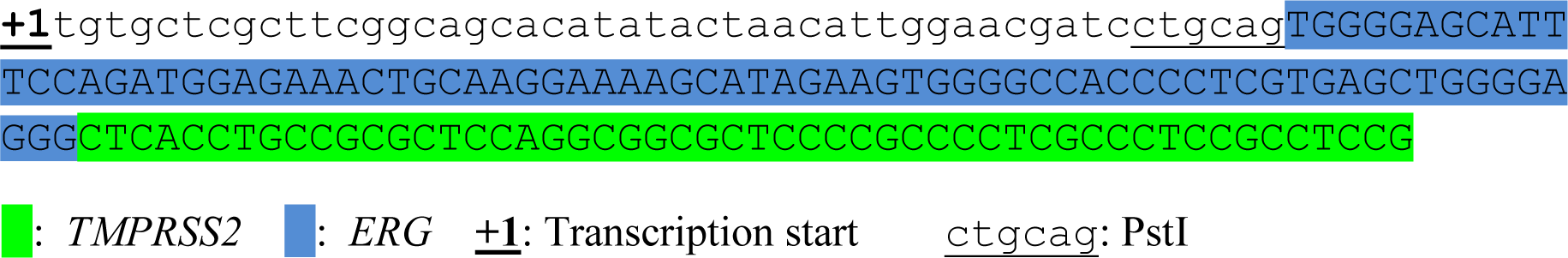

#### Antisense-C1

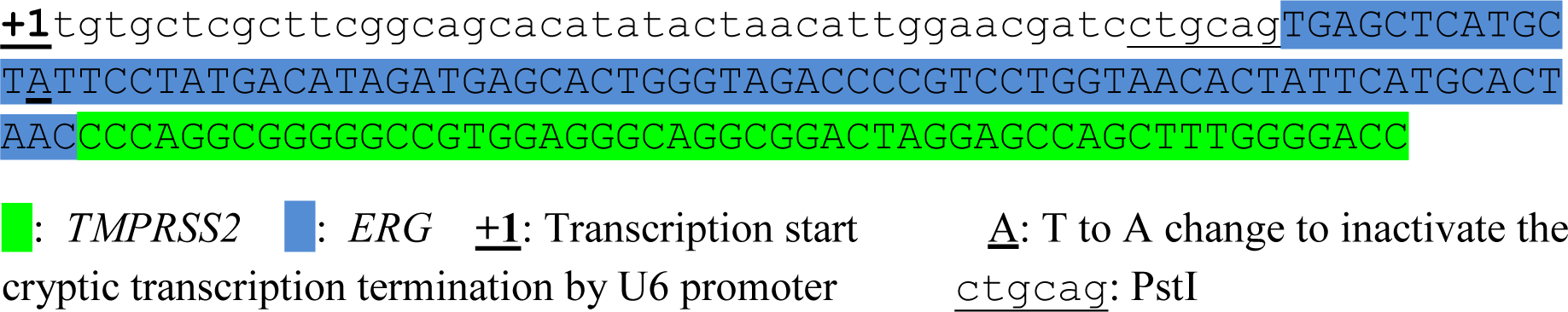

#### Antisense-C2

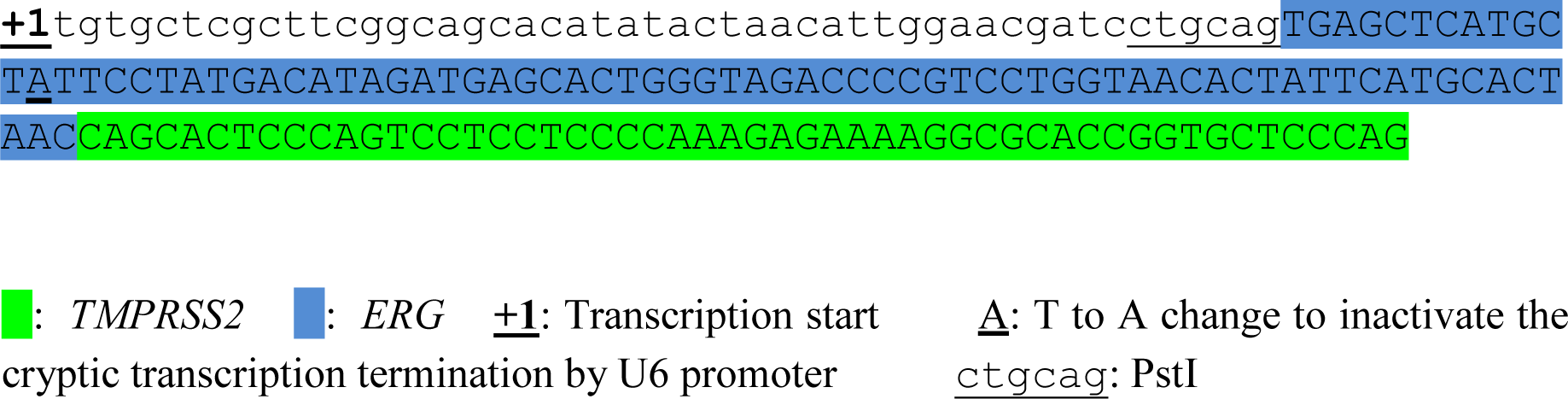

#### Antisense-C3

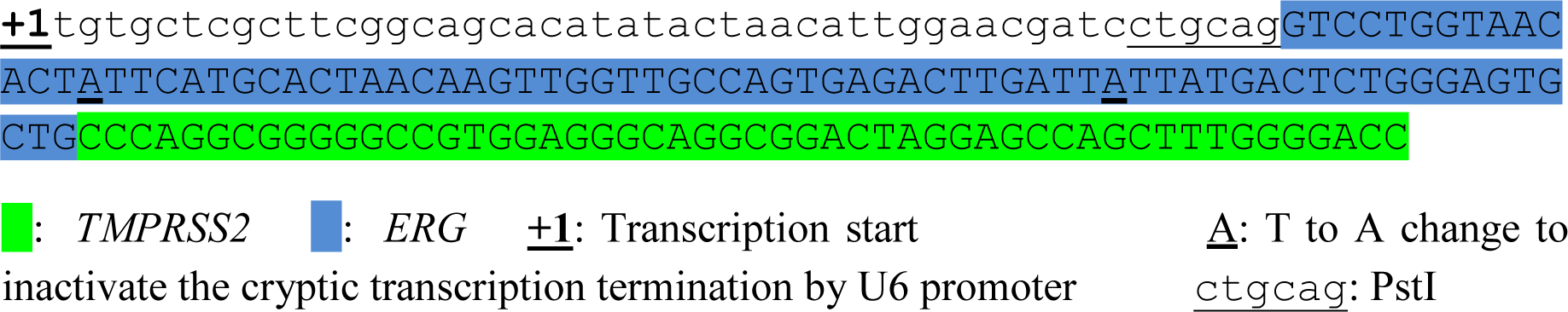

#### Antisense-D1

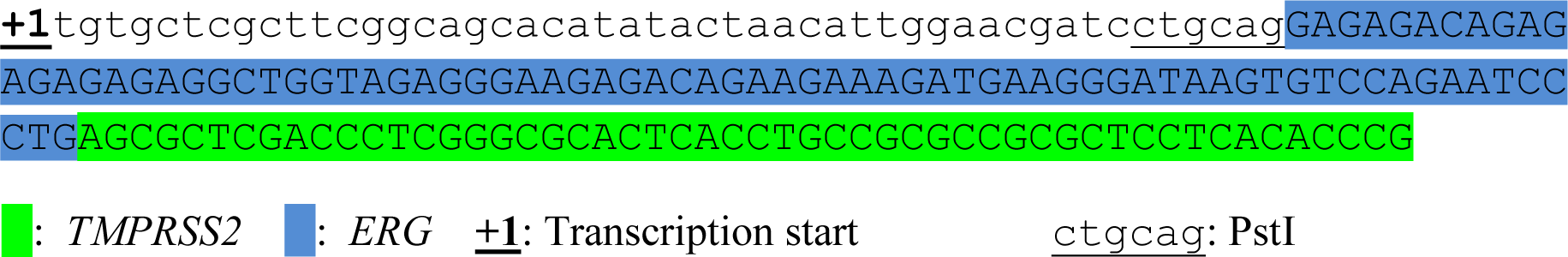

#### Antisense-D2

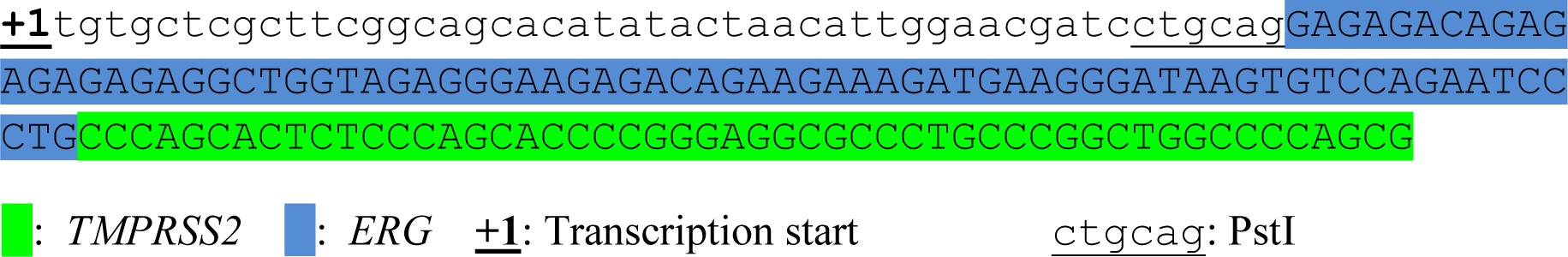

#### Antisense-D3

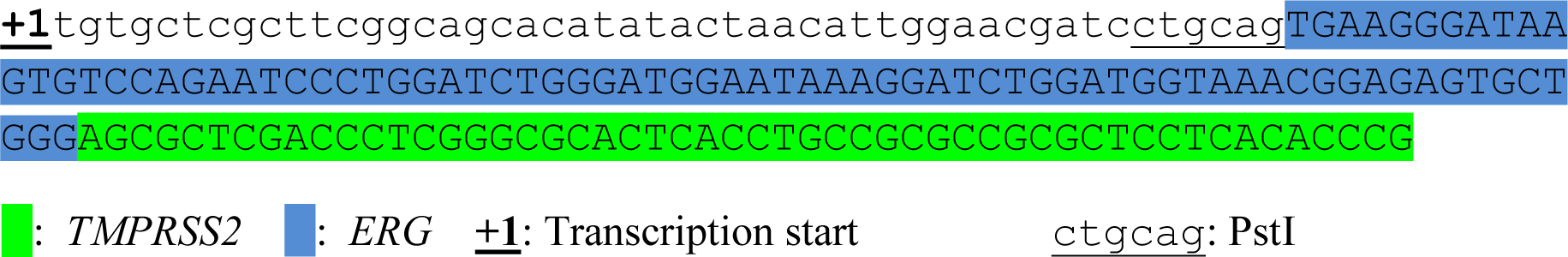

#### Antisense-E1

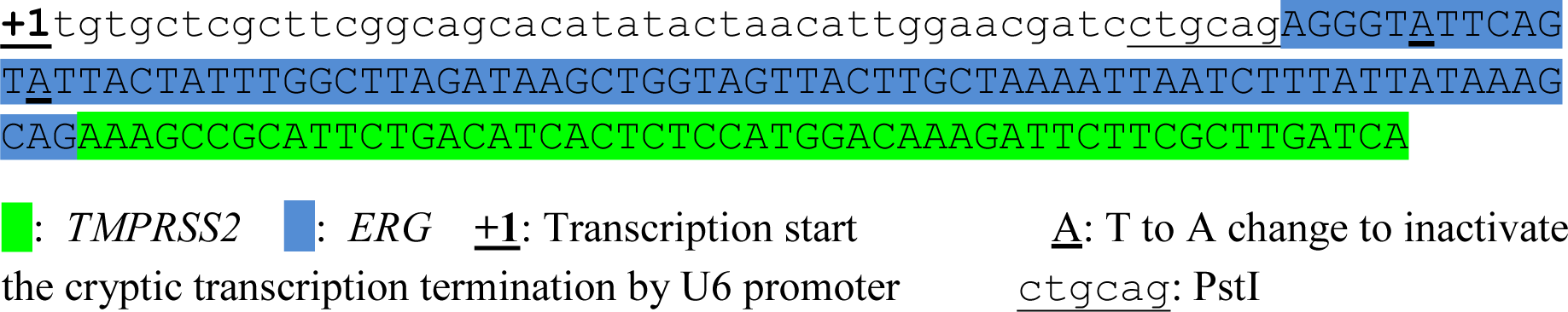

#### Antisense-F1

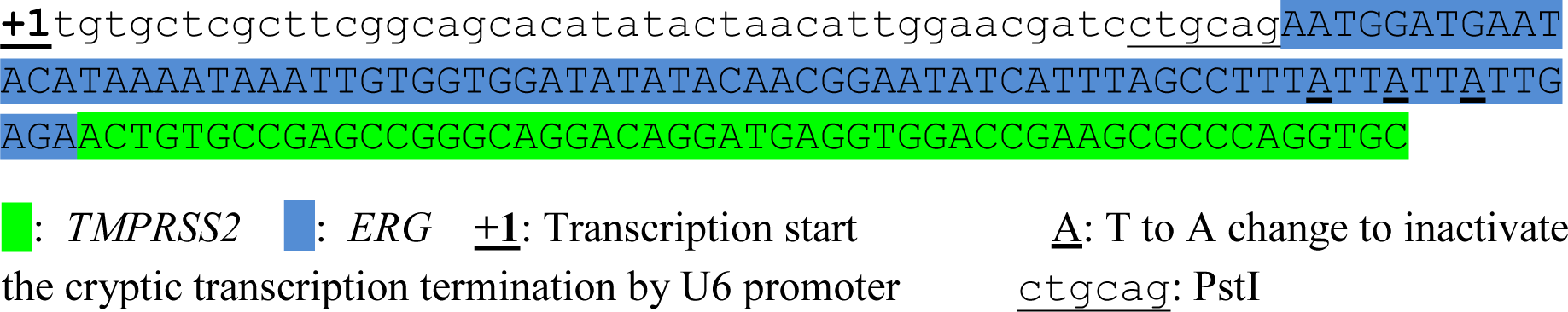

#### Antisense-G1

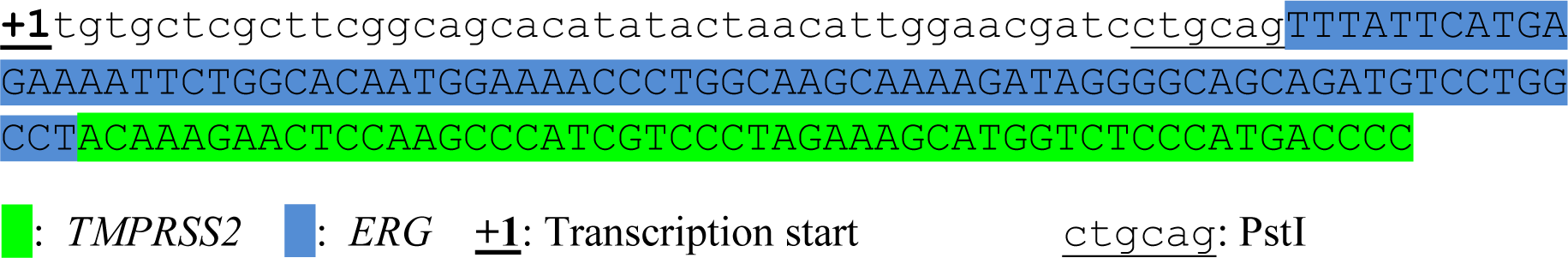

#### Sense-3

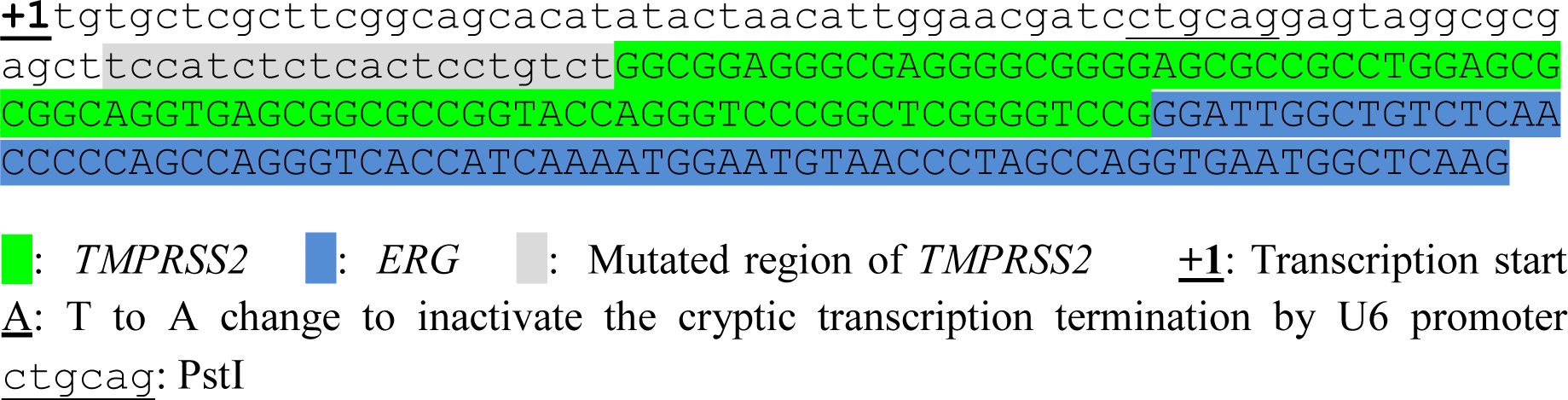

#### Sense-4

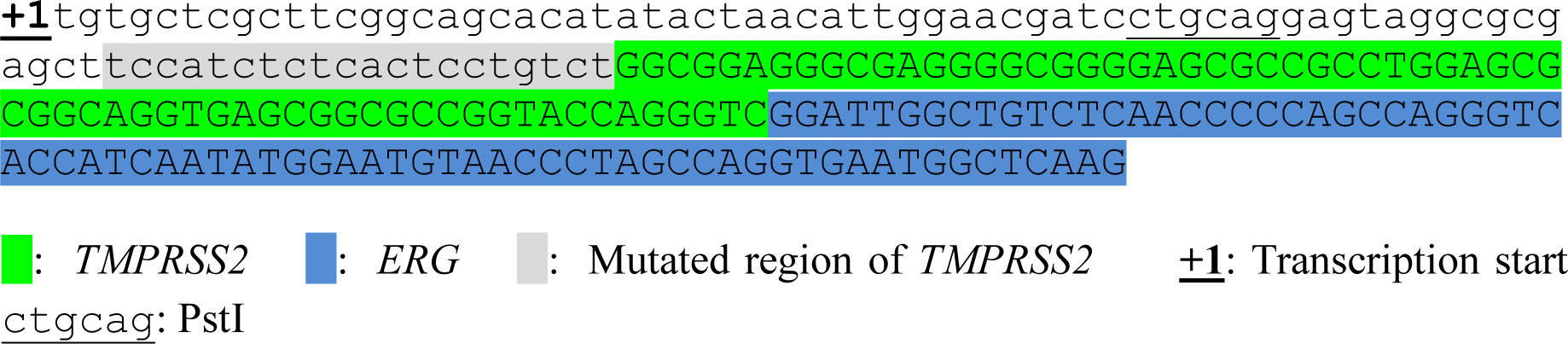

#### Sense-5

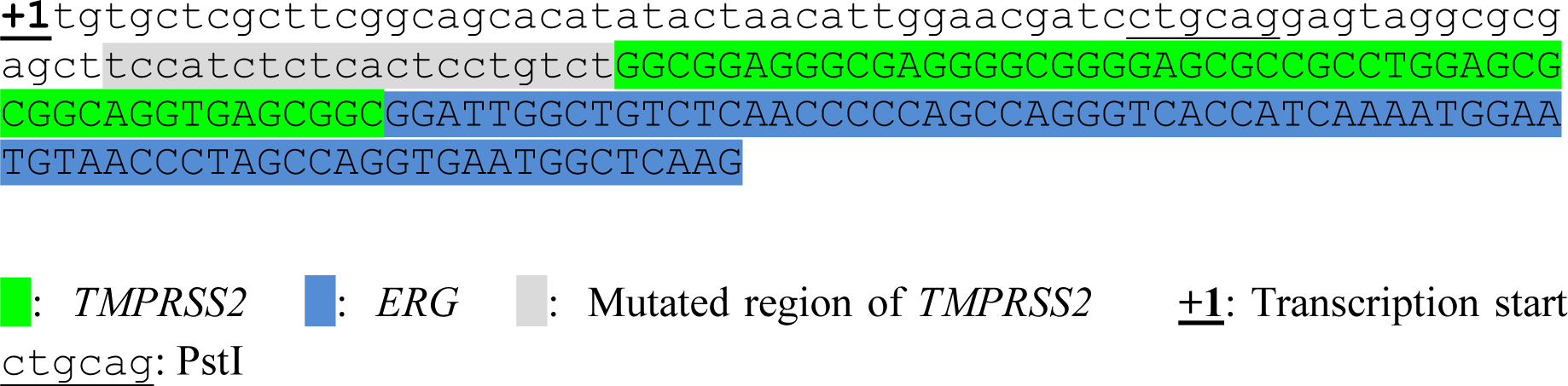

#### Sense-6

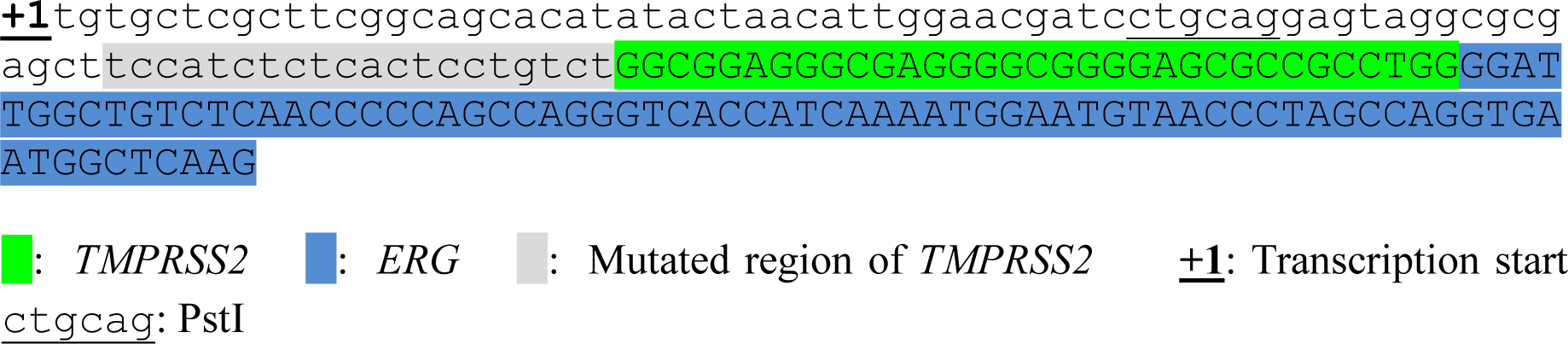

#### Sense-B1

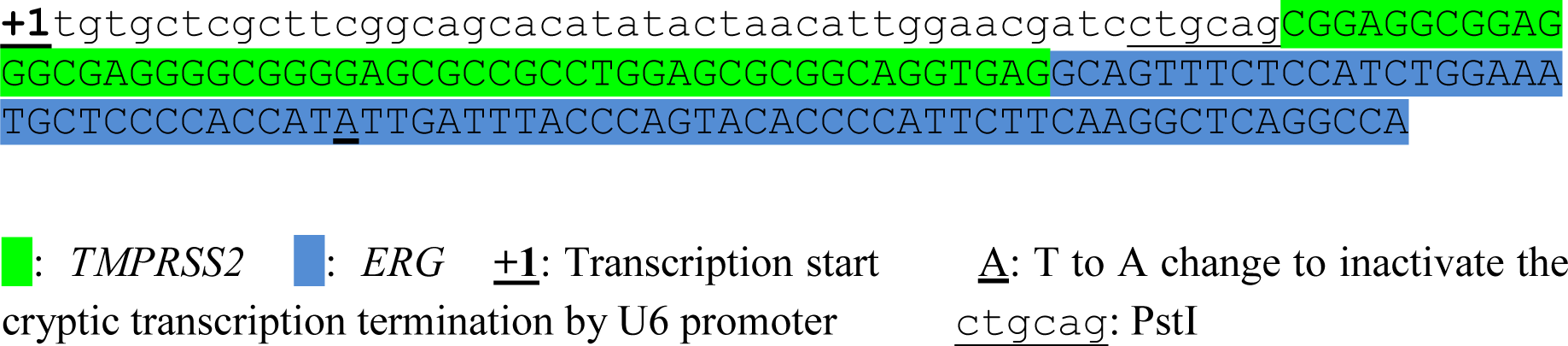

#### Sense-C1

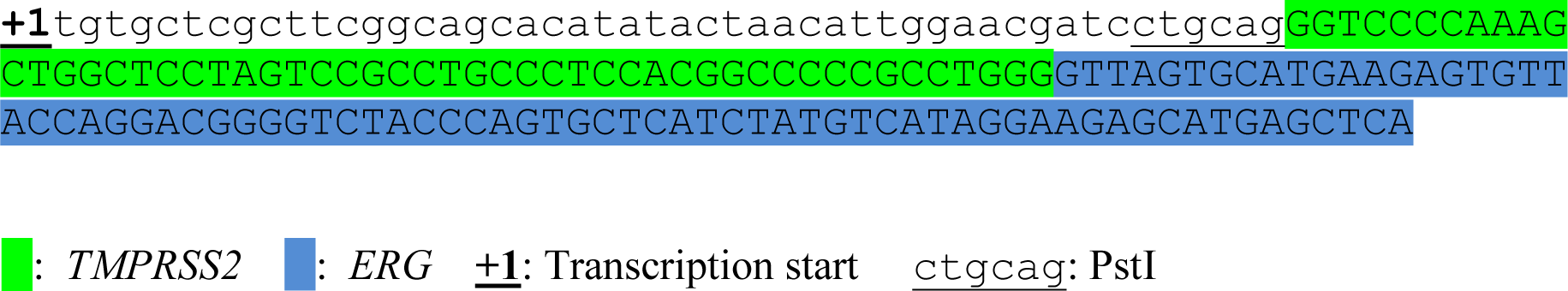

#### Sense-D1

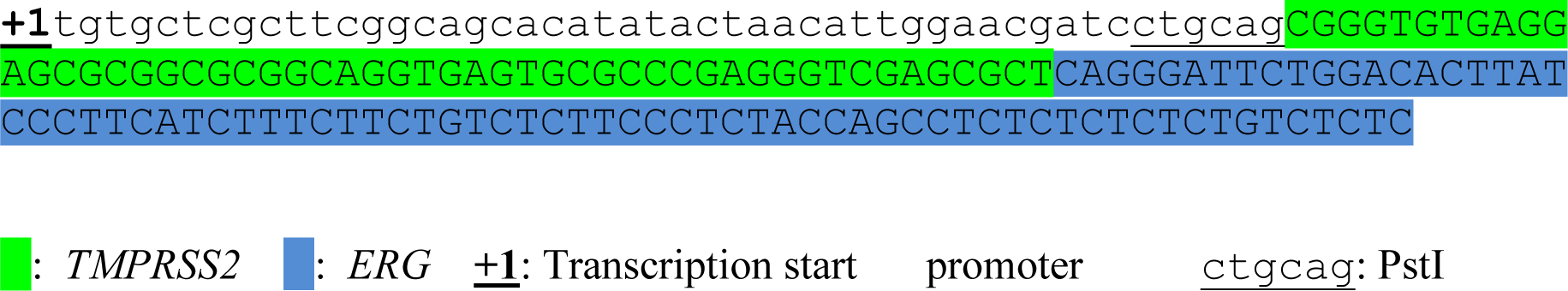

#### Antisense-*TMPRSS2-ETV1*-A1

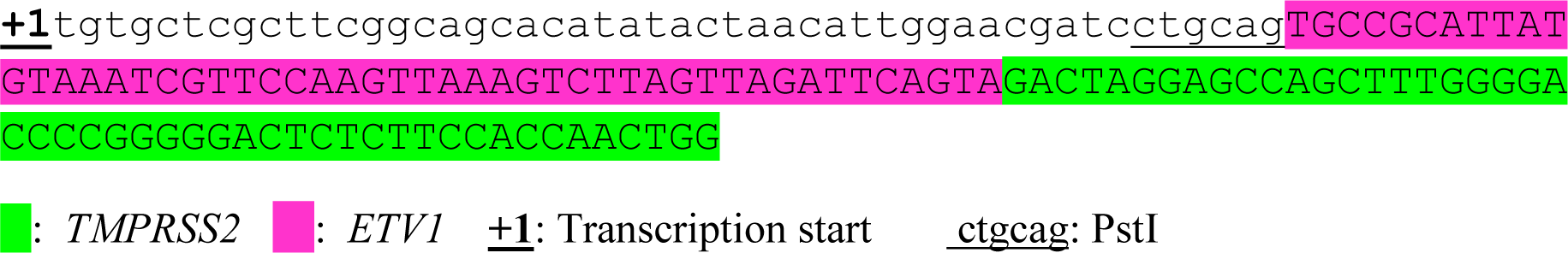

#### Antisense-TMPRSS2-ETV1-B1

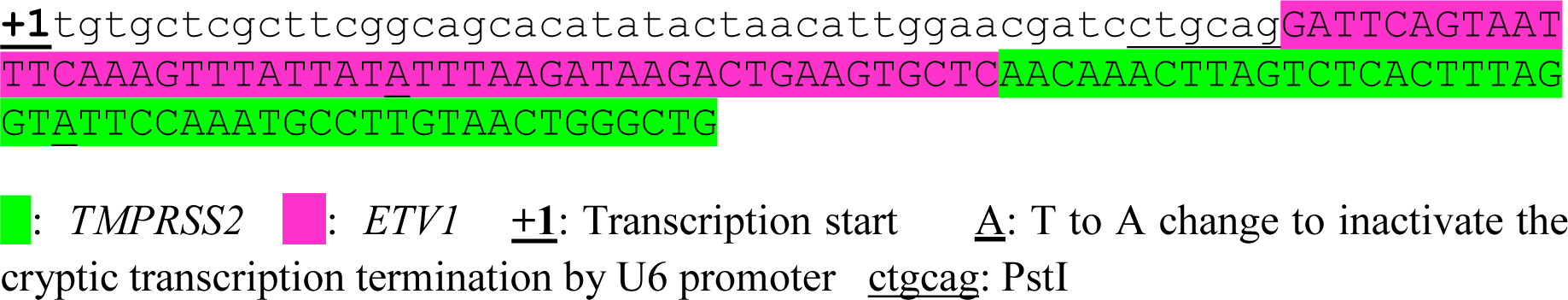

#### Antisense-TMPRSS2-ETV1-C1

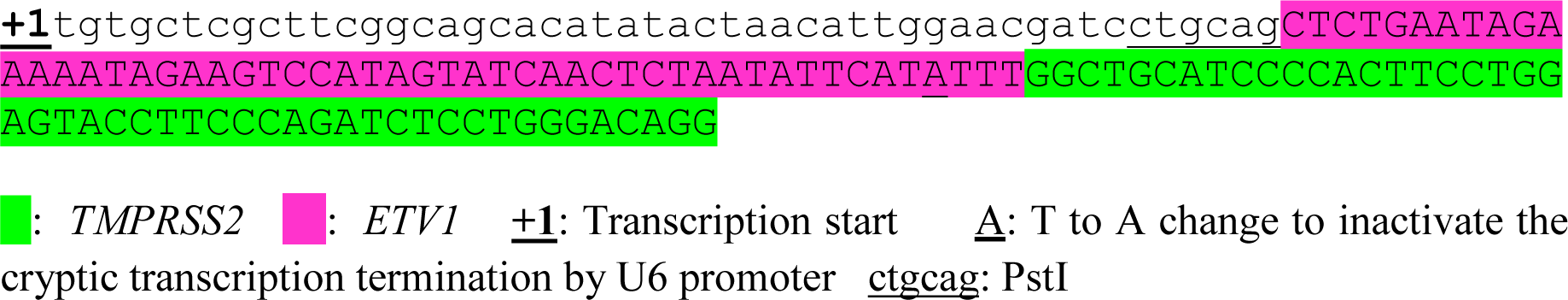

#### Antisense-TMPRSS2-ETV1-D1

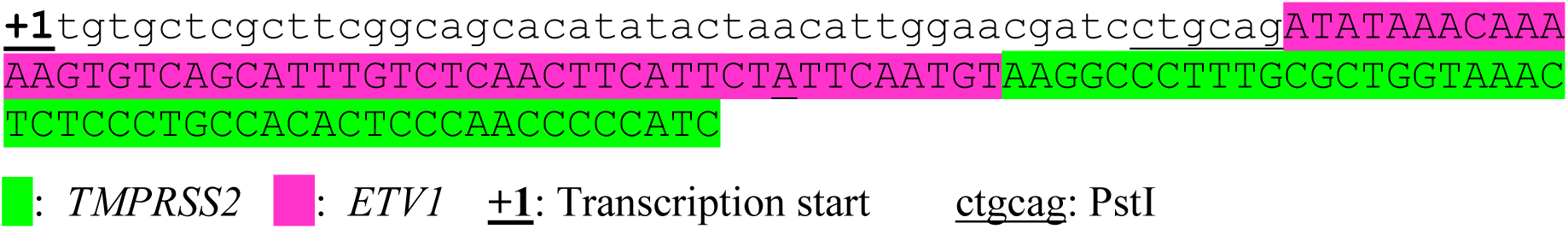

#### Antisense-TMPRSS2-ETV1-E1

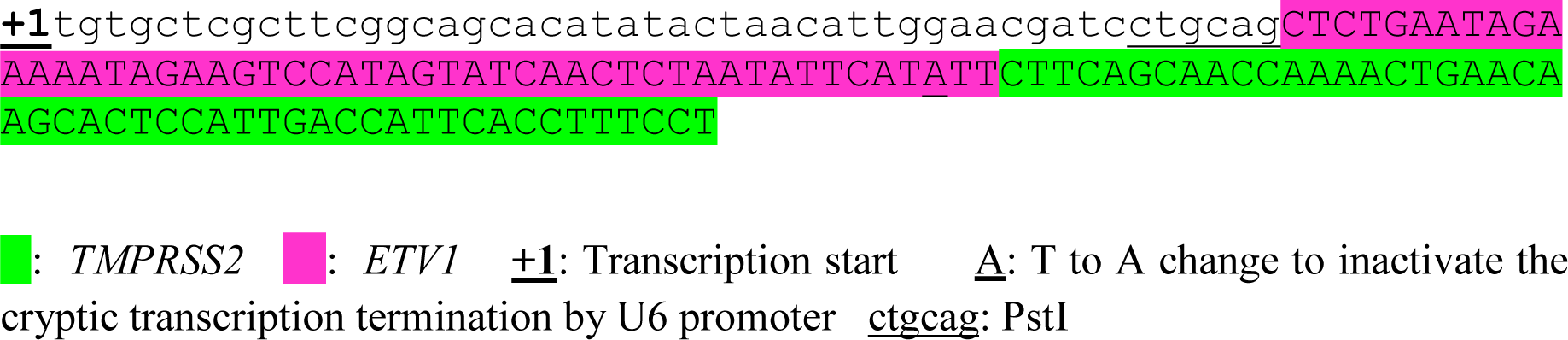

#### Antisense-TMPRSS2-ETV1-F1

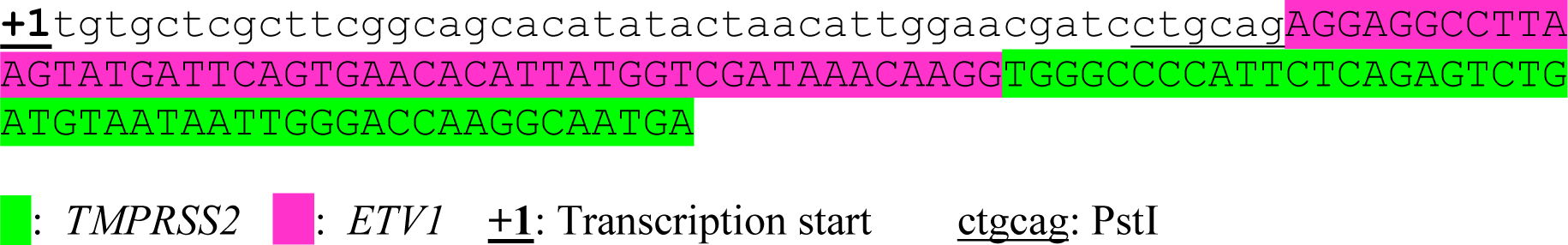

#### Antisense-TMPRSS2-ETV1-G1

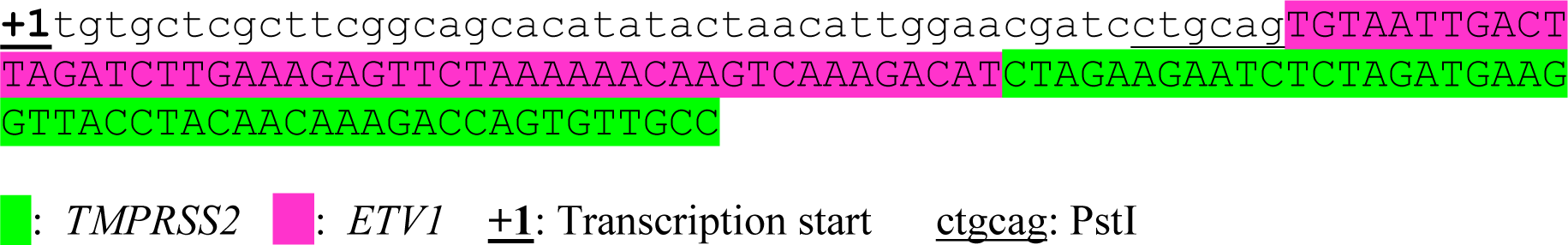

#### Antisense-TMPRSS2-ETV1-H1

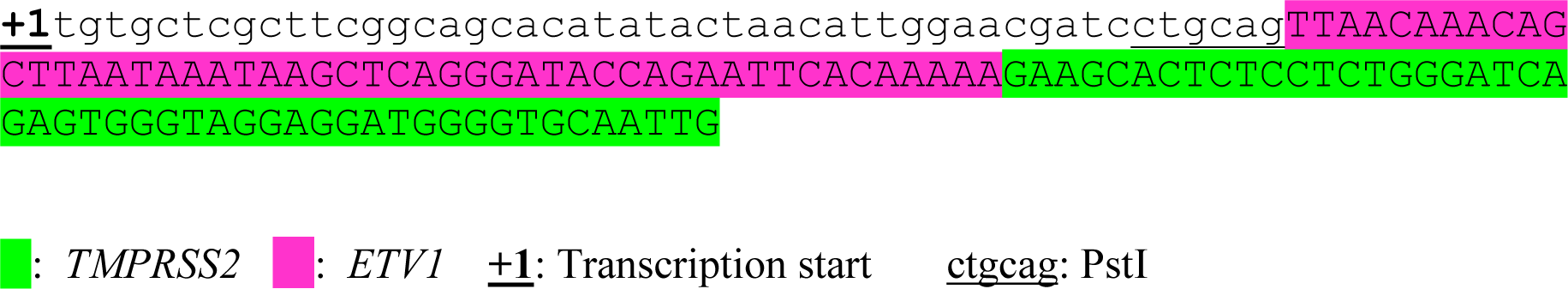

#### Sense-TMPRSS2-ETV1-A1

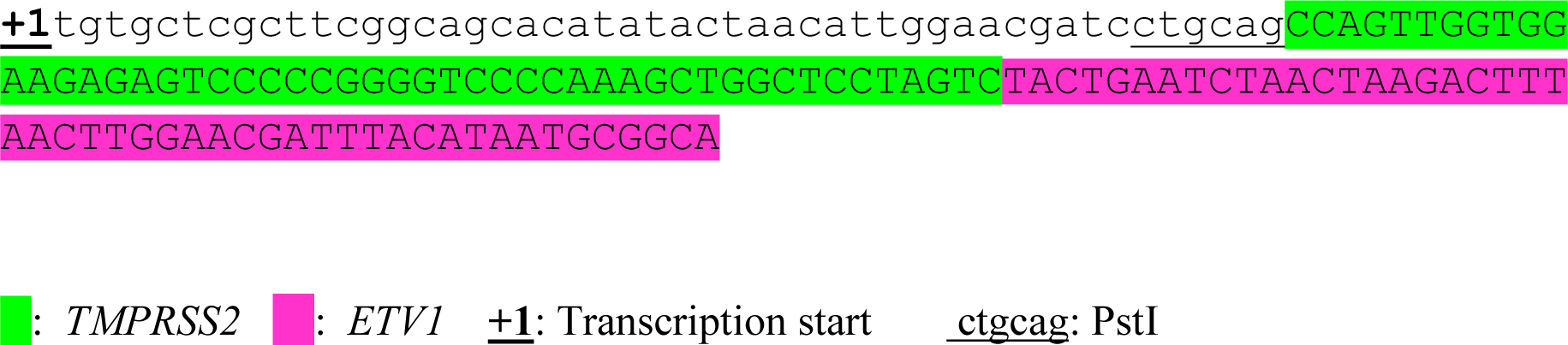

**<u>List of primers used</u>**

**RT-PCR primers for amplifying induced fusion RNAs:**

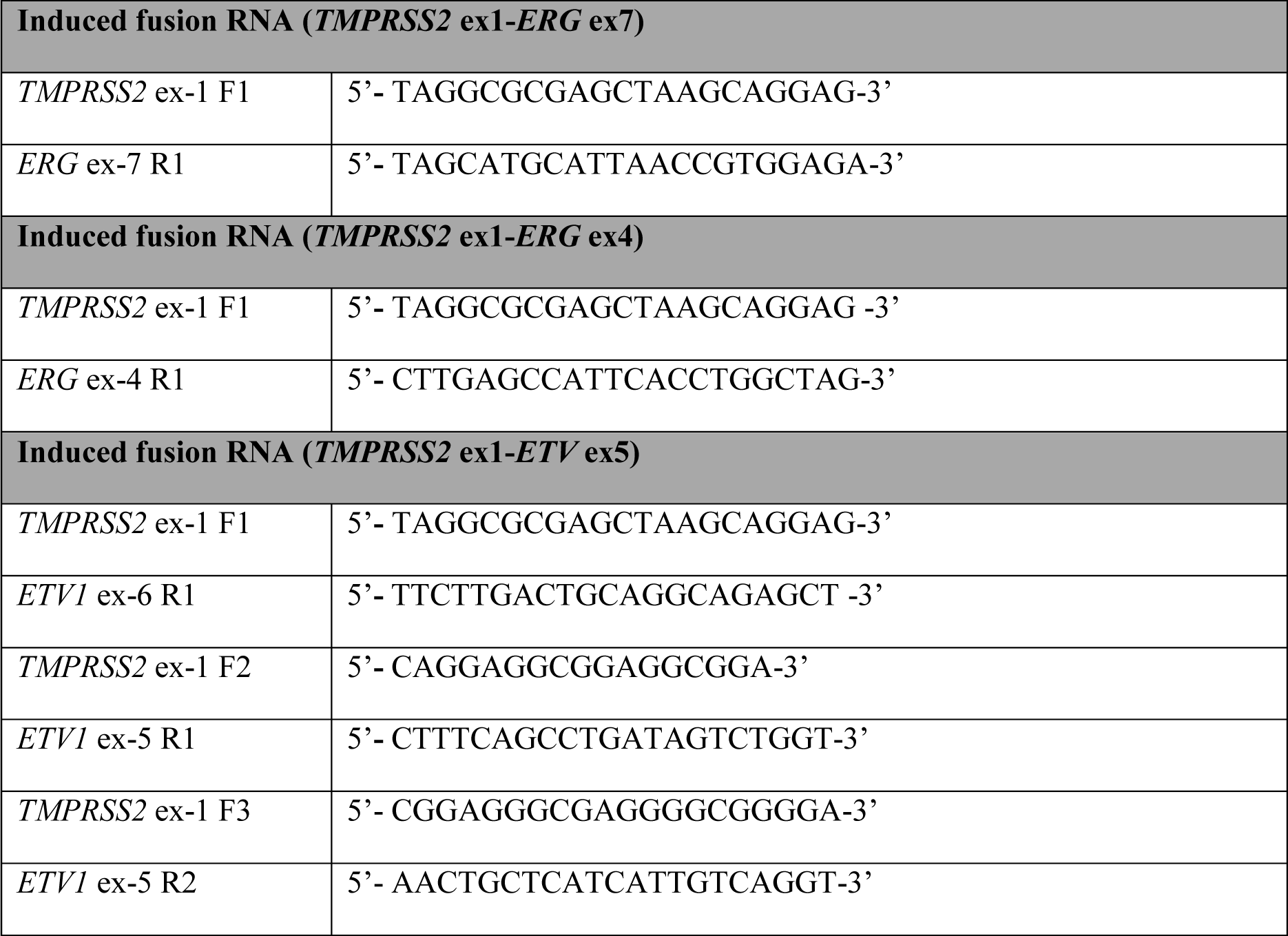

**Primers used in three-round PCR for amplifying induced fusion RNAs:**

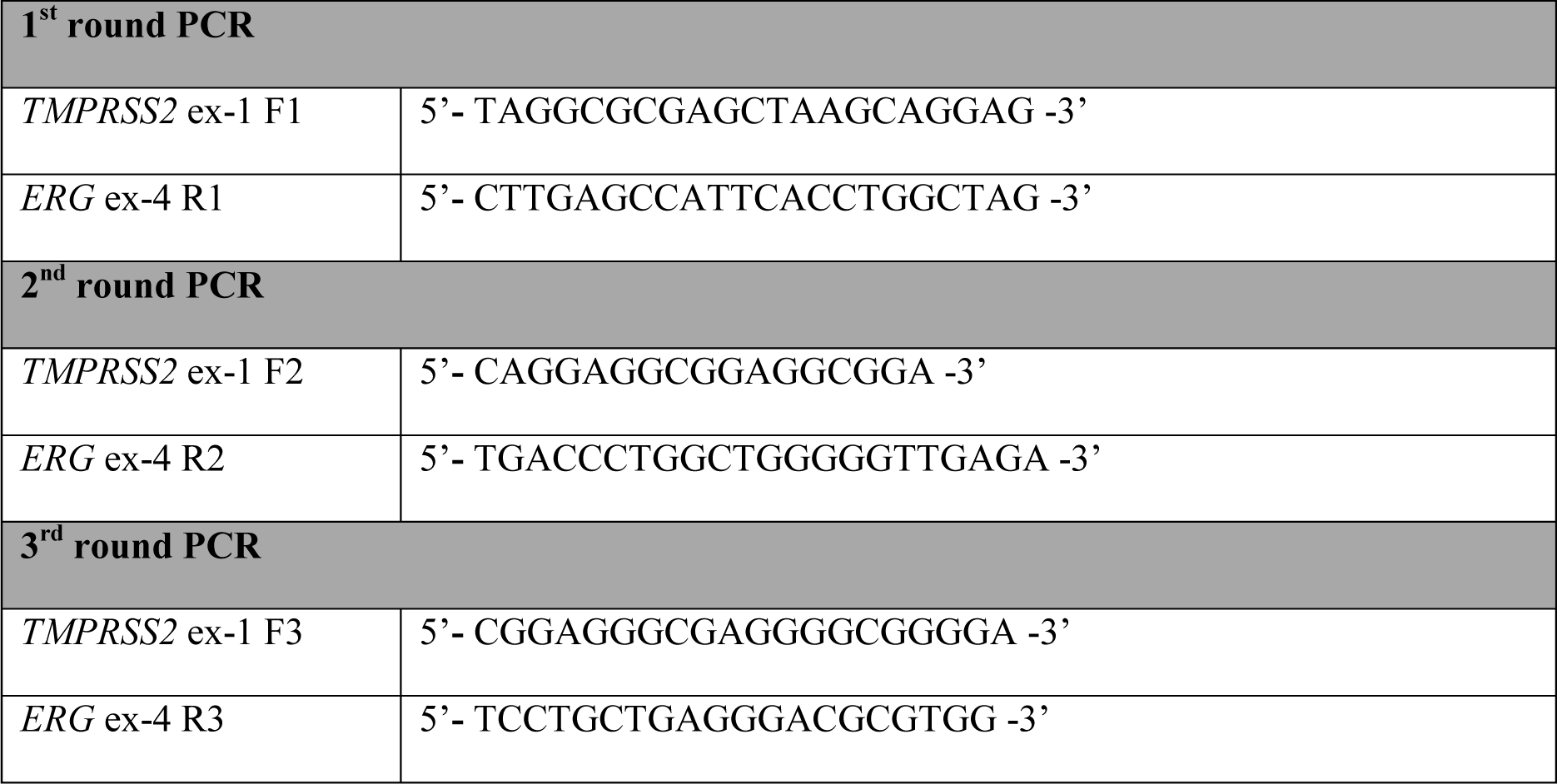

**RT-PCR primers for amplifying endogenous parental mRNAs:**

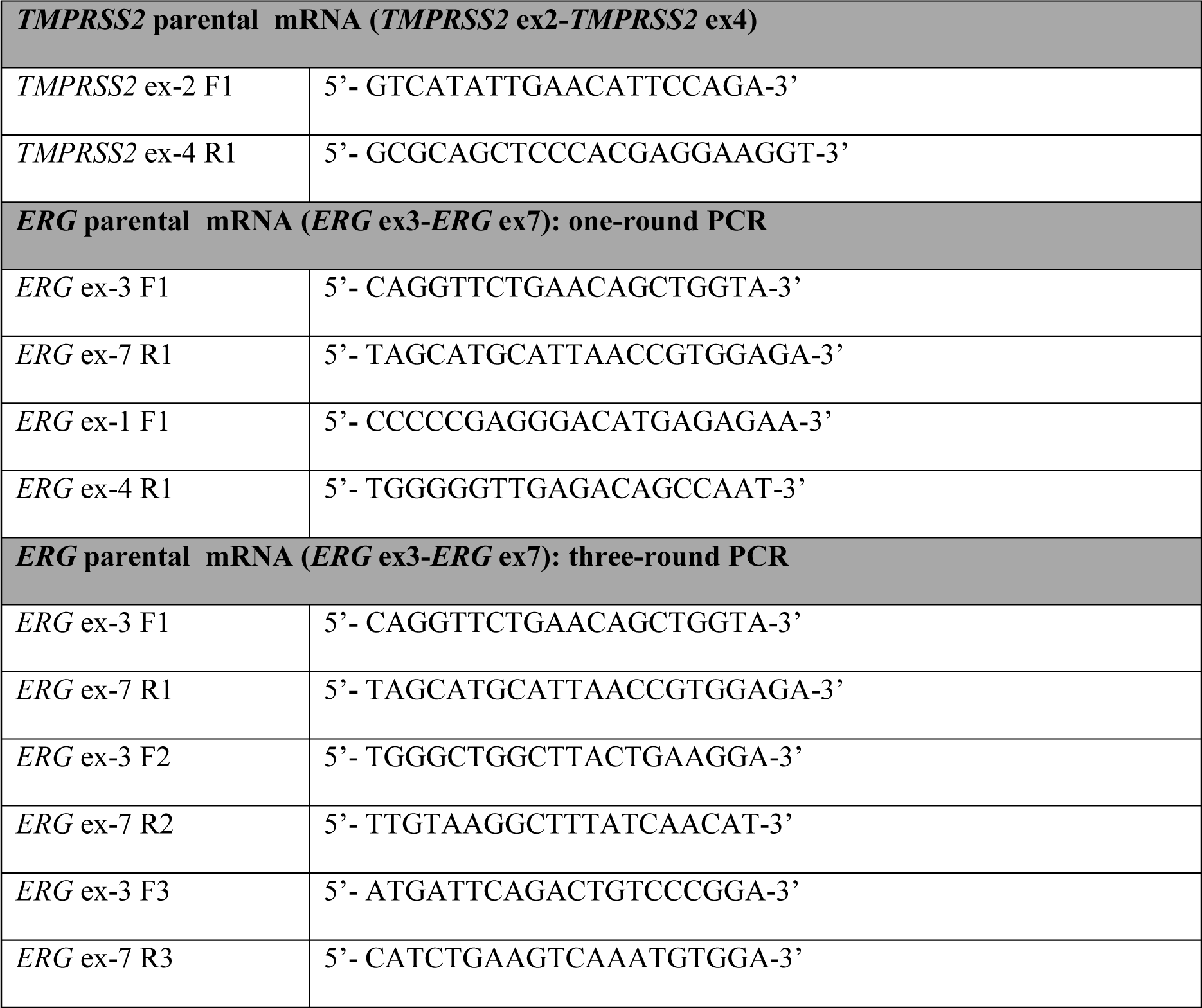

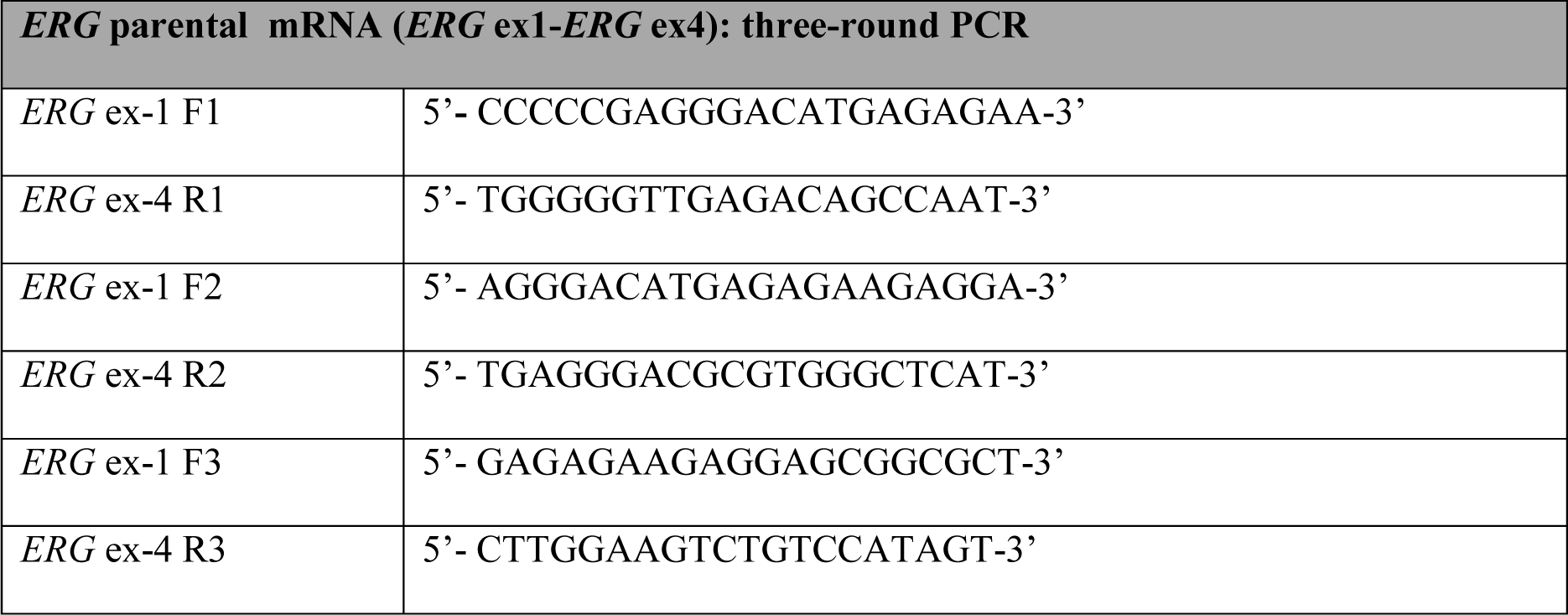

**RT-PCR primers for amplifying BCAM-AKT2 chimeric RNA:**

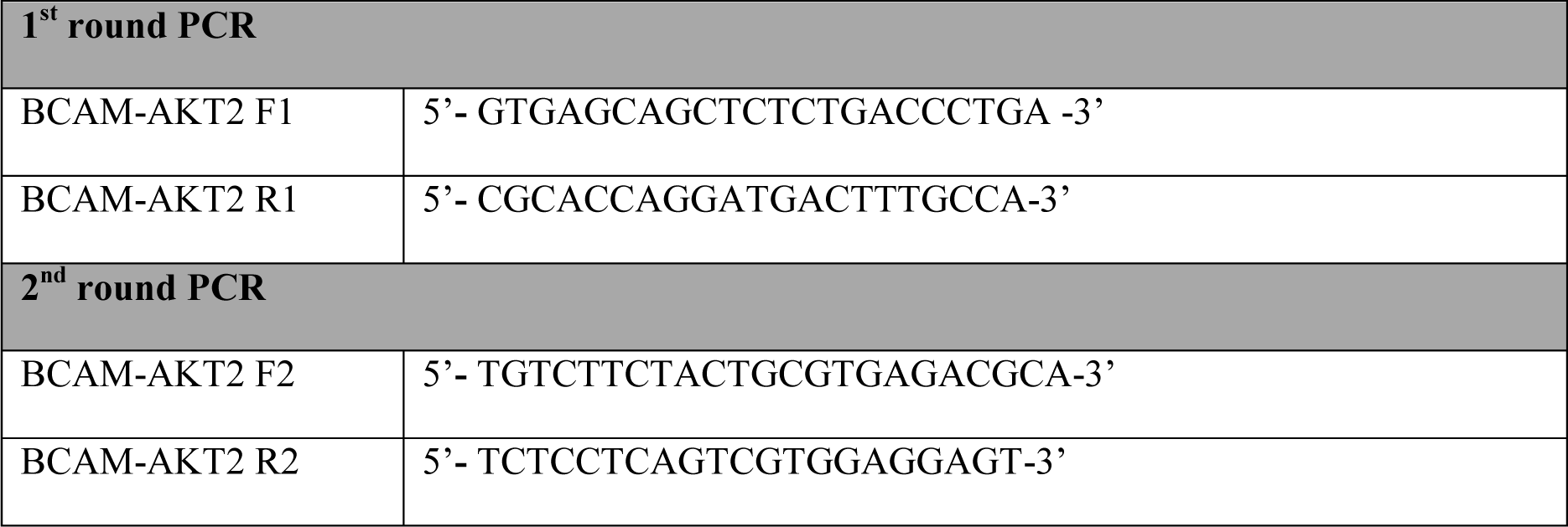

**RT-PCR primers for amplifying input RNAs:**

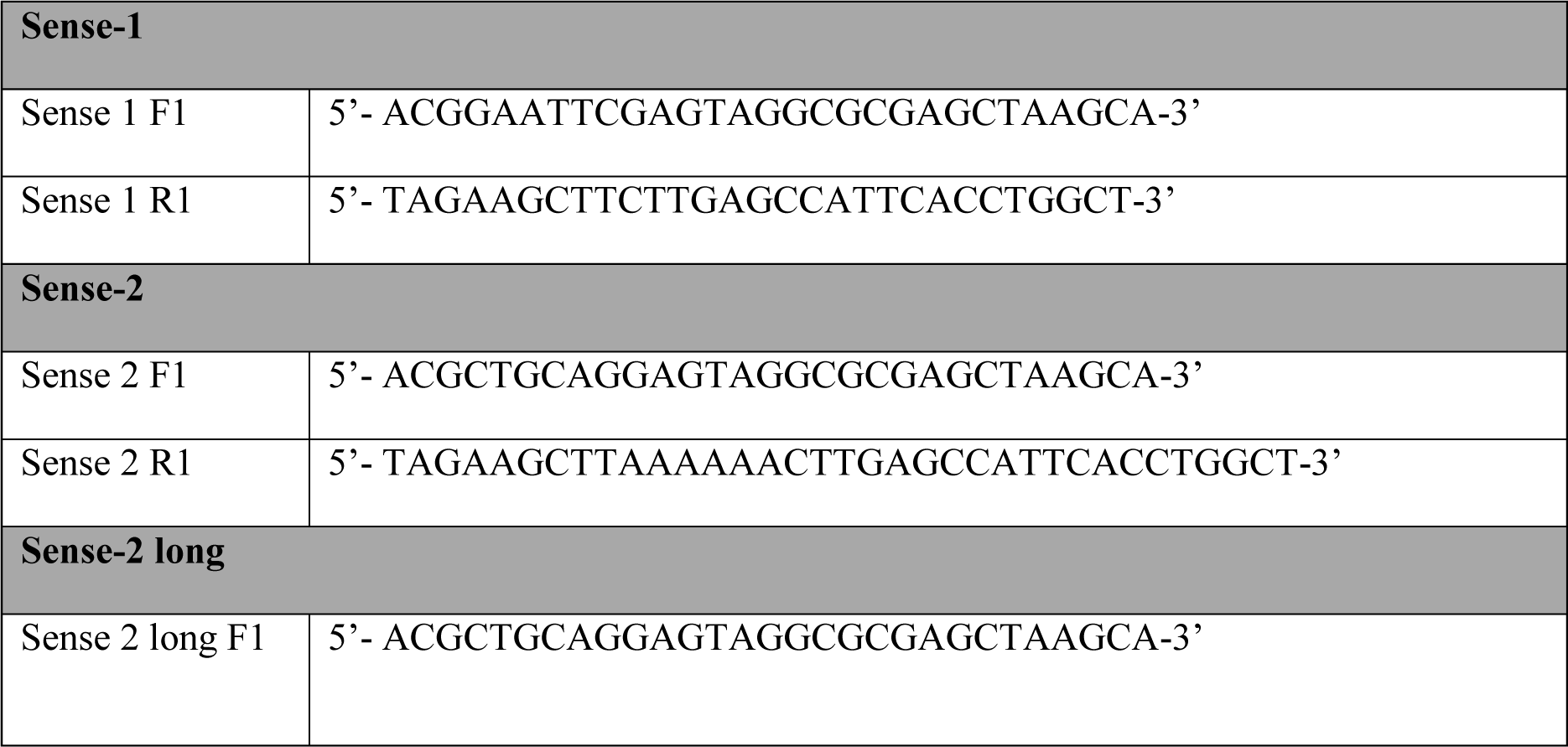

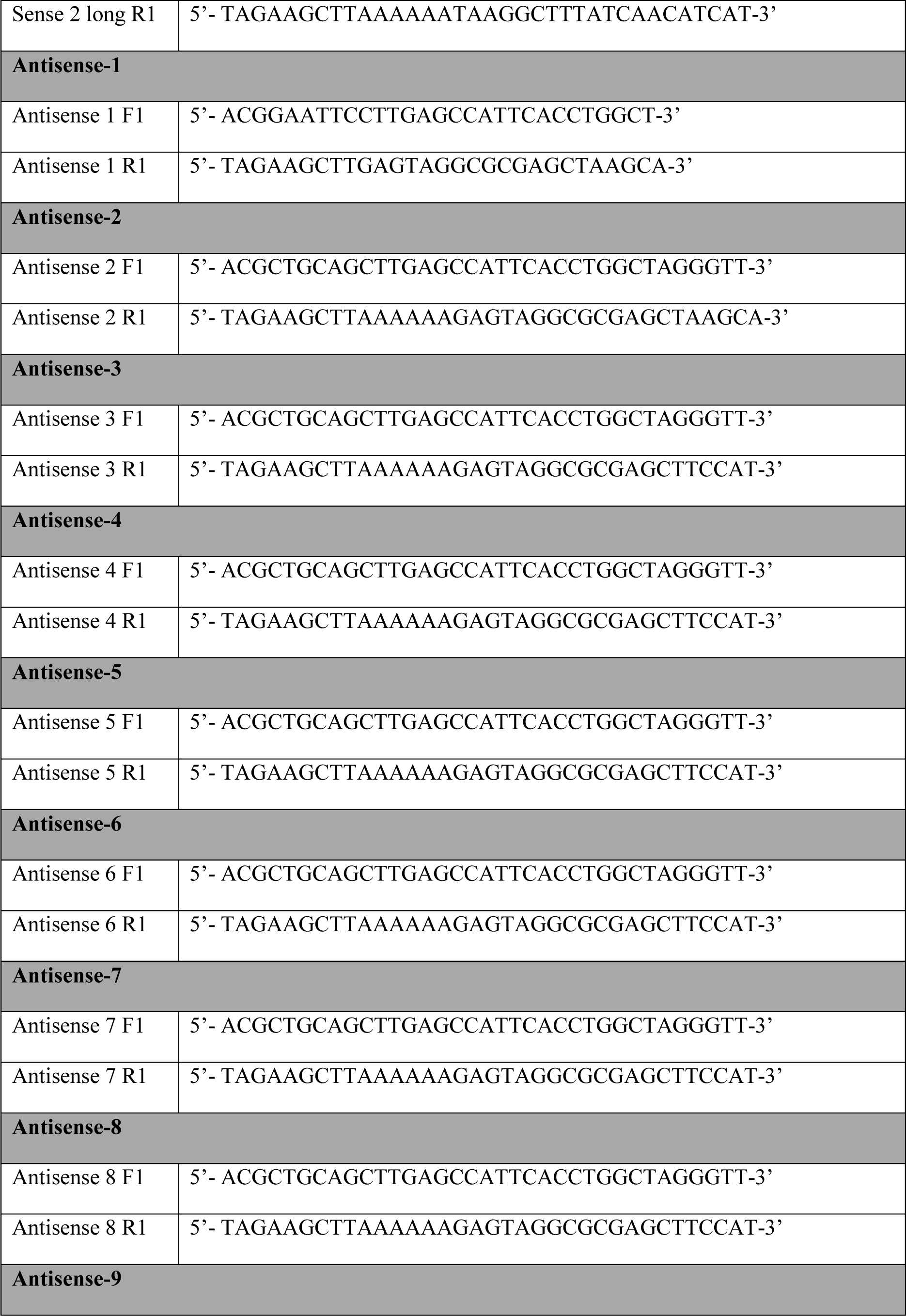

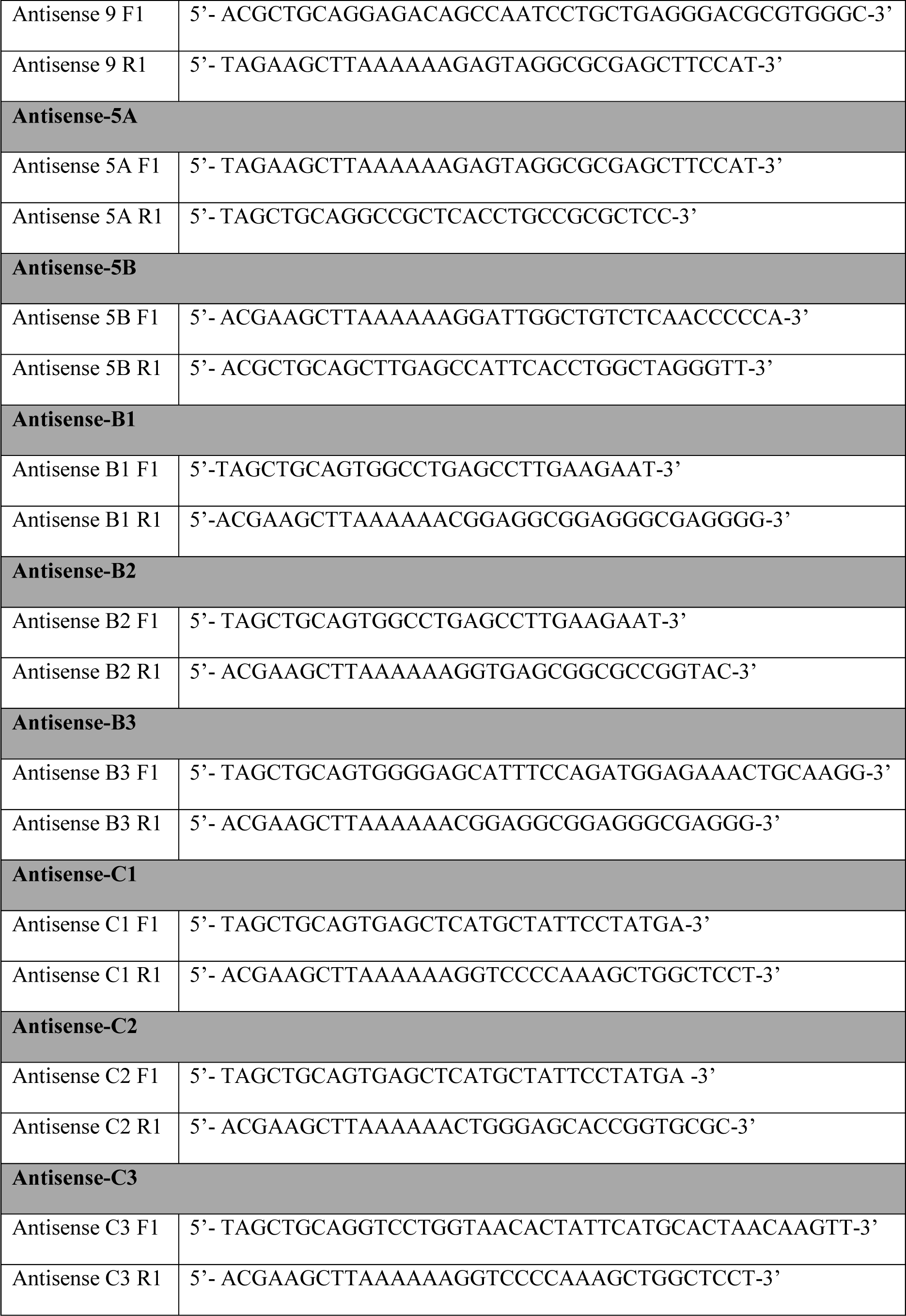

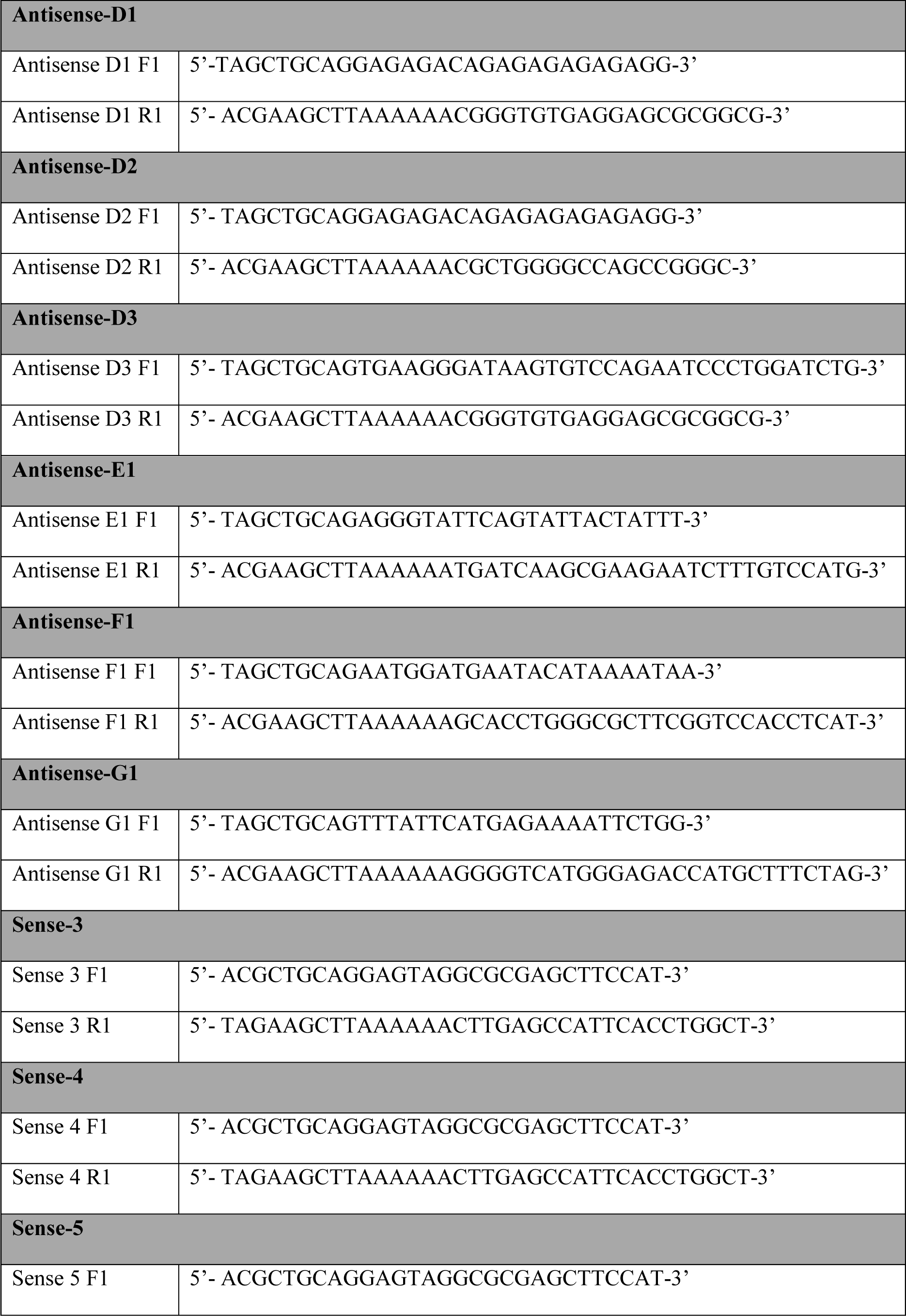

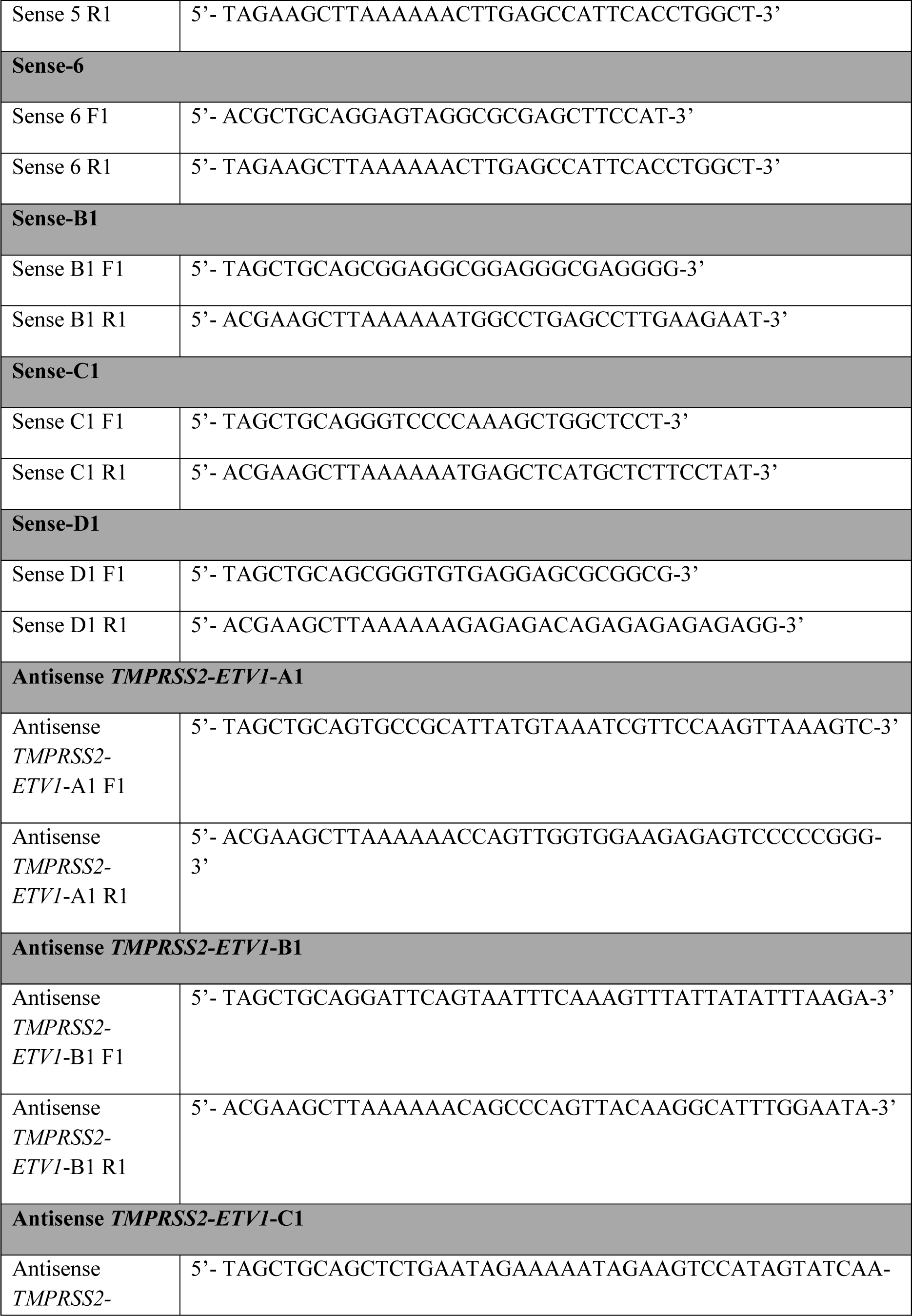

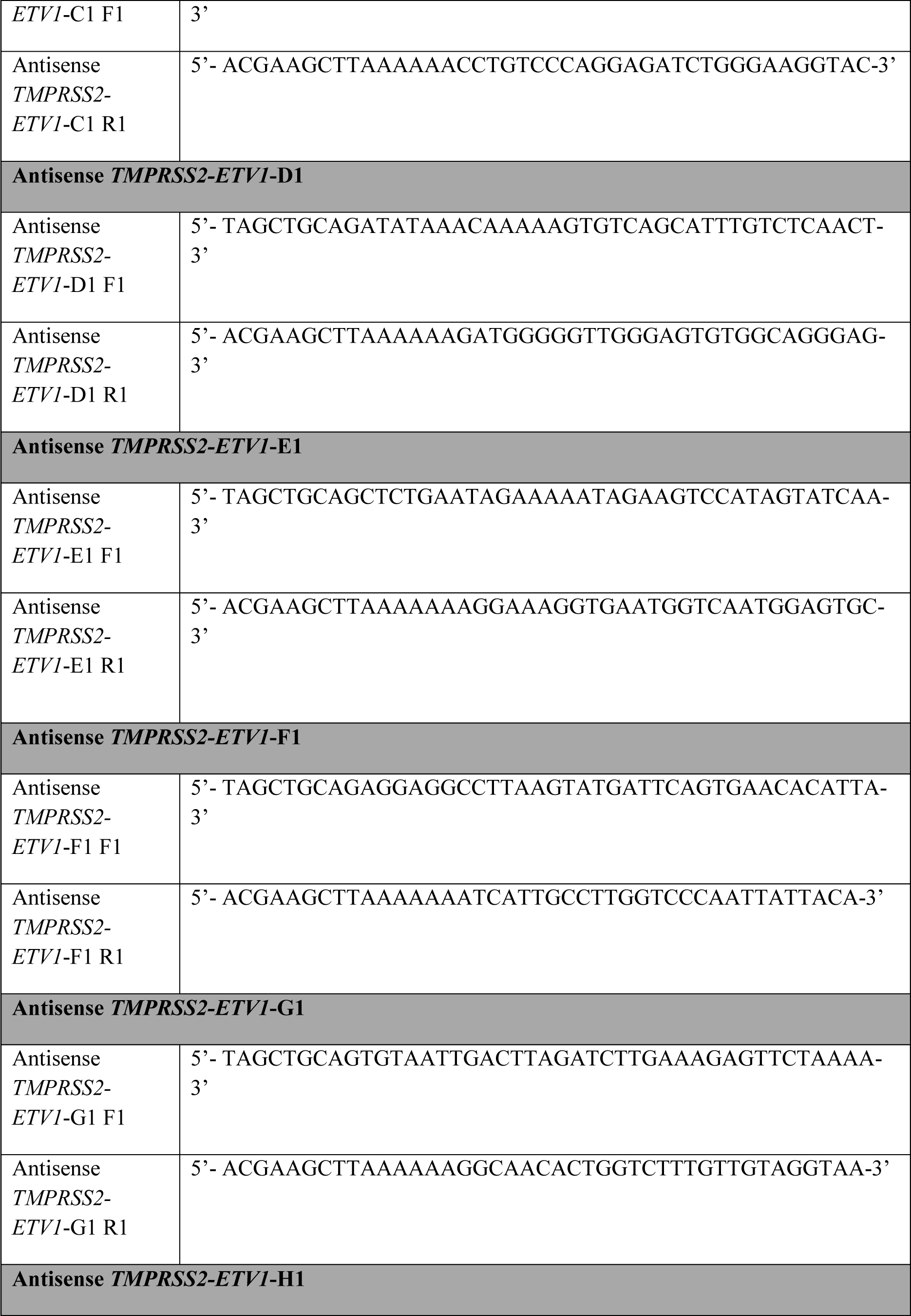

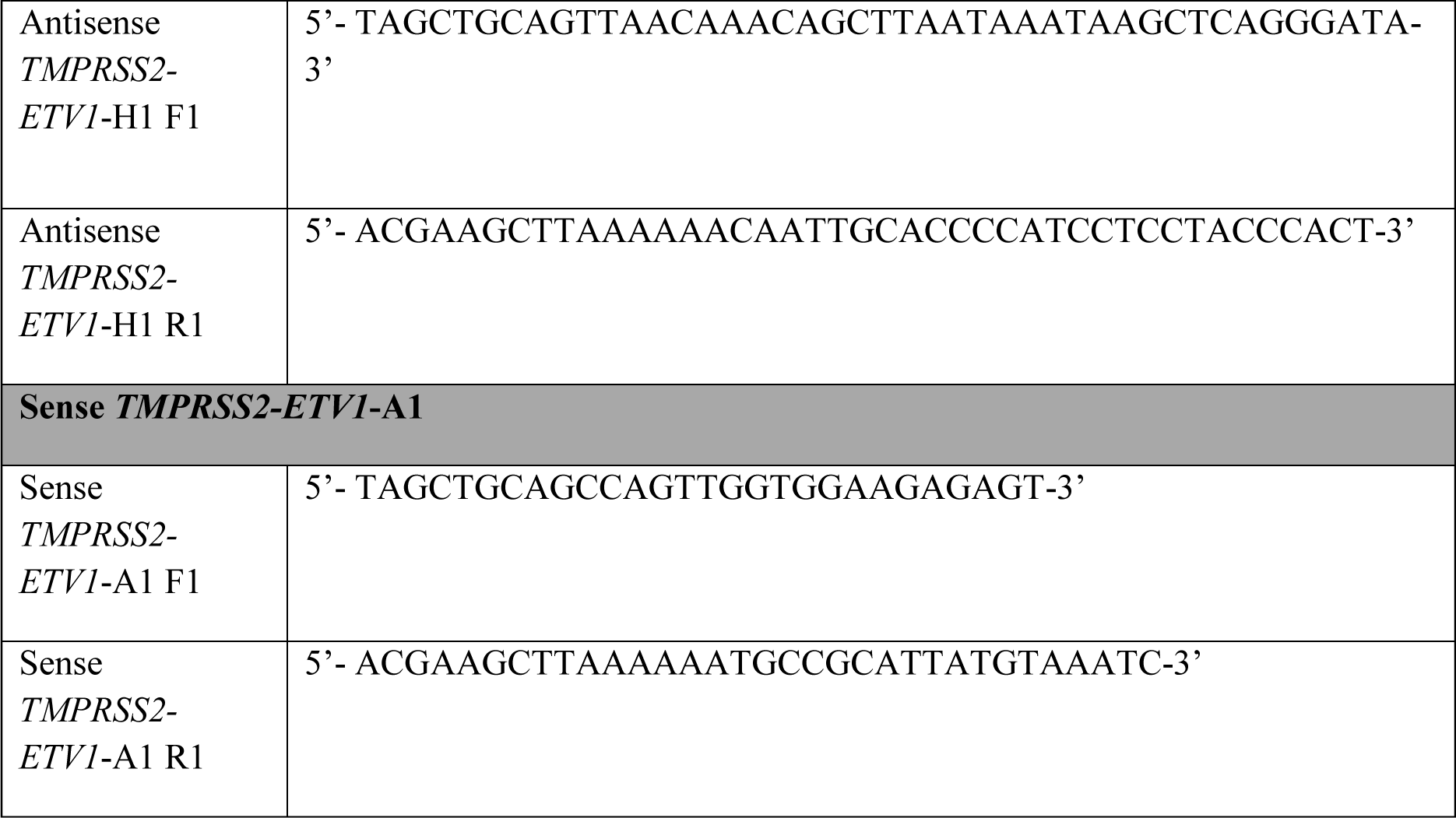

**PCR primers used for amplifying the identified TMPRSS2-ERG genomic DNA breakpoint:**

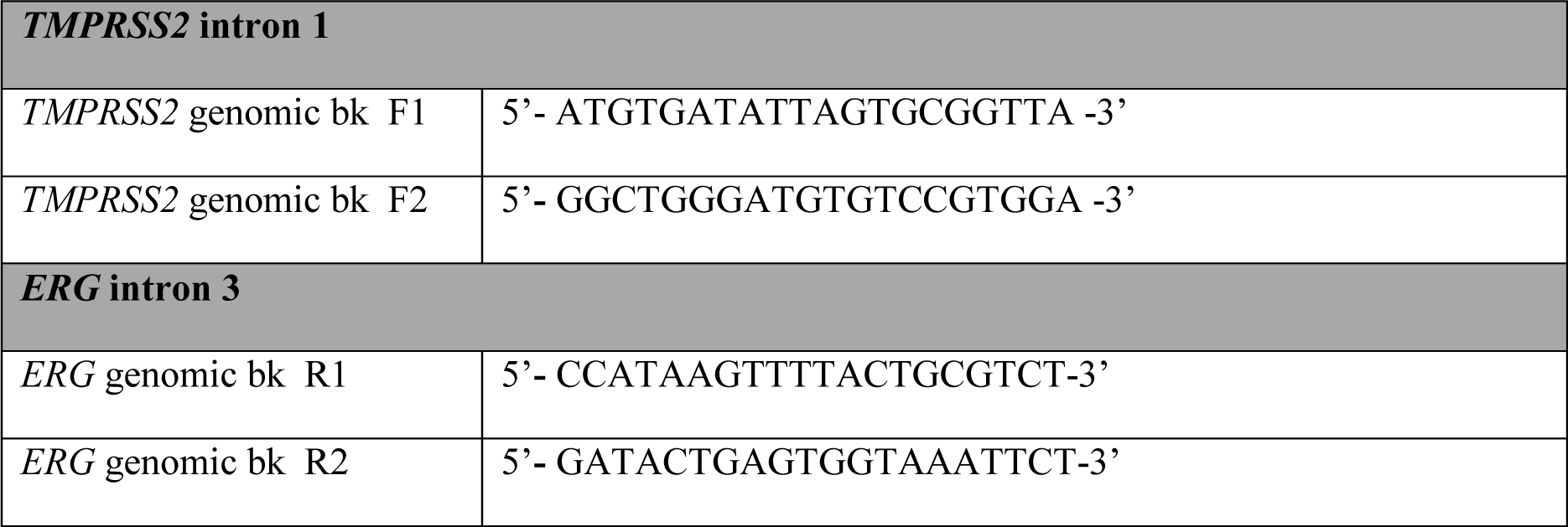

The rest of PCR primers used for genomic breakpoint analyses are not listed, but their locations are shown in Fig. S14A.

**PCR primers used for amplifying the identified TMPRSS2-ETV1 genomic DNA breakpoint:**

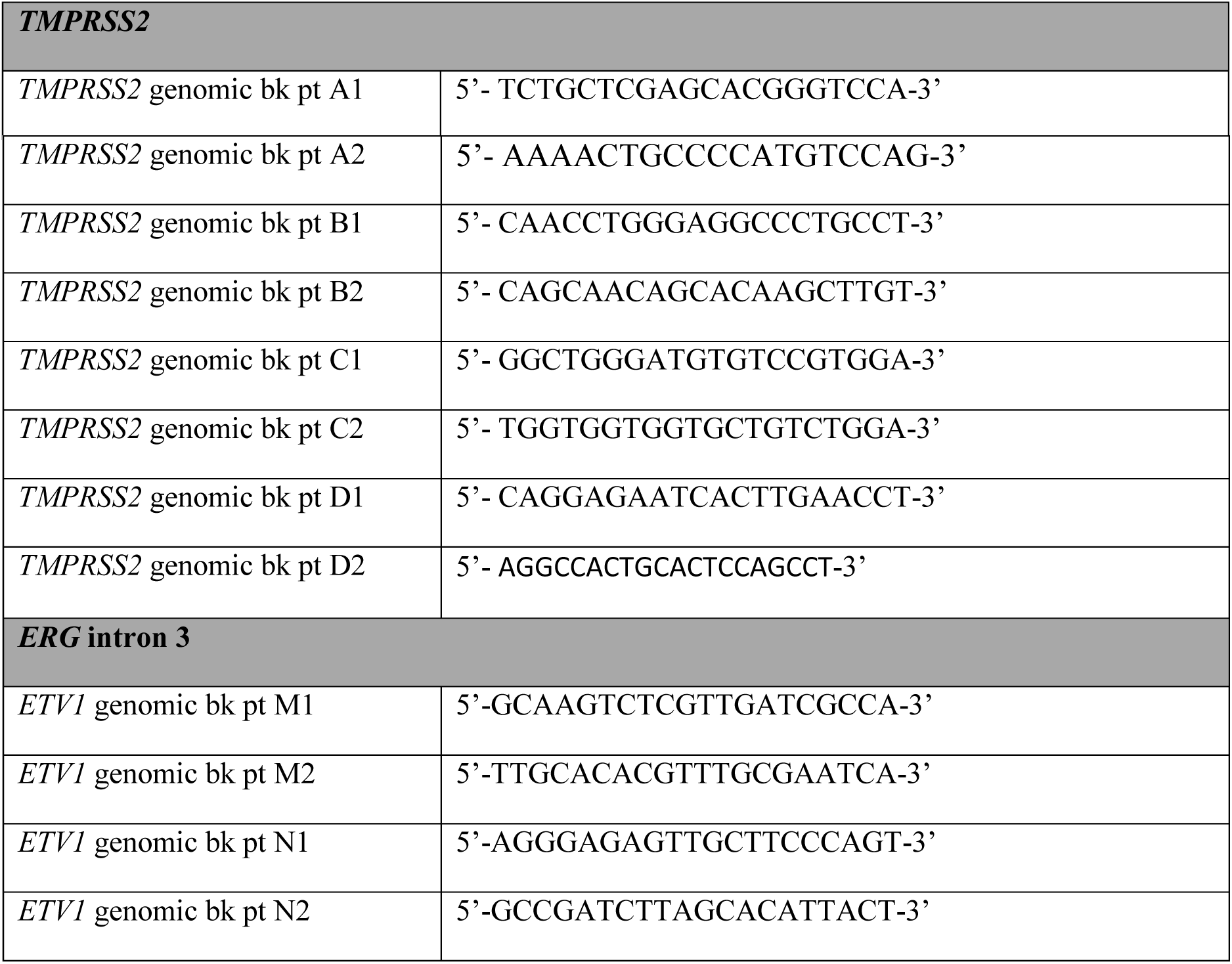

## References

Aoto, H., Miyake, Y., Nakamura, M., and Tajima, S. (1997). Genomic organization of the mouse AZ1 gene that encodes the protein localized to preacrosomes of spermatids. Genomics 40, 138-141.

Aoto, H., Tsuchida, J., Nishina, Y., Nishimune, Y., Asano, A., and Tajima, S. (1995). Isolation of a novel cDNA that encodes a protein localized to the pre-acrosome region of spermatids. Eur J Biochem 234, 8-15.

Bastus, N.C., Boyd, L.K., Mao, X., Stankiewicz, E., Kudahetti, S.C., Oliver, R.T., Berney, D.M., and Lu, Y.J. (2010). Androgen-induced TMPRSS2:ERG fusion in nonmalignant prostate epithelial cells. Cancer Res 70, 9544-9548.

Boyce, M.J., Baisley, K.J., Clark, E.V., and Warrington, S.J. (2004). Are published normal ranges of serum testosterone too high? Results of a cross-sectional survey of serum testosterone and luteinizing hormone in healthy men. in BJU Int, pp. 881-885.

Coll-Bastus, N., Mao, X., Young, B.D., Sheer, D., and Lu, Y.J. (2015). DNA replicationdependent induction of gene proximity by androgen. Hum Mol Genet 24, 963-971.

Crick, F. (1970). Central dogma of molecular biology. Nature 227, 561-563.

Fang, W., and Landweber, L.F. (2012). RNA-mediated genome rearrangement: hypotheses and evidence. Bioessays 35, 84-87.

Horoszewicz, J.S., Leong, S.S., Chu, T.M., Wajsman, Z.L., Friedman, M., Papsidero, L., Kim, U., Chai, L.S., Kakati, S., Arya, S.K., et al. (1980). The LNCaP cell line--a new model for studies on human prostatic carcinoma. Prog Clin Biol Res 37, 115-132.

Janz, S., Potter, M., and Rabkin, C.S. (2003). Lymphoma- and leukemia-associated chromosomal translocations in healthy individuals. Genes Chromosomes Cancer 36, 211-223.

Kannan, K., Wang, L., Wang, J., Ittmann, M.M., Li, W., and Yen, L. (2011). Recurrent chimeric RNAs enriched in human prostate cancer identified by deep sequencing. Proc Natl Acad Sci U S A 108, 9172-9177.

Kimsey, I., and Al-Hashimi, H.M. (2014). Increasing occurrences and functional roles for high energy purine-pyrimidine base-pairs in nucleic acids. Curr Opin Struct Biol 24, 72-80.

Langabeer, S.E., Walker, H., Rogers, J.R., Burnett, A.K., Wheatley, K., Swirsky, D., Goldstone, A.H., and Linch, D.C. (1997). Incidence of AML1/ETO fusion transcripts in patients entered into the MRC AML trials. MRC Adult Leukaemia Working Party. Br J Haematol 99, 925-928.

Li, H., Wang, J., Mor, G., and Sklar, J. (2008). A neoplastic gene fusion mimics trans-splicing of RNAs in normal human cells. Science 321, 1357-1361.

Lieber, M.R. (2010). The mechanism of double-strand DNA break repair by the nonhomologous DNA end-joining pathway. Annu Rev Biochem 79, 181-211.

Lin, C., Yang, L., Tanasa, B., Hutt, K., Ju, B.G., Ohgi, K., Zhang, J., Rose, D.W., Fu, X.D., Glass, C.K., et al. (2009). Nuclear receptor-induced chromosomal proximity and DNA breaks underlie specific translocations in cancer. Cell 139, 1069-1083.

Mani, R.S., Tomlins, S.A., Callahan, K., Ghosh, A., Nyati, M.K., Varambally, S., Palanisamy, N., and Chinnaiyan, A.M. (2009). Induced chromosomal proximity and gene fusions in prostate cancer. Science 326, 1230.

Mitelman, F., Johansson, B., and Mertens, F. (2007). The impact of translocations and gene fusions on cancer causation. Nat Rev Cancer 7, 233-245.

Nowacki, M., Vijayan, V., Zhou, Y., Schotanus, K., Doak, T.G., and Landweber, L.F. (2008). RNA-mediated epigenetic programming of a genome-rearrangement pathway. Nature 451, 153-158.

Perner, S., Demichelis, F., Beroukhim, R., Schmidt, F.H., Mosquera, J.M., Setlur, S., Tchinda, J., Tomlins, S.A., Hofer, M.D., Pienta, K.G., et al. (2006). TMPRSS2:ERG fusion-associated deletions provide insight into the heterogeneity of prostate cancer. Cancer Res 66, 8337-8341.

Rowley, J.D., and Blumenthal, T. (2008). Medicine. The cart before the horse. Science 321, 1302-1304.

Rubin, M.A., Maher, C.A., and Chinnaiyan, A.M. (2011). Common gene rearrangements in prostate cancer. J Clin Oncol 29, 3659-3668.

Rudiger, N.S., Gregersen, N., and Kielland-Brandt, M.C. (1995). One short well conserved region of Alu-sequences is involved in human gene rearrangements and has homology with prokaryotic chi. Nucleic Acids Res 23, 256-260.

Tomlins, S.A., Rhodes, D.R., Perner, S., Dhanasekaran, S.M., Mehra, R., Sun, X.W., Varambally, S., Cao, X., Tchinda, J., Kuefer, R., et al. (2005). Recurrent fusion of TMPRSS2 and ETS transcription factor genes in prostate cancer. Science 310, 644-648.

Watson, J.D., and Crick, F.H. (1953). Genetical implications of the structure of deoxyribonucleic acid. Nature 171, 964-967.

Weier, C., Haffner, M.C., Mosbruger, T., Esopi, D.M., Hicks, J., Zheng, Q., Fedor, H., Isaacs, W.B., De Marzo, A.M., Nelson, W.G., et al. (2013). Nucleotide resolution analysis of TMPRSS2 and ERG rearrangements in prostate cancer. J Pathol 230, 174-183.

Zaphiropoulos, P.G. (2011). Trans-splicing in Higher Eukaryotes: Implications for Cancer Development? Front Genet 2, 92.

Zhang, F., Carvalho, C.M., and Lupski, J.R. (2009). Complex human chromosomal and genomic rearrangements. Trends Genet 25, 298-307.

## References for Supplementary Information

Mertz, K.D., Setlur, S.R., Dhanasekaran, S.M., Demichelis, F., Perner, S., Tomlins, S., Tchinda, J., Laxman, B., Vessella, R.L., Beroukhim, R., et al. (2007). Molecular characterization of TMPRSS2-ERG gene fusion in the NCI-H660 prostate cancer cell line: a new perspective for an old model. Neoplasia 9, 200-206.

Rychlik, W., and Rhoads, R.E. (1989). A computer program for choosing optimal oligonucleotides for filter hybridization, sequencing and in vitro amplification of DNA. Nucleic Acids Res 17, 8543-8551.

Teles Alves, I., Hiltemann, S., Hartjes, T., van der Spek, P., Stubbs, A., Trapman, J., and Jenster, G. (2013). Gene fusions by chromothripsis of chromosome 5q in the VCaP prostate cancer cell line. Hum Genet 132, 709-713.

Weier, C., Haffner, M.C., Mosbruger, T., Esopi, D.M., Hicks, J., Zheng, Q., Fedor, H., Isaacs, W.B., De Marzo, A.M., Nelson, W.G., et al. Nucleotide resolution analysis of TMPRSS2 and ERG rearrangements in prostate cancer. J Pathol 230, 174-183.

